# Dynamic Modeling of Antibody Repertoire Reshaping in Response to Viral Infections

**DOI:** 10.1101/2024.05.28.596342

**Authors:** Zhaobin Xu, Junxiao Xu, Hongmei Zhang, Jian Song, Dongqing Wei, Qiangcheng Zeng

## Abstract

For many years, researchers have emphasized the production of high-affinity specific antibodies by hosts during viral infections. However, this has made it challenging for immunologists to systematically evaluate the initiation mechanisms of humoral immunity in specific immune responses. Employing mathematical modeling, we have systematically investigated the dynamic changes of the entire antibody atlas in response to exogenous antigenic stimuli, including viral infections. Our study reveals that the host’s antibody atlas is reshaped during viral infection, not through the proliferation of individual antibody types, but rather through the proliferation of antibody pools with strong binding activity. Moreover, we observe a contraction in pools of antibodies with low binding activity. We have identified the crucial role of self-antigens in maintaining antibody persistence, which can effectively explain the organism’s lifelong protection against pathogens that are less prone to mutation. Using this model, we further explore the mechanisms underlying original antigenic sin and elucidate the specific practical applications of this model. This research transcends the limitations of mere mathematical parameter fitting, as we endeavor to elucidate the complex humoral immune processes using physical mechanisms as a foundation. Our work contributes to a renewed understanding of the antibody elicitation process in specific immune responses.

## Main

Antibodies play a pivotal role in adaptive immunity. Investigating the interaction between antibodies and viruses is of paramount importance, as it contributes to a systemic understanding of the entire process of adaptive immune response activation. Furthermore, such research holds significant theoretical implications for the development of vaccines and antibody-based therapeutics [1–3].

Over the past decades, the dynamics of virus-host interactions have been extensively studied, leading to the emergence of numerous mathematical models [4–8]. While these models can effectively simulate individual immunological phenomena, they often fail to accurately describe the kinetic changes of antibodies following organismal infection. The primary issues stem from two aspects: firstly, nearly all models struggle to account for the long-term persistence of antibodies. Experimental evidence suggests that antibodies undergo a significant decline in the short term, but this decline is not sustained, as antibodies exhibit remarkable fluctuations over an extended period [9–10]. Secondly, almost all models consider the changes of individual antibodies rather than the overall changes in the antibody repertoire. With the vast diversity of B cells in the organism, viral infection is not merely a process of enhancing specific neutralizing antibodies but rather a reshaping of the entire antibody atlas [11–12].

Based on the shortcomings of our previous model [13] and fundamental principles of immunology, we propose a novel mathematical model to study virus-host interactions. This model incorporates antibodies with various binding affinities and considers the reshaping process of the entire antibody atlas during viral infection. Our mathematical model holds several significant implications:

Ⅰ: Successfully explains how the immune system selects antibodies with high binding activity. Ⅱ: Effectively simulates the dynamic changes of the antibody atlas following viral infection. Ⅲ: Reconceptualizes the long-term maintenance effect of IgG from a kinetic perspective. Ⅳ: Provides a coherent explanation for the formation of original antigenic sin. Ⅴ: Through parameter fitting, identifies differences in individual-specific immune dynamic parameters, which lead to variations in immune responses among individuals. Ⅵ: Suggests that antibodies with high binding activity are more effective in preventing secondary infections than antibodies with high binding strength, emphasizing the importance of prioritizing the discovery of antibodies with high binding activity in antibody screening. Ⅶ: Successfully explains the changes in IgM and IgG during the infection process, theoretically elucidating why many vaccines exhibit better efficacy upon secondary administration.

### Mathematical modeling of Virus-host interactions without considering antibody diversity

Molecular binding is a fundamental process in chemistry and biology, where two molecules interact and form a stable association (Molecular Binding, n.d.). The binding between antigens and antibodies is a crucial example of such molecular interactions, which is central to the immune system’s function. Understanding the principles and mechanisms governing antigen-antibody binding is essential for various applications, including antibody engineering, vaccine development, and disease diagnosis [14].

In antigen-antibody binding, the reaction is a reversible process, with an equilibrium constant (Kd) defined as the ratio of the dissociation rate constant (Koff) to the association rate constant (Kon) (Equilibrium Dissociation Constant (Kd), n.d.). Thermodynamically, the equilibrium constant Kd is related to the change in Gibbs free energy (ΔG) of the binding process through the equation ΔG = -RTln(Kd), where R is the gas constant and T is the absolute temperature [15]. This linear correlation between ln(Kd) and ΔG implies that the higher the binding energy (i.e., the lower the Kd value), the greater the proportion of bound complexes.

While the equilibrium constant Kd is often used to characterize antigen-antibody binding affinity, the association (Kon) and dissociation (Koff) rate constants are equally important in understanding the binding process. These kinetic parameters directly influence the time required to reach equilibrium, as demonstrated in Figure 1a. Two reactions with the same ΔG, and hence the same Kd, can have vastly different Kon and Koff values, resulting in significantly different times to reach equilibrium. Reactions with lower energy barriers for association and dissociation will reach equilibrium much faster than those with higher energy barriers.

**Figure 1a:**
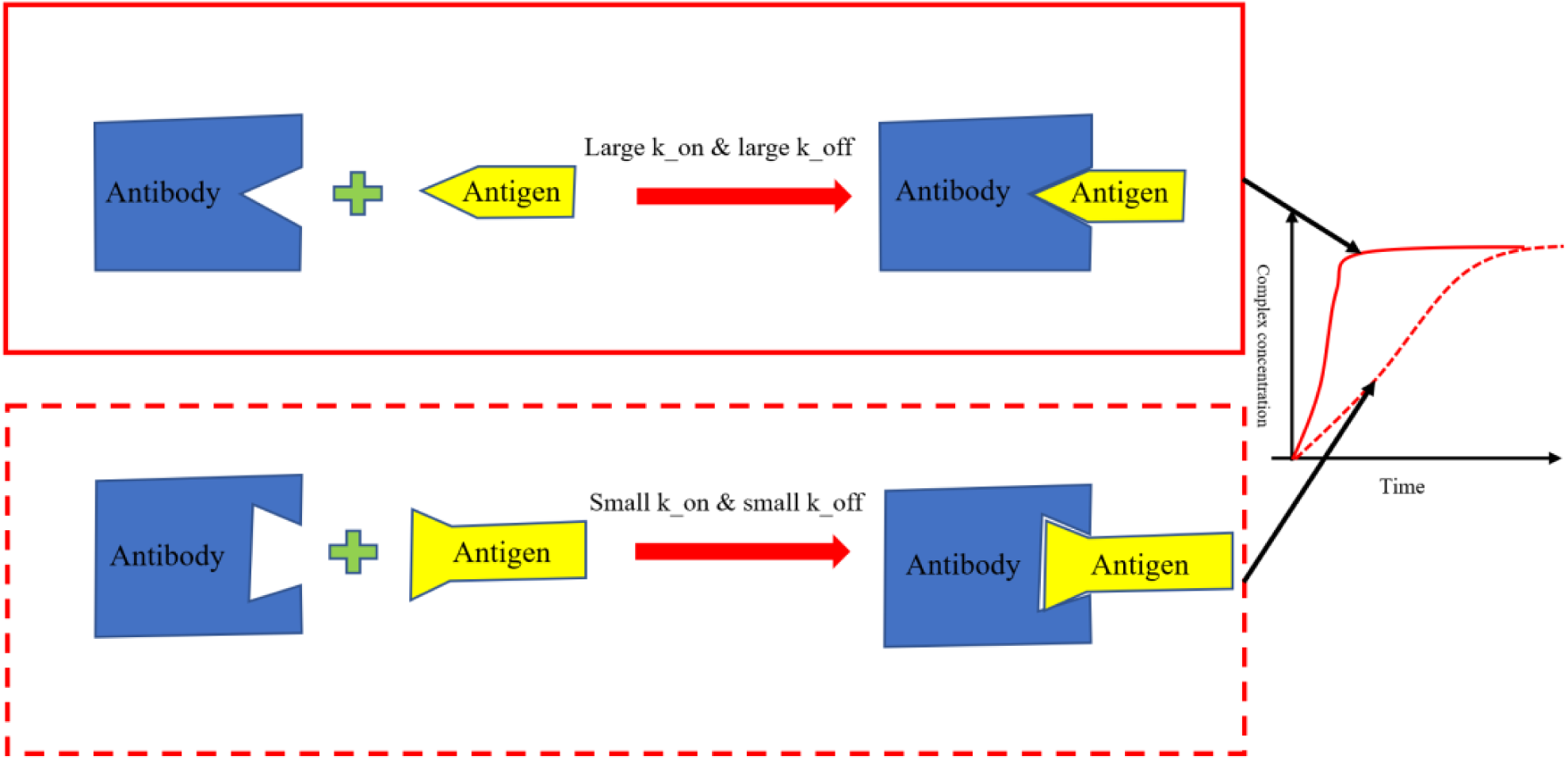
An illustration of antibody-antigen binding process. The above figure represents antibodies with relatively large forward binding coefficients and reverse dissociation coefficients, while the lower figure represents antibodies with relatively small forward binding coefficients and reverse dissociation coefficients. Both types of antibodies have the same binding affinity, possess the same equilibrium constant Kd, but the antibodies in the upper figure reach equilibrium more rapidly.

This concept is particularly relevant when simulating antigen-antibody interactions, as the binding process is inherently reversible. By explicitly incorporating Kon and Koff values, the binding dynamics can be more accurately modeled, providing deeper insights into the mechanisms governing antigen-antibody interactions.

Furthermore, the immune system’s preference for antibodies with specific binding kinetics has important implications. While traditional antibody screening often focuses on selecting the highest-affinity antibodies (i.e., those with the lowest Kd values), the immune system actually favors antibodies with faster association rates (high Kon) and faster dissociation rates (high Koff), even if their Kd values are similar. Such antibodies are more effective in clearing viruses and providing protection against re-infection, as they can rapidly bind to and dissociate from their targets.

The principles of molecular binding, including the reversible nature of antigen-antibody interactions and the importance of Kon, Koff, and Kd, are crucial for understanding and simulating the dynamics of these interactions. While Kd is a valuable metric for characterizing binding affinity, the kinetic parameters Kon and Koff play a pivotal role in determining the time required to reach equilibrium and the immune system’s preference for certain antibody characteristics. Incorporating these fundamental concepts into the analysis and design of antigen-antibody systems can lead to more effective and targeted approaches in areas such as antibody engineering, vaccine development, and disease diagnosis.

Before introducing the antibody atlas simulation, it is imperative to elucidate the kinetic characteristics of specific antibody types. This model is depicted in Figure 1b. Process 1 represents the virus self-replication, characterized by a proliferation coefficient (k1). Process 2 signifies the virus-antibody binding process, with a forward reaction coefficient (k2) and a reverse dissociation coefficient (k-2). Process 3 denotes the positive feedback effect of virus-antibody complexes on antibody regeneration, governed by a feedback coefficient (k3). Process 4 illustrates the in vivo clearance of virus-antibody complexes, governed by a reaction coefficient (k4).

**Figure 1b:**
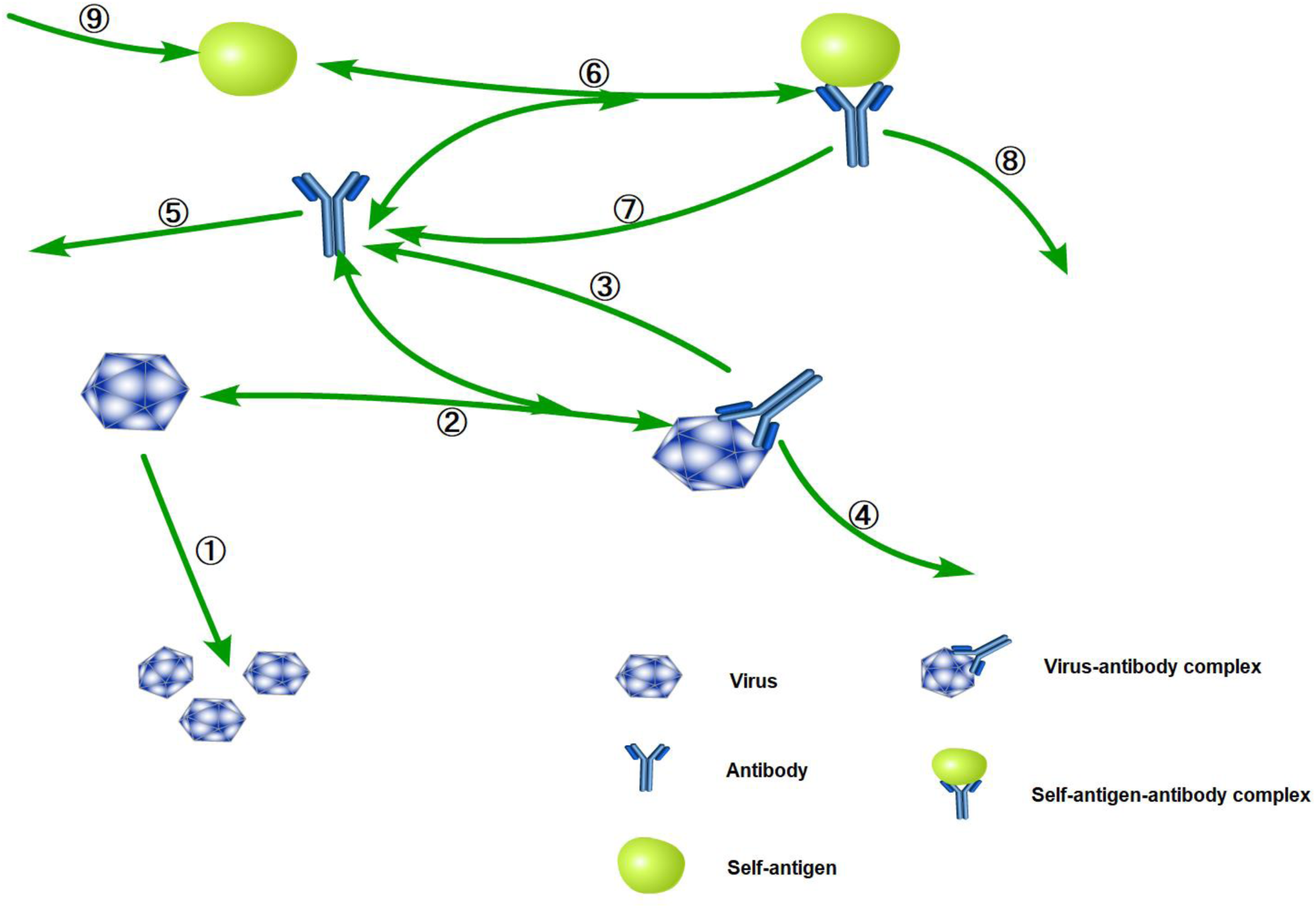
An illustration of antibody-virus interaction mechanism. Here, we utilized five components to represent the entire process, primarily including virus, antibody, self-antigen, virus-antibody complex, and self-antigen-antibody complex.

Process 5 symbolizes the self-decay of antibodies, governed by a reaction coefficient (k5). Reaction 6 delineates the binding process between self-antigenic substances and antibodies, characterized by a forward reaction coefficient (k6) and a reverse dissociation coefficient (k-6). Reaction 7 delineates the positive feedback effect of self-antigen-antibody complexes on antibody regeneration, governed by a feedback coefficient (k7). Reaction 8 elucidates the clearance process of self-antigen-antibody complexes, governed by a coefficient (k8). Reaction 9 delineates the supplementary action of self-antigens, with a supplementation rate (c1). The detailed physical mechanisms of each reaction are expounded in the supplementary materials.

It is worth noting the long-term sustaining effect of self-antigens on antibody levels. There is controversy regarding the role of self-antigens in the maintenance of memory B cells. While many studies indicate that the maintenance of memory B cells does not necessitate continuous stimulation by the primary antigen [16], it has been found that MHC-restricted B cell antigen presentation is crucial for memory cell maintenance [17–18]. Additionally, numerous studies have shown that memory B cells do not exhibit longer half-lives [19–20]. These findings collectively suggest that memory B cells require the proliferation of helper T cells to be sustained at a certain level, with self-antigens likely being the targets of antigen presentation.

Simulation results depicted in Figure 1c illustrate that in the presence of self-antigens, antibody levels within the body can be maintained at a stable level even after the complete clearance of the virus, with occasional fluctuations observed (blue dashed line). Conversely, as shown by the solid red line in Figure 1c, in the absence of self-antigens, antibody levels within the body undergo continuous decay (process 5), failing to be maintained above a threshold concentration and eventually diminishing until complete disappearance. Detailed information is provided as model 1.1 in supplementary materials. This observation clearly contradicts what is observed in reality. The relationship between the initial concentration of self-antigens and the kinetic coefficient of antibody binding is elaborated upon in the supplementary material.

**Figure 1c:**
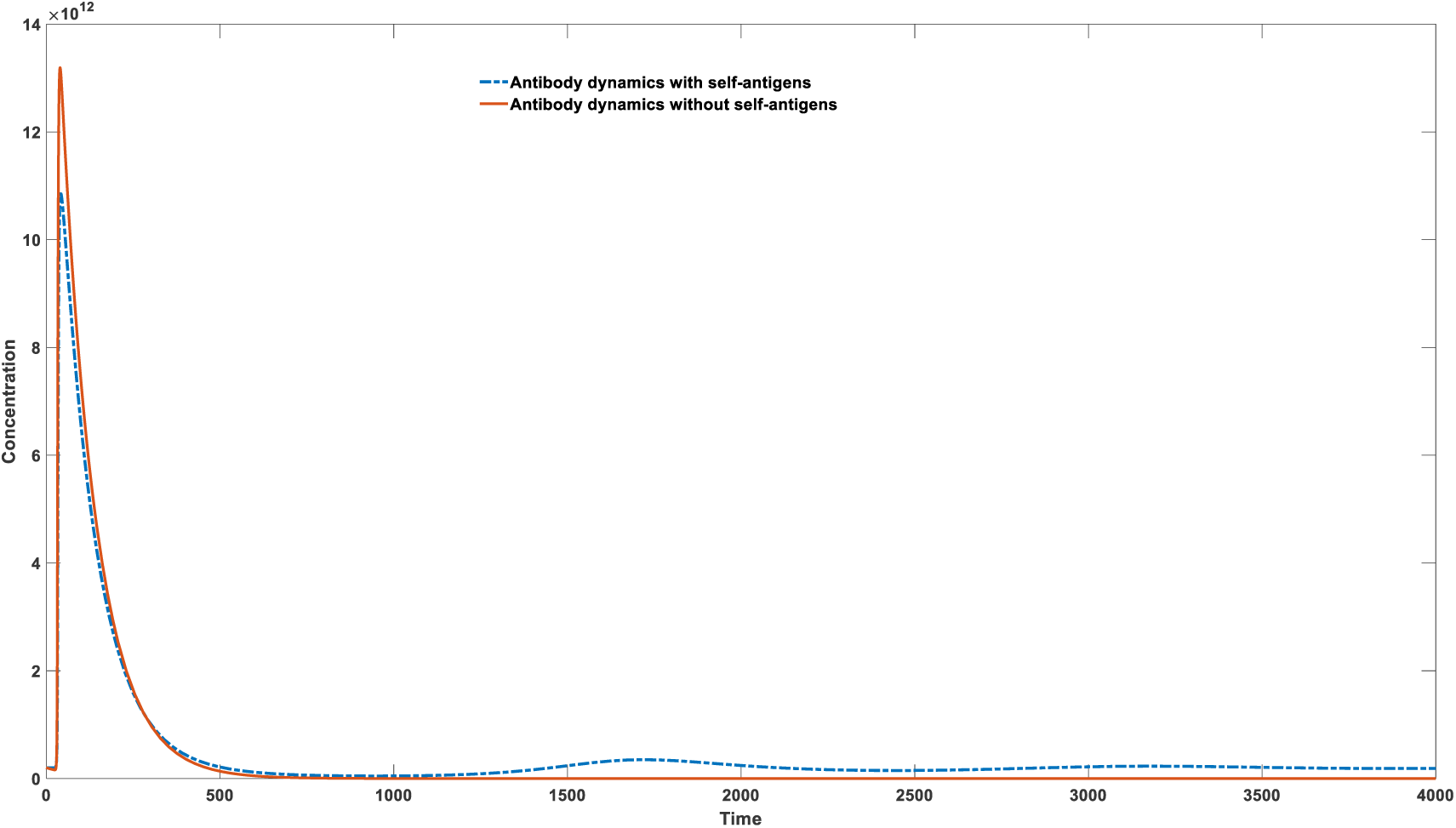
The critical roles of self-antigens in maintaining B-cell homeostasis. The blue dashed line represents the changes in antibodies in the presence of self-antigen, while the solid red line represents the changes in antibodies in the absence of self-antigen.

Upon viral infection, the immune system undergoes robust proliferation of specific antibodies with high affinity and binding strength. This process, orchestrated by the adaptive immune response, is crucial for combating viral pathogens effectively. When a virus invades the body, specialized immune cells, such as B lymphocytes, recognize viral antigens and undergo clonal expansion, generating a large pool of antigen-specific B cells [21–22]. These B cells differentiate into plasma cells, which are antibody-producing factories. Through a process called affinity maturation, these plasma cells refine their antibody production, selecting for variants with the highest affinity for the viral antigen. This results in the generation of antibodies that exhibit strong binding activity, effectively neutralizing the virus and preventing its further spread. Additionally, memory B cells are formed during this process, providing long-lasting immunity against future encounters with the same virus. Overall, the immune system’s ability to produce a vast quantity of specific antibodies with enhanced binding affinity plays a pivotal role in the host’s defense against viral infections.

We further investigated the mechanism by which specific immunity efficiently selects antibodies with high binding activity. Figure 1d illustrates this process. All four antibodies have the same initial concentration. Antibody 1 exhibits a strong forward binding coefficient (Kon = 10^-12^) and dissociation coefficient (Koff = 0.1), resulting in a Kd value of 10^11^. Antibody 2 has the same forward binding coefficient (Kon = 10^-12^) but a higher dissociation coefficient (Koff = 1), yielding a Kd value of 10^12^. Antibodies 3 and 4 have the same Kd value as antibody 1. However, antibody 3 has a Kon of 0.9*10^-12^ and a Koff of 0.9*0.1, while antibody 4 has a Kon of 0.8*10^-12^ and a Koff of 0.8*0.1.

**Figure 1d:**
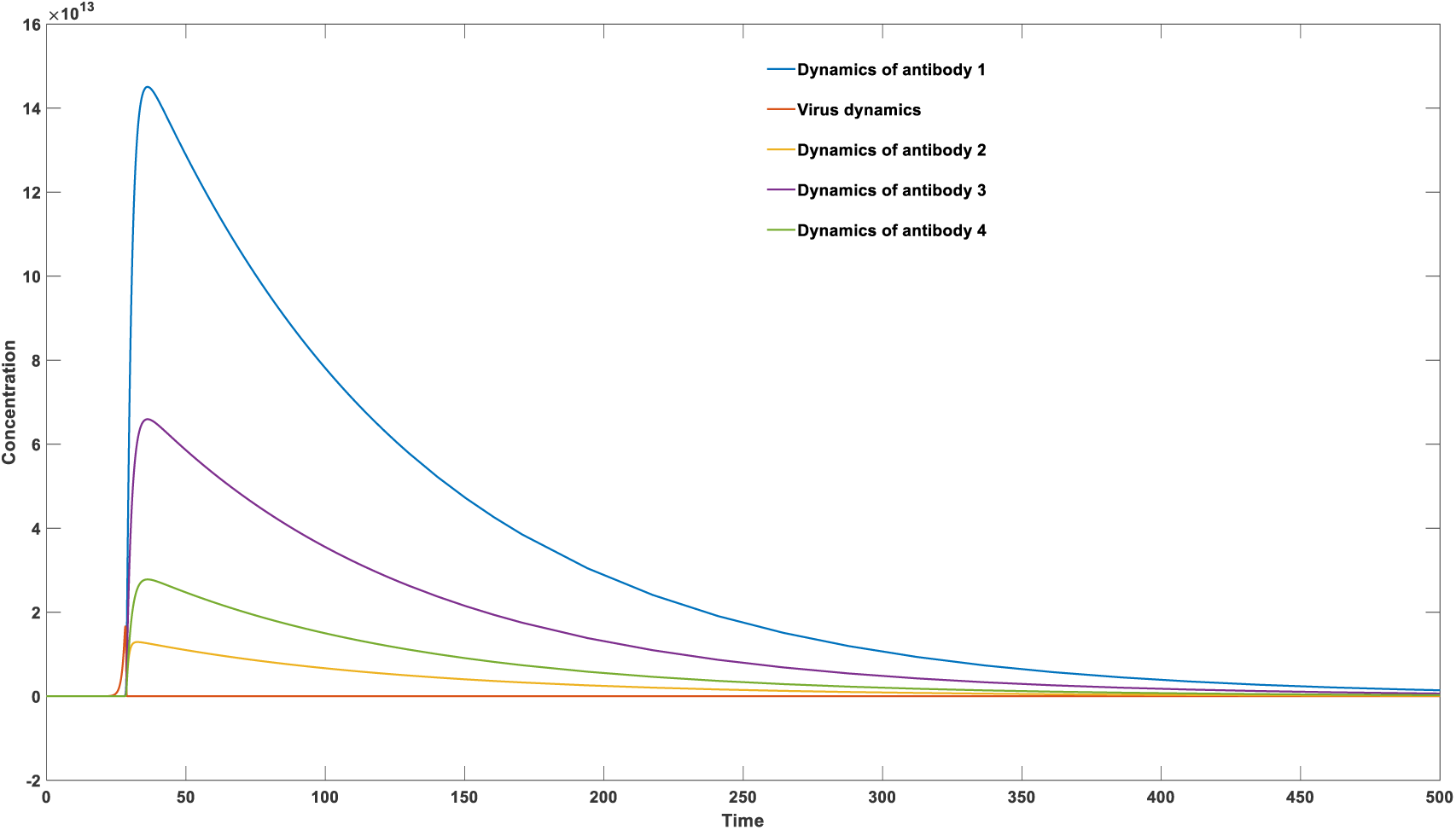
An illustration of how the immune system select strong binding antibodies. The solid red line represents the trend of the virus, the solid blue line represents the kinetic changes of Antibody 1, the solid yellow line represents the kinetic changes of Antibody 2, the solid purple line represents the kinetic changes of Antibody 3, and the solid green line represents the kinetic changes of Antibody 4. Antibody 1 has the same Kon value as Antibody 2, but Antibody 1 has a smaller Koff value than Antibody 2. Antibodies 1, 3, and 4 have the same Kd value, but the Kon values follow the pattern of Antibody 1 > Antibody 3 > Antibody 4.

From Figure 1d, it displays that antibodies with stronger binding affinity or more favorable equilibrium coefficients undergo greater proliferation. When equilibrium coefficients are equal, the organism tends to proliferate antibodies with stronger forward binding coefficients. Detailed information is provided as model 1.2 in supplementary materials.

The manifestation of symptoms during viral infections often exhibits considerable variability, ranging from asymptomatic cases to mild and severe presentations [23–24]. This diversity in symptomatology poses challenges for disease management and epidemiological control efforts. Understanding the underlying factors contributing to this variability is crucial for effective prevention and treatment strategies. Several factors contribute to the differential expression of symptoms during viral infections, including viral load, host immune response, genetic predisposition, and comorbidities. Variation in viral load influences the severity of symptoms, with higher viral loads typically associated with more severe manifestations [25–26]. Additionally, the host immune response plays a critical role, with differences in immune competence influencing the ability to control viral replication and mitigate symptoms. Genetic factors, such as variations in host cell receptors and immune-related genes, contribute to individual susceptibility and response to viral infections [27–29]. Moreover, pre-existing health conditions and comorbidities can exacerbate symptoms and increase the risk of severe disease outcomes [30–31]. Understanding the interplay between these factors is essential for elucidating the mechanisms underlying symptom variability and developing targeted interventions to mitigate disease severity and transmission.

In our model, we quantitatively define the severity of symptoms using the concentration of antigen-antibody complexes, as the manifestation of symptoms, including the severity of fever mechanisms, is primarily associated with the concentration of antigen-antibody complexes. Additionally, the antibody-dependent cellular cytotoxicity (ADCC) effect is directly correlated with the concentration of antigen-antibody complexes [32–33]. Elevated levels of either high viral load or high antibody levels do not directly exacerbate the severity of symptoms. Figure 1e compares the changes in virus concentration under different antibody treatments. The peak virus concentration (represented by the blue dashed line in the lower graph) was significantly lower when using antibodies with a higher Kon value than when using antibodies with a lower Kon value (represented by the blue solid line in the lower graph). Additionally, the concentration of virus-antibody complexes (represented by the red dashed line in the lower graph) was also significantly lower when using antibodies with a higher Kon value compared to those with a lower Kon value (represented by the red solid line in the lower graph). This suggests that antibodies with a faster binding rate can lead to milder infection symptoms. At the same time, the highest excitation concentration for antibodies with a higher Kon value (represented by the red dashed line in the upper graph) was also lower than that for antibodies with a lower Kon value (represented by the blue solid line in the upper graph). Detailed information is provided as model 1.3 in supplementary materials.

**Figure 1e:**
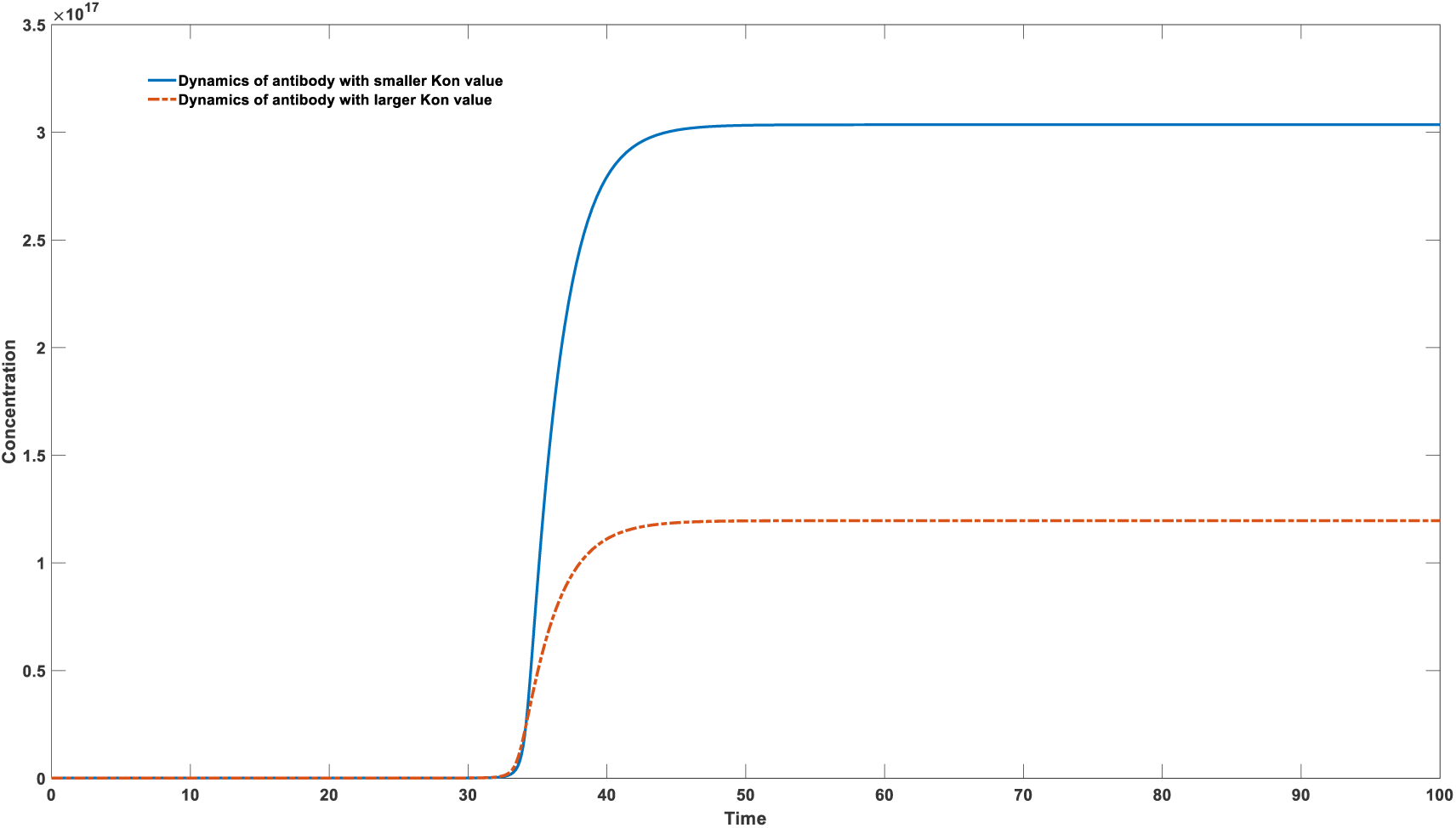
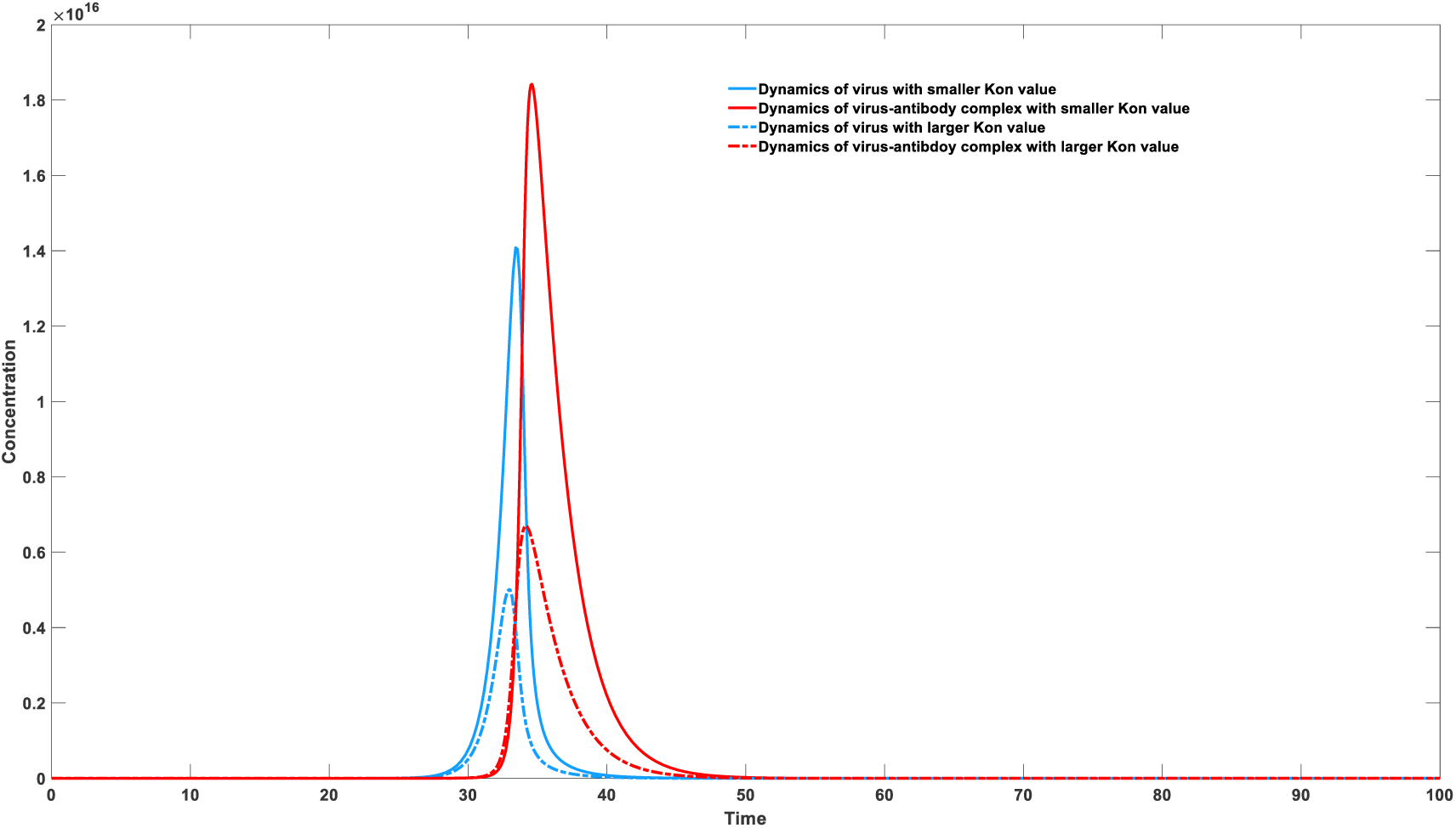
The severity of symptom can be reflected in the concentration of antibody-virus complex. The figure above illustrates the proliferation of antibodies with different kinetic characteristics in response to the same viral infection. The solid blue line represents the proliferation of antibodies with smaller Kon coefficients, while the dashed red line represents the proliferation of antibodies with larger Kon coefficients. The lower figure corresponds to the dynamic changes of the virus and virus-antibody complexes. The solid blue line denotes the viral dynamics in the presence of antibodies with smaller Kon coefficients, whereas the dashed blue line represents the viral dynamics in the presence of antibodies with larger Kon coefficients. Similarly, the solid red line represents the dynamics of virus-antibody complexes in the presence of antibodies with smaller Kon coefficients, while the dashed red line represents the dynamics of virus-antibody complexes in the presence of antibodies with larger Kon coefficients.

### Mathematical simulation of virus-host interactions considering antibody atlas mapping

Host antibodies are not a singular entity but rather comprise a vast array of antibodies constituting the antibody repertoire. This diversity arises due to several factors contributing to the formation of antibodies with distinct specificities within the host. Firstly, the genetic recombination and somatic hypermutation processes during B cell development generate a wide range of antibody sequences, leading to the production of antibodies recognizing diverse antigens [34]. Secondly, exposure to a plethora of antigens throughout an individual’s lifetime triggers the activation and clonal expansion of various B cell clones, each producing antibodies targeting specific antigens encountered [35]. Additionally, the process of affinity maturation further enhances antibody diversity by selecting for B cells producing antibodies with higher affinity towards specific antigens through iterative rounds of selection and mutation. Moreover, cross-reactivity, where antibodies can recognize structurally similar but distinct antigens, contributes to the overall diversity of the antibody repertoire [36]. Collectively, these mechanisms underpin the formation of a highly diverse antibody repertoire within the host, enabling effective immune responses against a wide range of pathogens and antigens.

Therefore, when studying antibodies, it is advisable not to simulate specific types of antibodies but rather to model the characteristics of the entire population, including the dynamic changes of antibodies with different binding affinities. For host antibody libraries that have not been exposed to specific viral infections, we assume that the distribution of antibodies follows a normal distribution pattern. Here, we consider that both the forward binding coefficient ln(Kon) and the reverse dissociation coefficient ln(Koff) follow a normal distribution. This assumption is based on the belief that the energy barrier heights ΔG for forward binding and reverse dissociation also follow a normal distribution pattern. In the supplementary material, based on binding experiments of macromolecules [37–38], we plotted the distributions of ln(Kon) and ln(Koff) for different antibodies binding to antigens, which preliminarily indicate a close fit to normal distribution. Therefore, for simulation purposes, we selected a normal distribution for ln(Kon) with parameters N(-15.5, 0.5) and for ln(Koff) with parameters N(1.5, 0.8), with specific units and thresholds detailed in the supplementary material. Figures 2a and 2b illustrate the characteristics of these distributions.

**Figure 2a:**
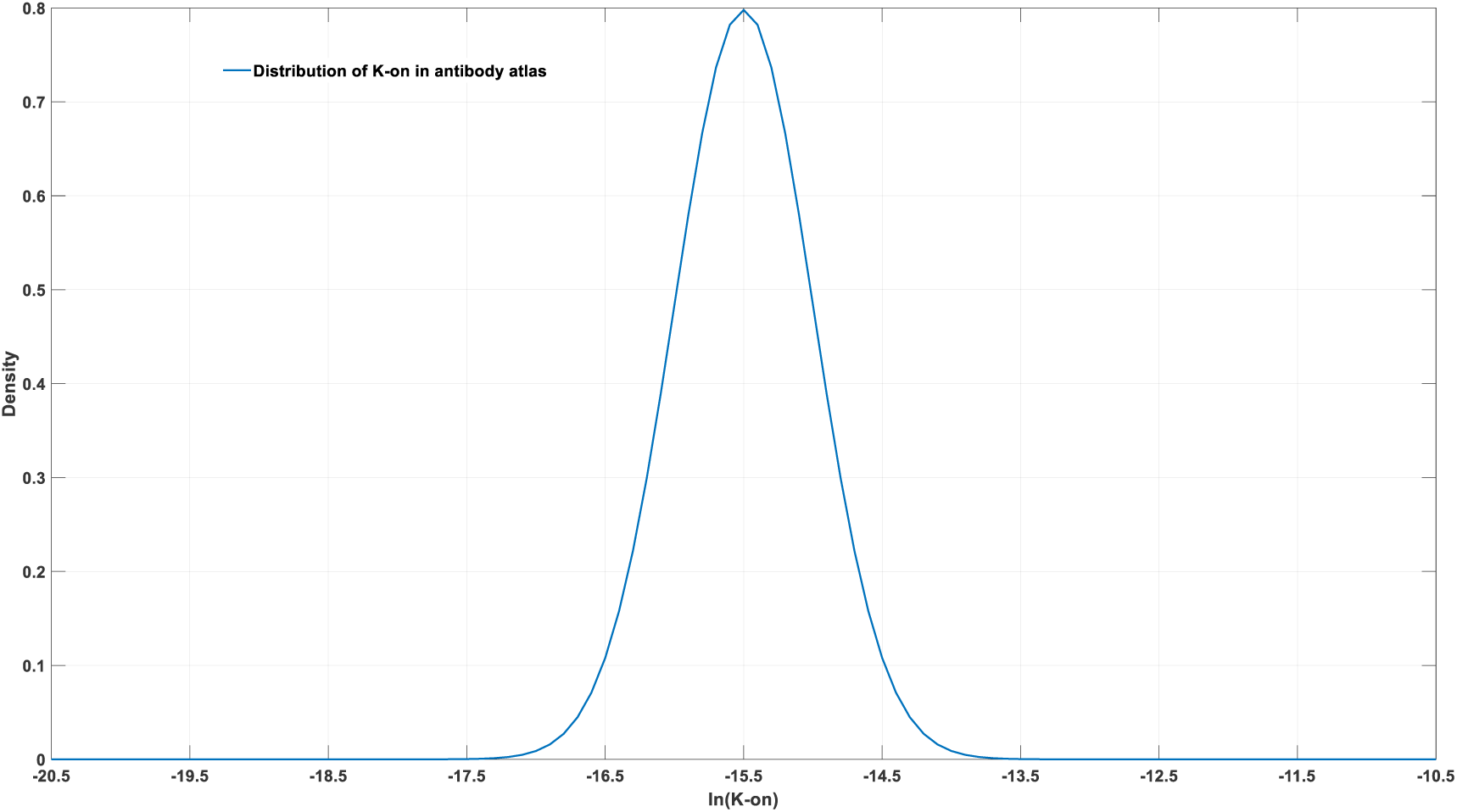
Distribution of Kon values in B-cell antibody repertoire. We discretize ln(Kon) into 10 parts, and the probability for each part can be obtained using the cumulative normal distribution function.

**Figure 2b:**
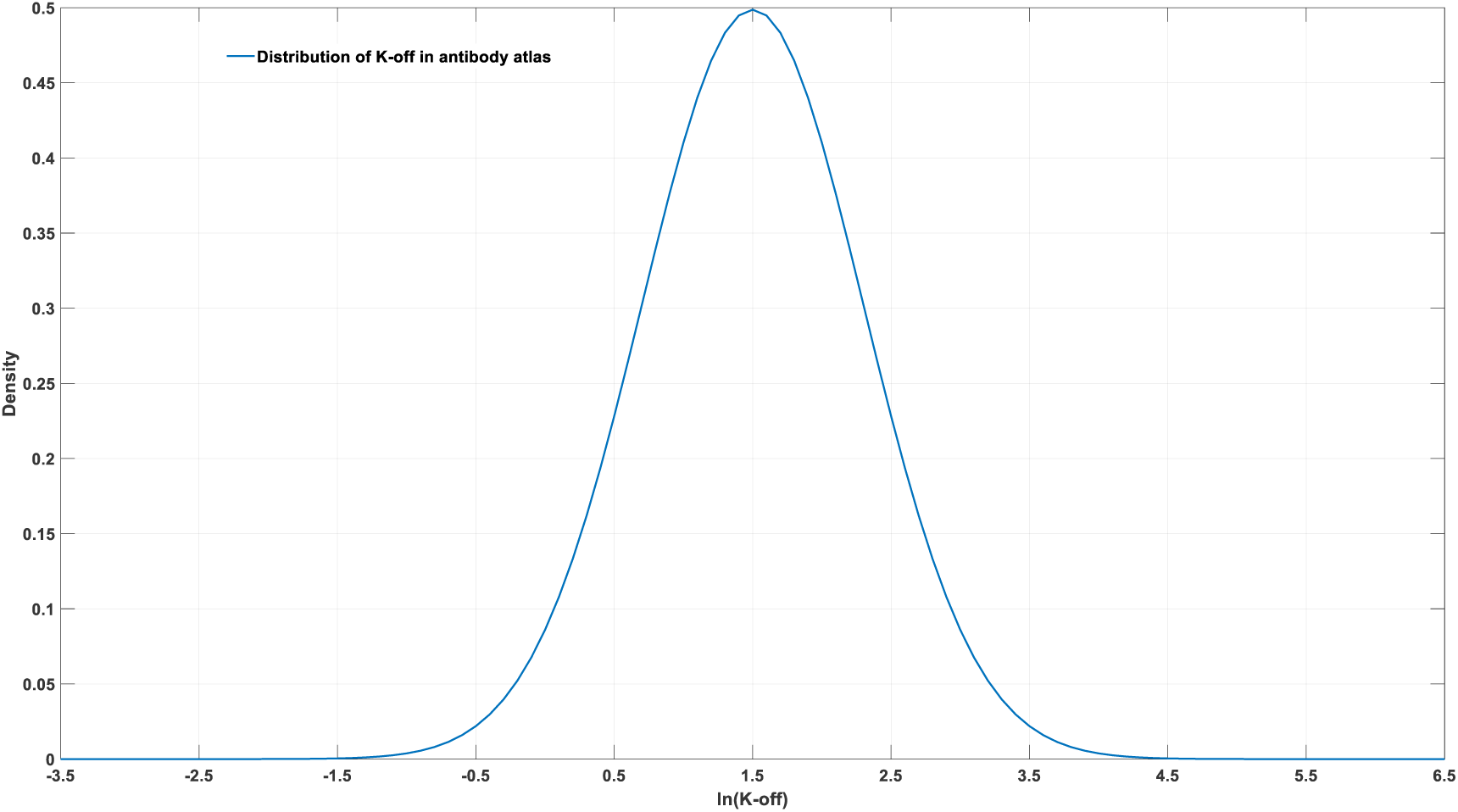
Distribution of Koff values in B-cell antibody repertoire. We discretize ln(Koff) into 10 parts, and the probability for each part can be obtained using the cumulative normal distribution function.

To efficiently conduct simulations, we discretized ln(Kon) into 10 intervals ranging from -20.5 to -10.5 with intervals of 1. Similarly, ln(Koff) was discretized into 10 intervals, as depicted in Figure 2b. Thus, these combinations result in 100 different antibodies, each with varying Kd values. The ln(Kd) values range from 8 to 26 with intervals of 1, totaling 18 different ln(Kd) values, and their distribution is shown in Figure 2c. This completes the construction of a simple antibody library consisting of 100 antibodies, each with different initial concentrations and combinations of Kon and Koff. A finer discretization would generate more antibody combinations, approaching real-world scenarios, but would increase computational costs and simulation time. Therefore, in this article, we only considered a 10*10 combination type. From these distributions, we observe that in the antibody atlas of unexposed hosts, antibodies with both high and low binding strengths have low initial concentrations, while antibodies with intermediate binding strengths dominate.

**Figure 2c:**
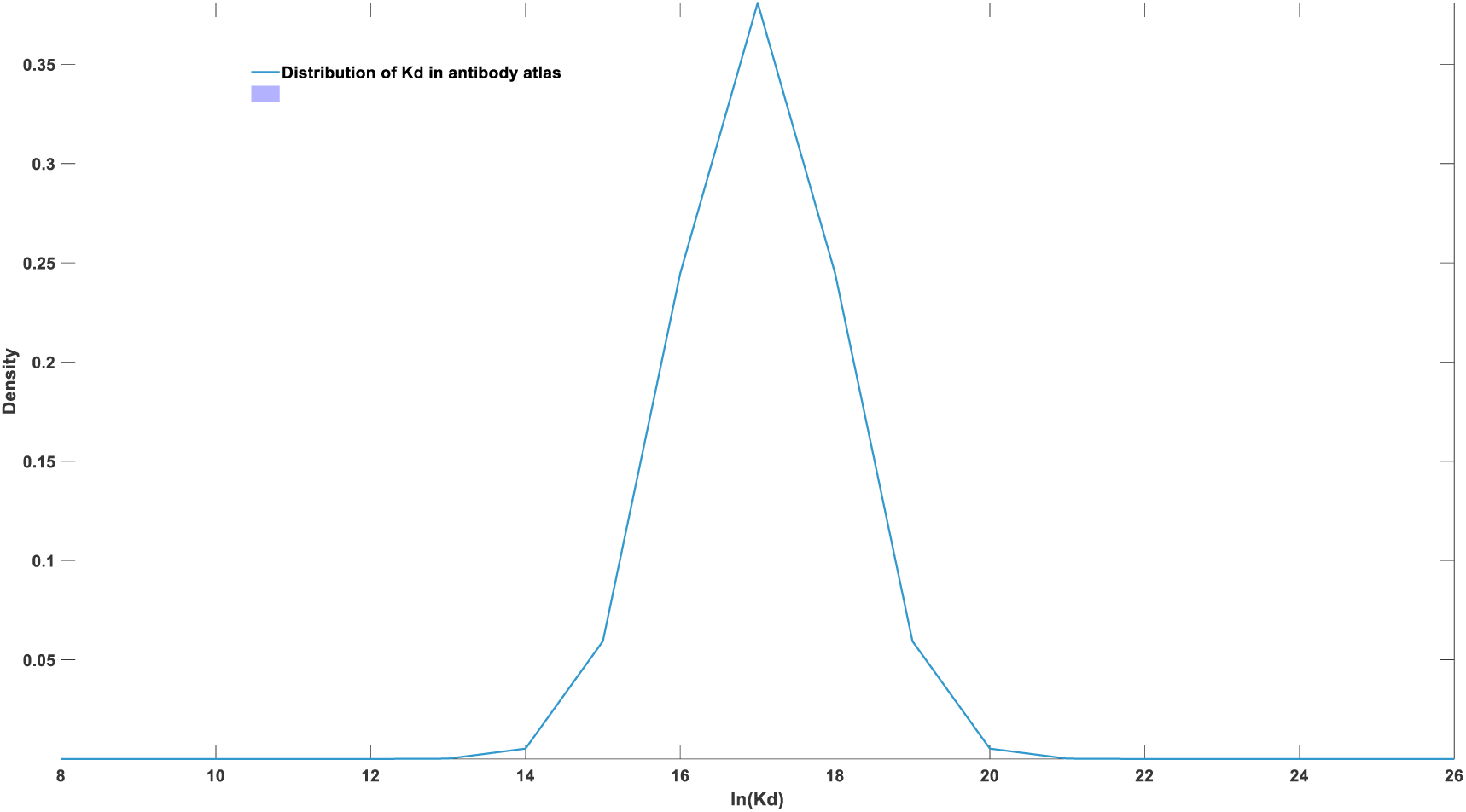
Distribution of Kd values in B-cell antibody repertoire. Based on the discretized probabilities from Figure 2a and 2b, we can obtain the density distribution of 100 antibodies within the range of 8-26.

We further investigated the role of endogenous antigenic substances in maintaining the host antibody library. The specific model is outlined in the Methods section. Considering the presence of endogenous antigenic substances, the distribution of host antibodies can persistently maintain a stable state, as depicted in the upper graph of Figure 2d. Otherwise, the antibody library would experience atrophy, leading to a decay in all antibody levels, as illustrated in the lower graph of Figure 2d. Detailed information is provided as model 2.1 in supplementary materials.

**Figure 2d:**
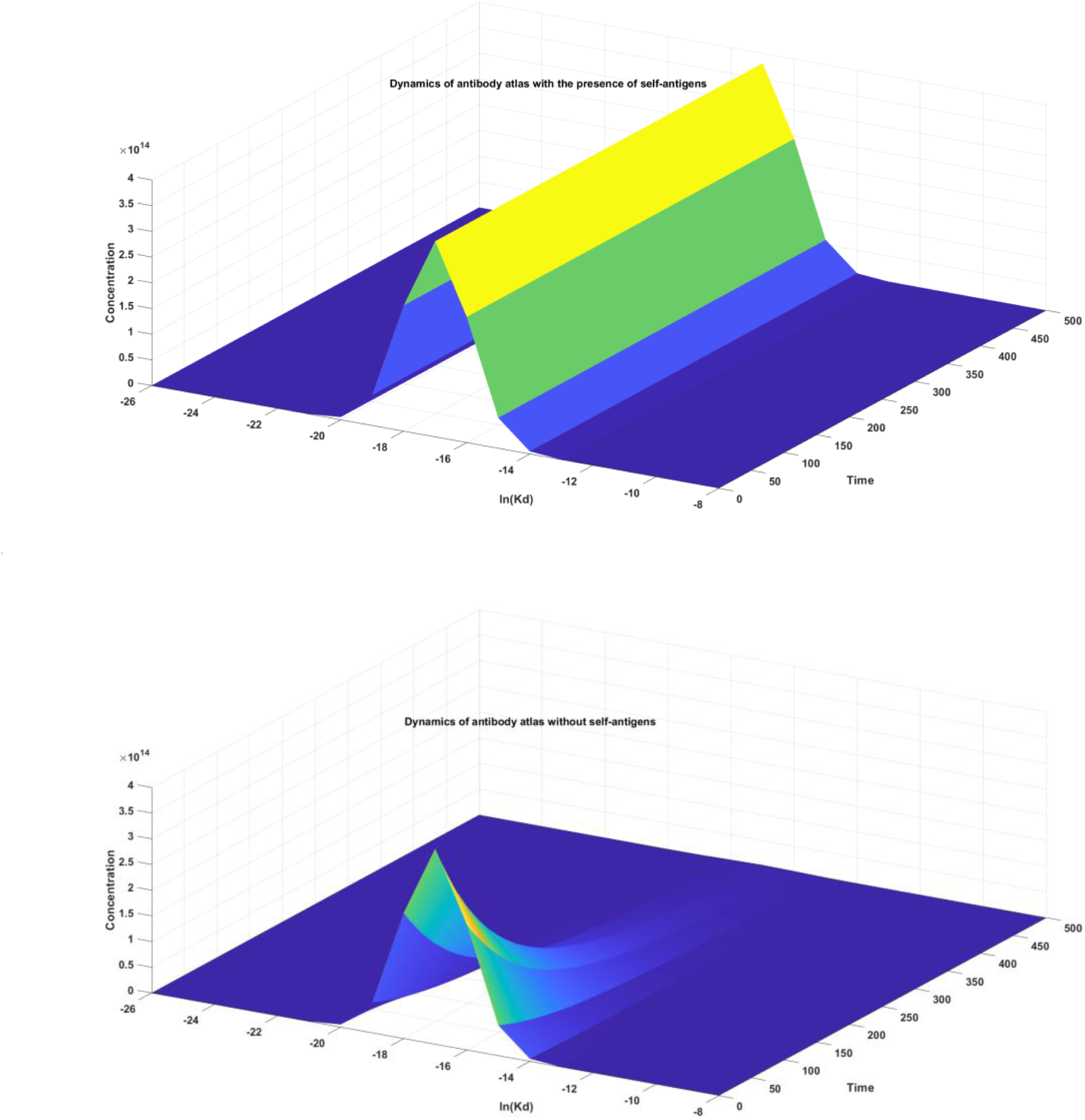
The homeostasis atlas of B-cells with the help of self-antigens. The upper figure represents the changes in the antibody repertoire in the presence of self-antigen, which can be maintained at a stable level. The lower figure represents the decay of the antibody repertoire in the absence of self-antigen.

**Figure 2e:**
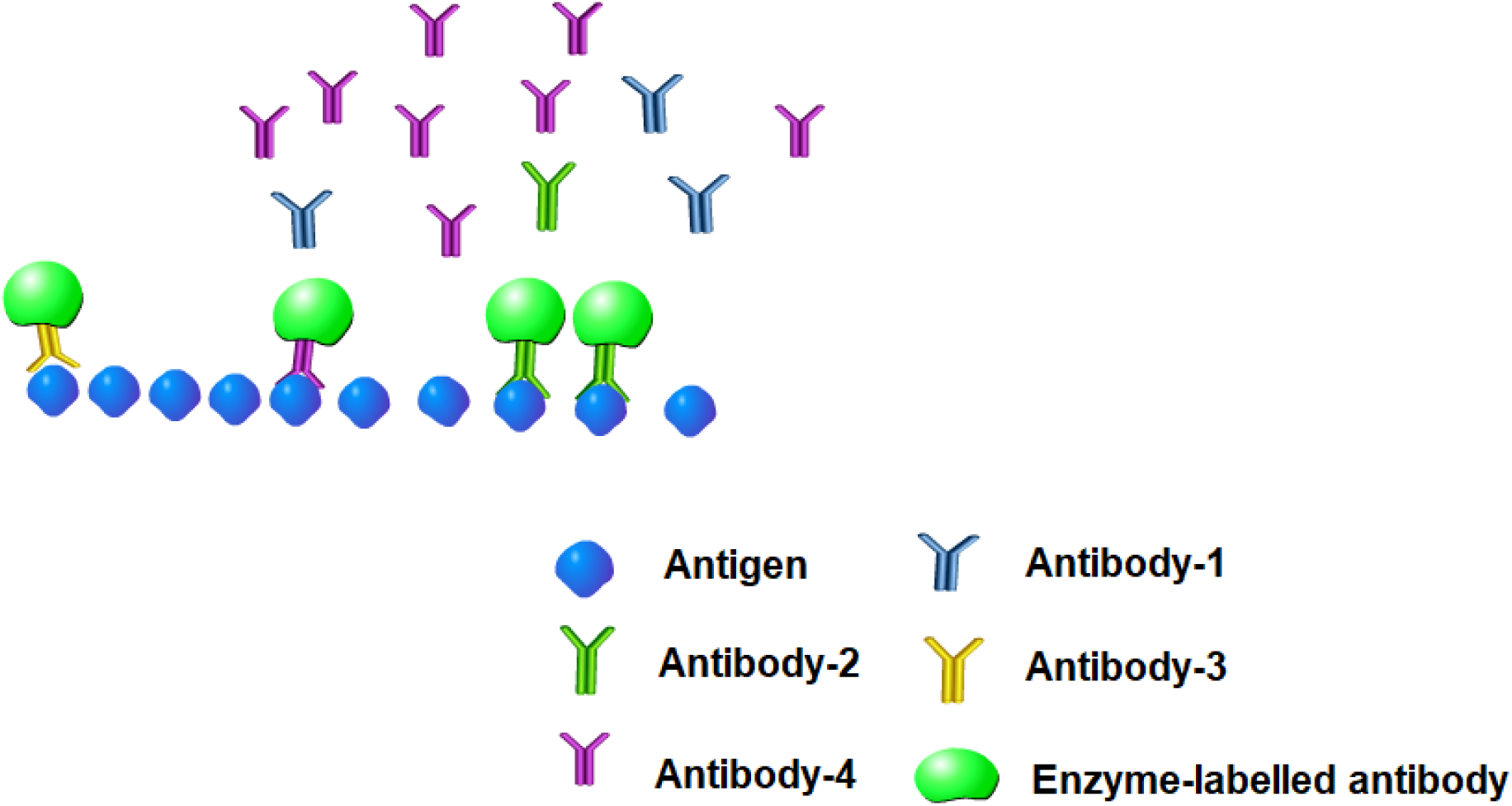
The mechanism of ELISA. As shown in the figure, there are four different types of antibodies, designated as antibody 1, antibody 2, antibody 3, and antibody 4. Antibody 1 exhibits very weak binding affinity to the antigen, as indicated by the absence of any binding between three antibody 1 molecules and the antigen in the figure. Antibody 2 demonstrates relatively strong binding affinity to the antigen, with two out of three antibody 2 molecules binding to the antigen. Although antibody 3 exhibits extremely strong binding affinity, its concentration is very low, with only one antibody 3 molecule present, but fully bound to the antigen. Antibody 4 shows relatively weak binding affinity, but it is present at a high concentration, with one out of nine antibody 4 molecules binding to the antigen. Upon addition of the enzyme-labeled antibody, the fluorescence intensity displays four fluorescent complexes. This does not represent the concentration of a specific antibody, but rather provides a comprehensive reflection of the overall antibody concentration and binding levels within the entire antibody repertoire.

Enzyme-Linked Immunosorbent Assay (ELISA) is a crucial experimental technique for measuring antibody levels [39–40]. Its principle relies on the specific binding between antigens and antibodies, which is detected using enzyme-conjugated secondary antibodies. The process typically involves several key steps. Antigens are immobilized onto a solid surface, such as a microplate, through physical adsorption or chemical binding. Then, the surface is blocked to prevent nonspecific binding of antibodies. The sample containing the antibodies of interest is added to the microplate wells. If present, the antibodies will bind to the immobilized antigens through antigen-antibody interactions. After washing away unbound components, a secondary antibody conjugated with an enzyme, such as horseradish peroxidase (HRP), is added. This secondary antibody recognizes and binds to the primary antibody, forming a complex. Following another washing step to remove excess secondary antibodies, a substrate specific to the enzyme is added. The enzyme catalyzes a reaction with the substrate, producing a detectable signal, such as a color change or fluorescence. The intensity of the signal is proportional to the amount of bound enzyme, which in turn reflects the quantity of primary antibody present in the sample. ELISA offers a sensitive and specific method for detecting and quantifying antibodies, making it indispensable in various research and diagnostic applications. It is important to emphasize that ELISA experiments do not directly reflect the concentration levels of specific antibodies; rather, they reflect the concentration of antibody complexes formed by all antibodies capable of binding. Many misconceptions often equate ELISA results with corresponding antibody concentrations, which is highly erroneous. To mitigate nonspecific binding, bovine serum is often used to pretreat samples, aiming to remove nonspecifically bound antibodies [41]. Sample serum pretreatment can significantly affect experimental results, as detailed simulations and explanations are provided in the supplementary materials. Nevertheless, ELISA cannot directly determine the concentration levels of a specific antibody after serum pretreatment. ELISA results represent the concentration of all antigen-antibody complexes in the mixture. Because ELISA testing involves the use of antigenic substances and multiple mixed antibodies, knowing the concentration and binding kinetics of individual antibodies enables the calculation of the final concentration of all antibody-antigen complexes, which constitutes the ultimate ELISA result. This feature is one of the characteristics of our model, where ELISA results at different times are calculated based on changes in the composition of the antibody library at different periods, thereby rendering them more reliable and physiologically meaningful.

### The evolutionary trend of antibody repertoires after virus infection

According to the mathematical model covering different types of antibodies, we simulated the compositional changes of the antibody library following viral infection. These changes are illustrated separately in three figures. Figure 3a depicts the variation of ln(Kon), showing a significant rightward shift after infection, indicating a notable proliferation of antibodies with higher forward binding coefficients in the antibody library. The dynamic characteristics of this change can be observed more clearly in Extended Video 1. Figure 3b represents the change in ln(Koff), demonstrating a significant leftward shift after infection, suggesting a notable proliferation of antibodies with slower dissociation rates in the antibody library. The dynamic characteristics of this change can be observed more clearly in Extended Video 2. Figure 3c illustrates the variation of ln(Kd), showing a significant rightward shift after infection, indicating a notable proliferation of antibodies with overall stronger binding affinity in the antibody library. The dynamic characteristics of this change can be observed more clearly in Extended Video 3. The specific trends of each antibody are displayed in Extended Figure 1. Detailed information is provided as model 2.2 in supplementary materials.

**Figure 3a:**
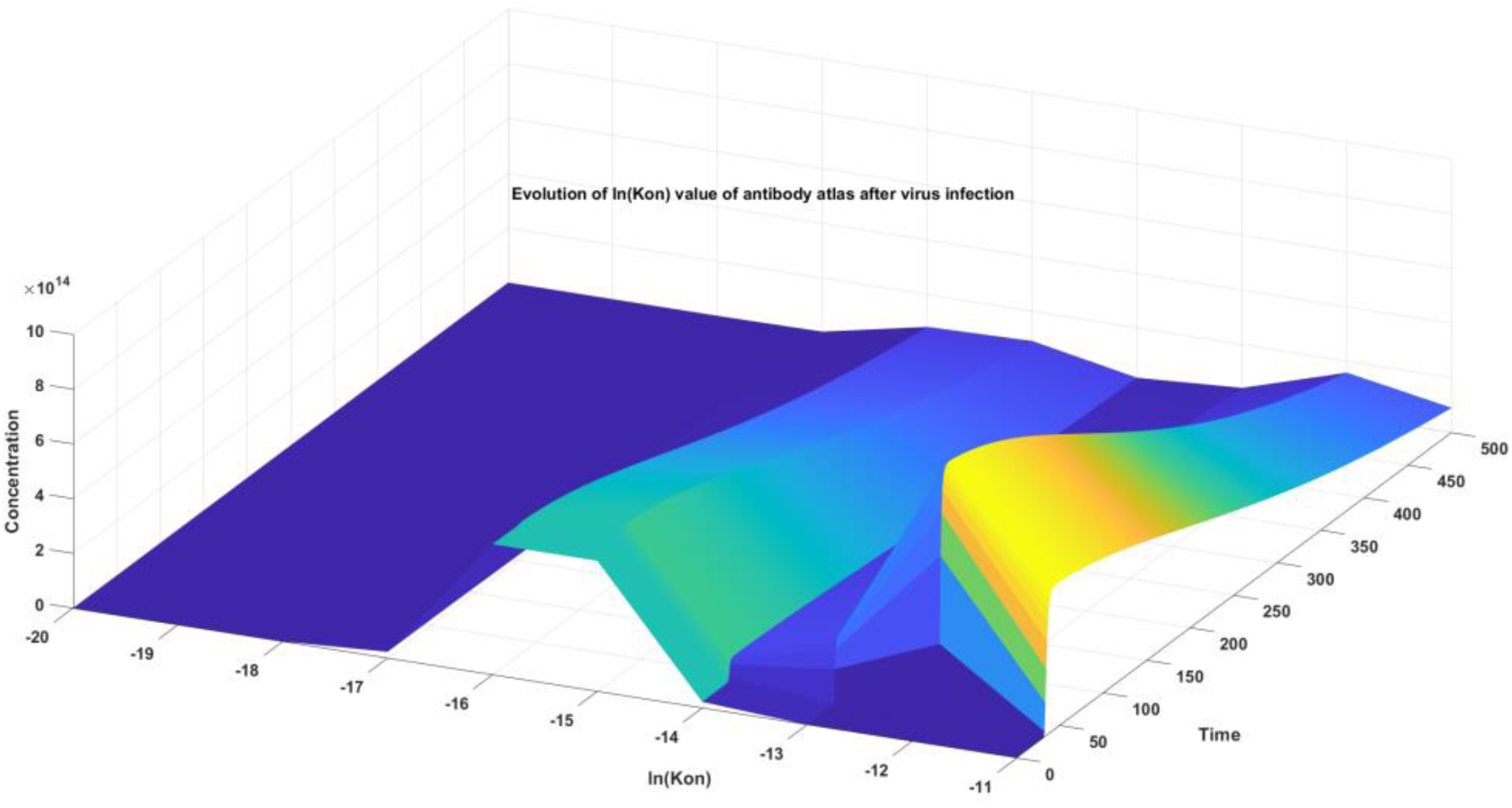
The evolutionary trend of antibody Kon value after virus infection. From the graph, it can be observed that the initial ln(Kon) values follow a normal distribution with a mean of -15.5 and a standard deviation of 0.5. Viral infection promotes a rightward shift in ln (Kon) values, indicating an amplification of antibodies with faster binding coefficients.

**Figure 3b:**
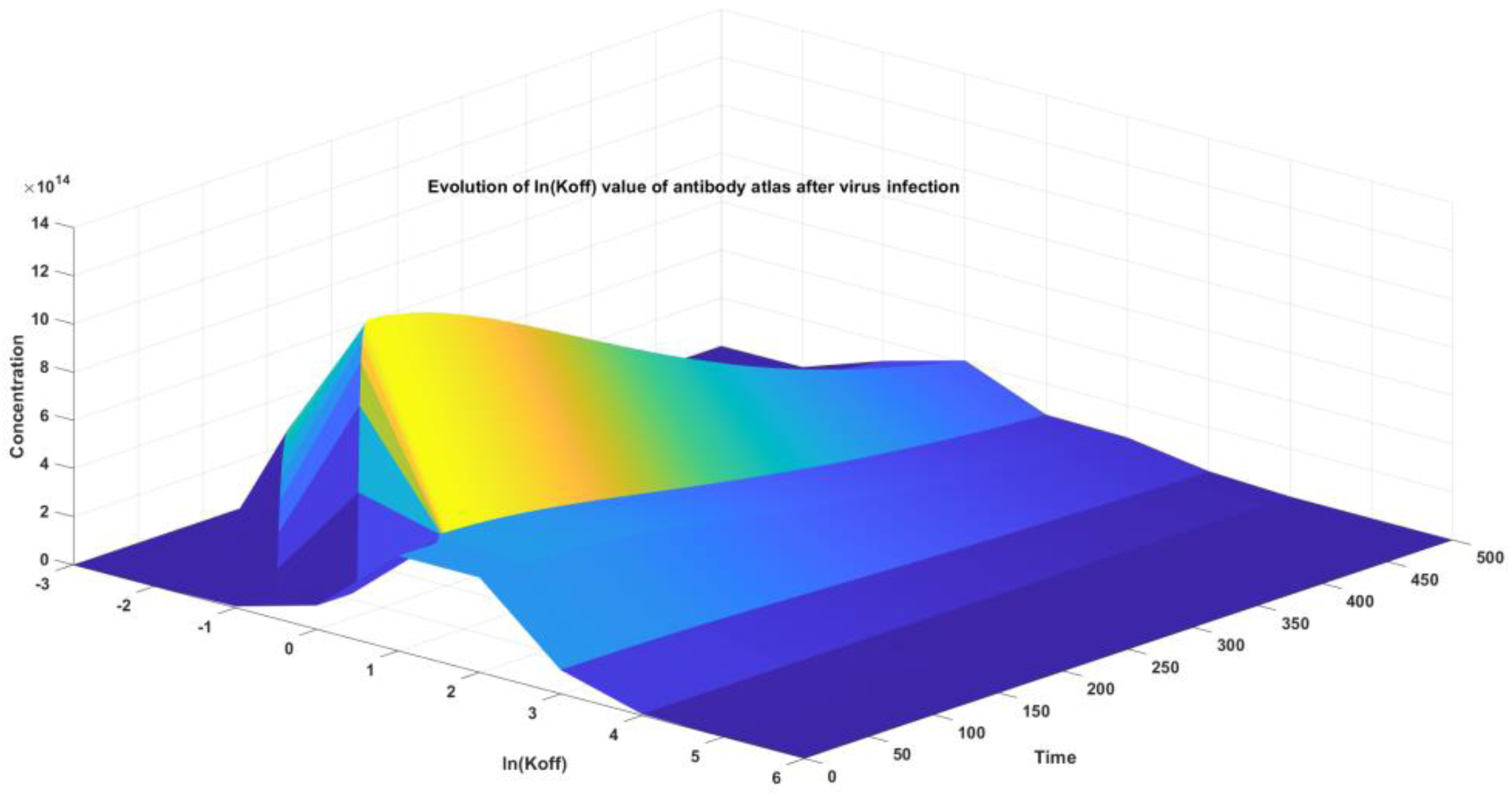
The evolutionary trend of antibody Koff value after virus infection. From the graph, it can be observed that the initial ln (Koff) values follow a normal distribution with a mean of 1.5 and a standard deviation of 0.8. Viral infection promotes a leftward shift in ln (Koff) values, indicating an amplification of antibodies with slower dissociation coefficients.

**Figure 3c:**
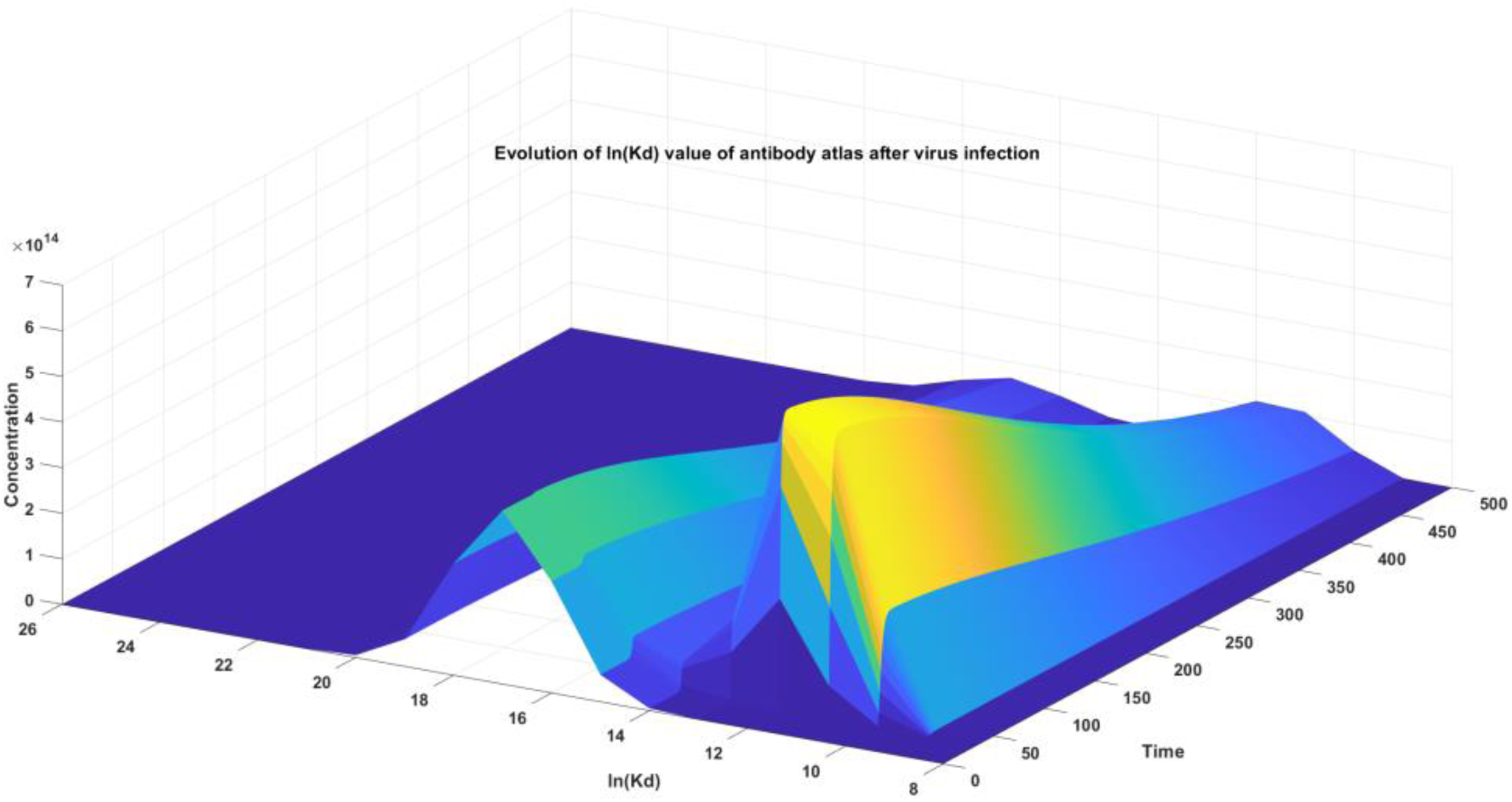
The evolutionary trend of antibody Kd value after virus infection. From the graph, it can be observed that the initial average value of ln (Kd) is 17. Viral infection promotes a rightward shift in ln (Kd) values, indicating an amplification of antibodies with stronger affinity.

From Figure 3d, it can be observed that during the viral infection process, there is a significant decrease in the Kd value within the compositional changes of the antibody library. However, the main contribution to this decrease in Kd value stems from the increase in K_on values (2/3 proportion), with a smaller contribution (1/3 proportion) originating from the decrease in K_off values.

**Figure 3d:**
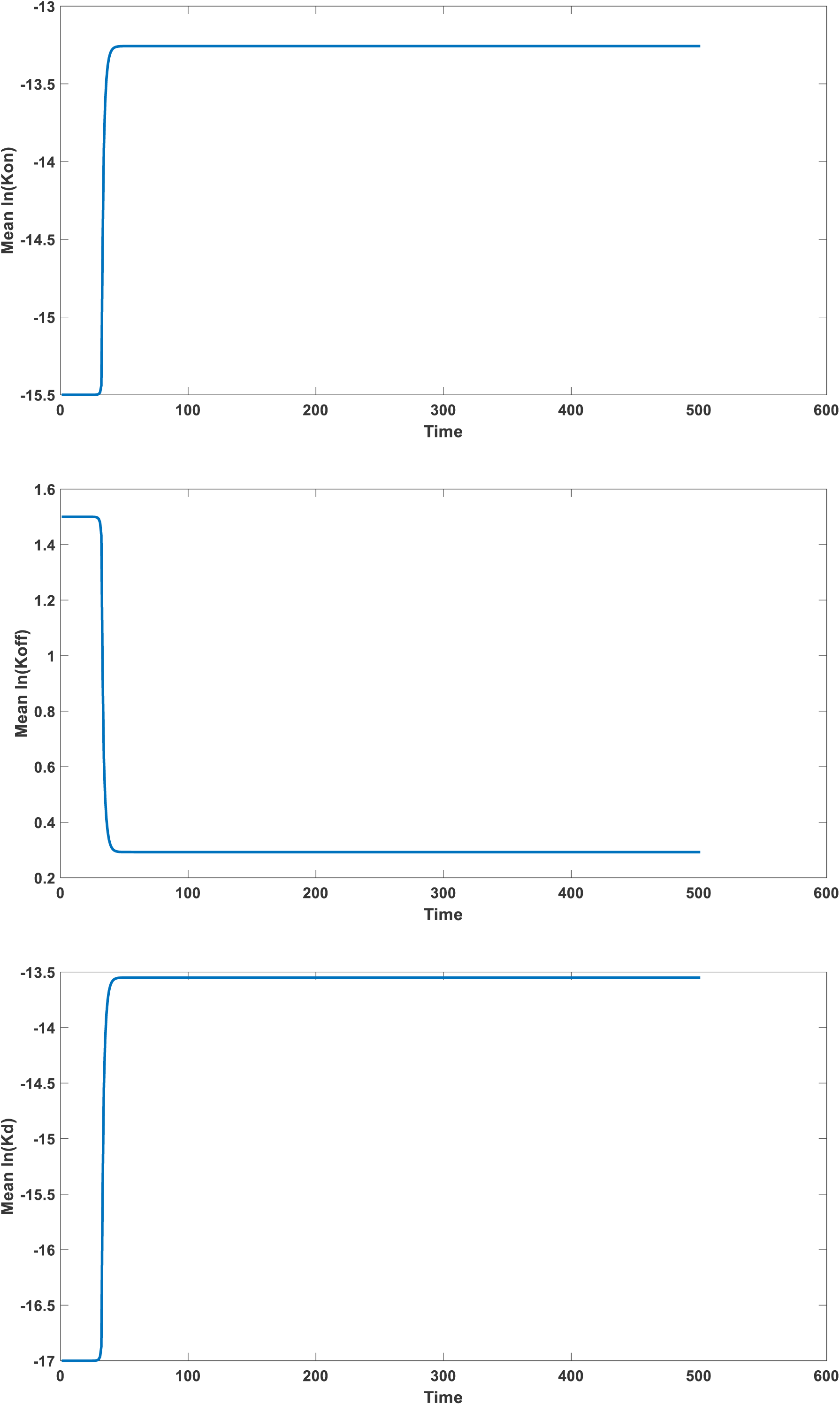
The immune system tends to proliferate antibodies with good Kon rather than the good Kd antibodies. The average value of ln (Kon) increased from -15.5 to -13.2. The average value of ln (Koff) decreased from 1.5 to 0.3. The average value of ln (Kd) decreased from 17 to 13.5, as shown in the graph.

Given that the organism tends towards changes in Kon values, it indicates that alterations in Kon values play a more significant role in viral clearance. Simulations of individual antibodies demonstrate that antibodies with excellent Kon properties have a greater advantage in inhibiting viral proliferation compared to those with excellent Kd properties. As illustrated in Figure 3e, when faced with the same initial virus concentration during infection, antibodies with excellent Kon properties exhibit lower peak viral loads, and the peak concentration of virus-antibody complexes is also lower than that of antibodies with excellent Kd properties. These findings underscore the critical role of forward binding coefficients in suppressing viral infection. Detailed information is provided as model 1.4 in supplementary materials.

**Figure 3e:**
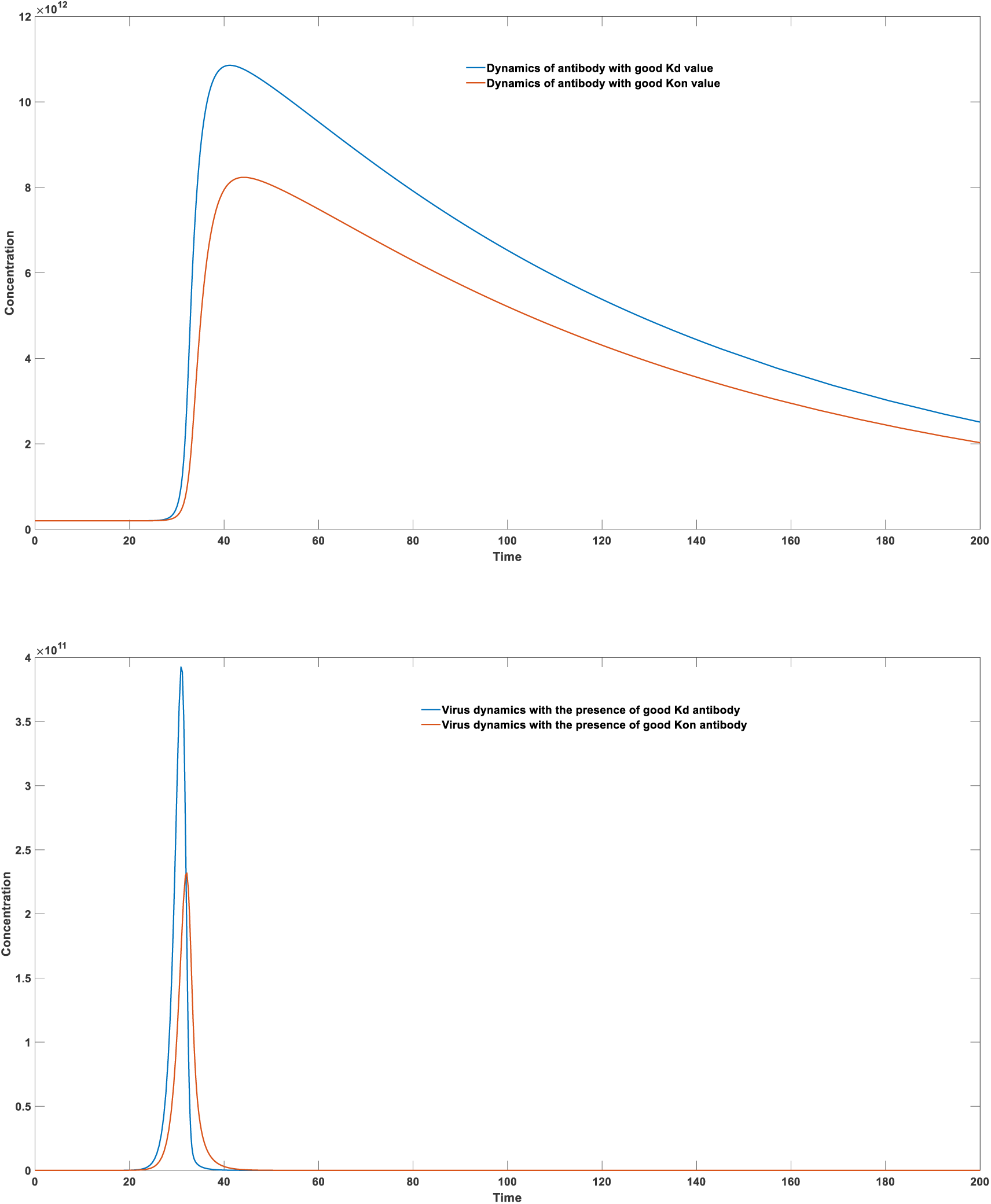
Antibodies with good K_fig3e. The upper graph shows the kinetic change curves of antibodies with favorable Kd values represented by the solid blue line, and antibodies with favorable Kon values represented by the solid red line. In the lower graph, the solid blue line represents the viral change curve when antibodies with favorable Kd values are present, while the solid red line represents the viral change curve when antibodies with favorable Kon values are present. It can be observed from the graph that the virus is more significantly inhibited when antibodies with favorable Kon values are present.

Based on the changes in the antibody atlas, we further computed the dynamic variations in ELISA results. Detailed information is provided as model 3 in supplementary materials, and the results are depicted in Figure 3f. It can be observed that there is no significant change in ELISA results during the early stages of infection, corresponding to the slow evolution of antibody components during the initial phase of infection. Subsequently, there is a sharp increase in ELISA results followed by a gradual decline. It is noteworthy that regardless of sample handling or the concentration gradient of antigen-like substances used for ELISA testing, the initial results will not be zero. Often, especially when samples are not pretreated with bovine serum, the initial values of ELISA results may be relatively large. However, this does not necessarily indicate a substantial proportion of high-affinity antibodies within the body. At this point, the concentration of high-affinity antibodies in the body may still be at a very low level. ELISA results primarily reflect the concentration of antibody-antigen complexes formed by antibodies with moderate binding affinities.

**Figure 3f:**
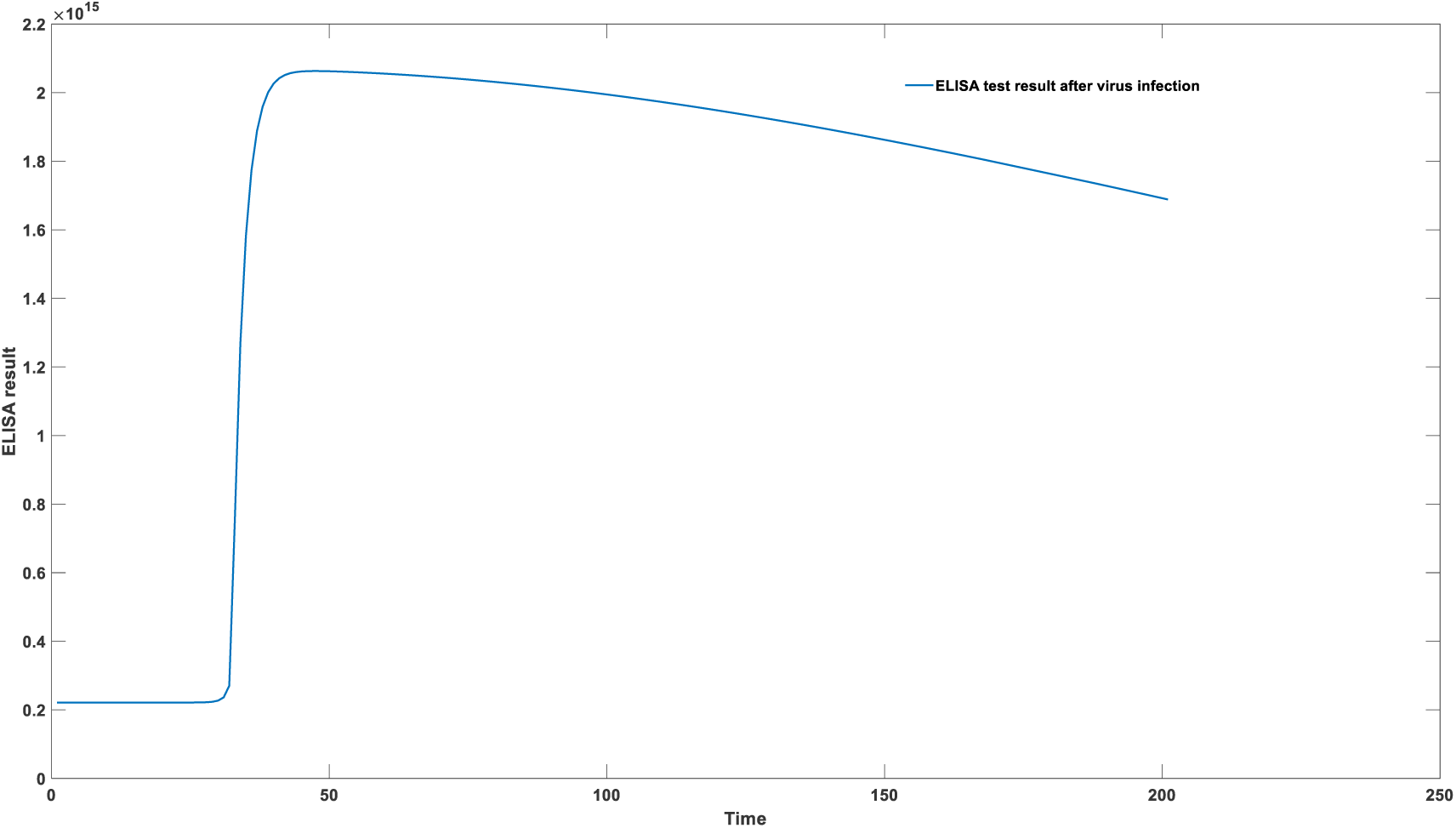
ELISA results after virus infection. The x-axis represents time, while the y-axis represents the concentration of antigen-antibody complexes. It can be observed from the graph that the ELISA results show a relatively stable plateau period followed by a rapid increase, and then a gradual decline.

### Mathematical simulation based on Antibody atlas mapping indicates the potential roles of self-antigens in maintaining B-cell homeostasis

Our model demonstrates that self-antigens play a crucial role in the long-term maintenance of antibodies. While antibodies exhibit a significant decline after reaching peak concentrations, this decline is not sustainable, with the rate gradually decreasing until stabilizing within a certain concentration range, accompanied by fluctuations. However, in the presence of self-antigen-like substances, the trend of antibody atlas changes, as depicted in the graph above in Figure 4a, where the decline in antibodies is more gradual, eventually stabilizing within a distributed range over the long term. For specific change plots, please refer to Extended Video 4. The specific trends of each antibody are displayed in Extended Figure 2. Conversely, in the absence of self-antigen-like substances, all antibodies show a rapid decline until eventual elimination, as illustrated in the lower graph of Figure 4a. Detailed information is provided as model 2.2 in supplementary materials.

**Figure 4a:**
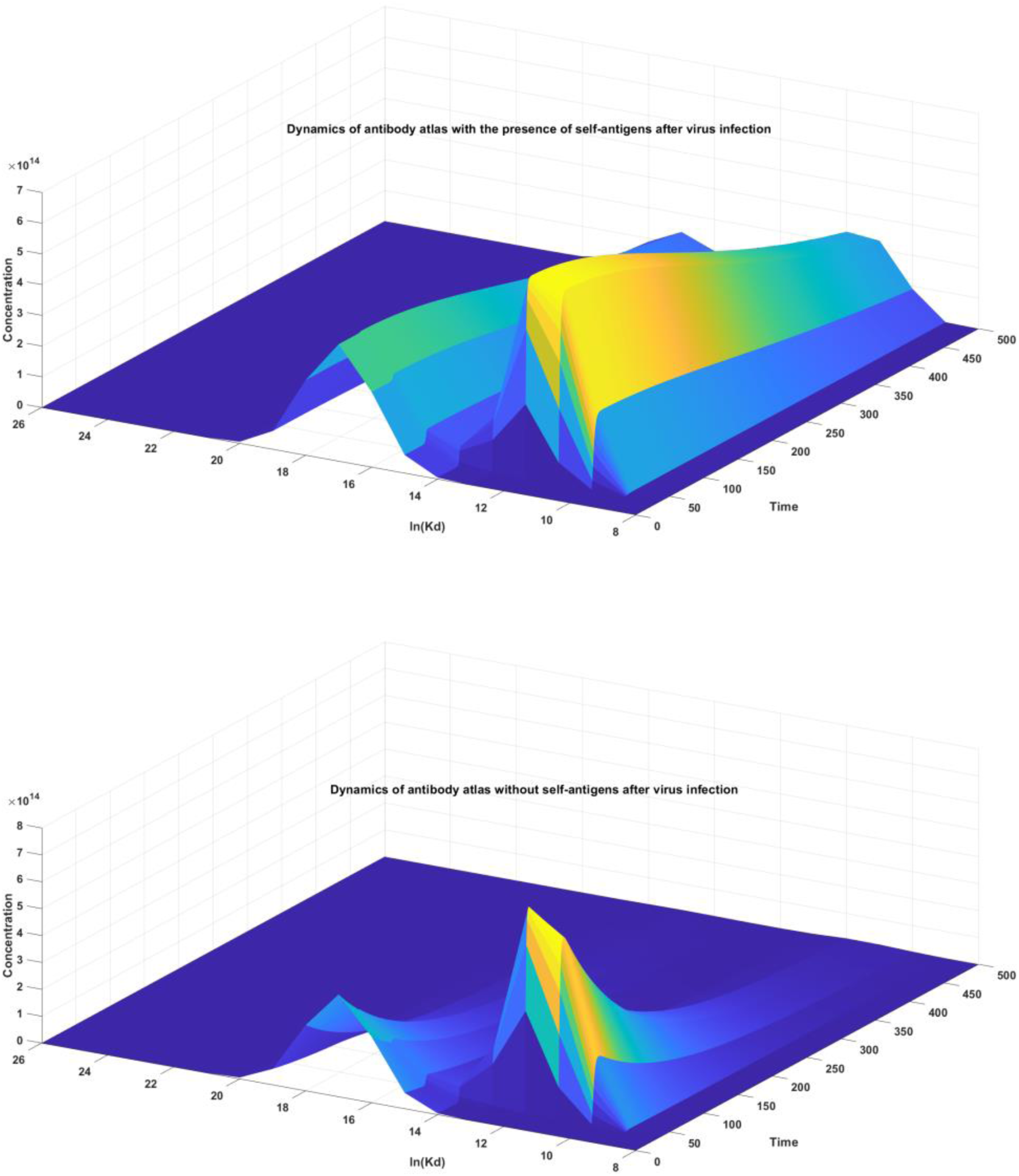
Comparison of antibodies dynamics with and without self-antigens. The figure above illustrates that in the presence of self-antigen, the reshaped antibody repertoire following viral infection will remain stable for an extended period. Conversely, the figure below indicates that in the absence of self-antigen, the reshaped antibody repertoire following viral infection will exhibit rapid decay.

In the long term, the presence of self-antigen-like substances not only decelerates the decay rate of high-affinity antibodies but also sustains them above a threshold for an extended period, thereby potentially providing lifelong protection to the host. Moreover, this sustaining relationship exhibits remarkable dynamics, with initial phases showing intriguing fluctuations. Specifically, there is a phenomenon of short-term increase in ELISA results, as indicated by the red dashed line in Figure 4b. This rebound effect has been validated in numerous long-term ELISA experiments [42–43], suggesting that it is not merely attributable to data error.

**Figure 4b:**
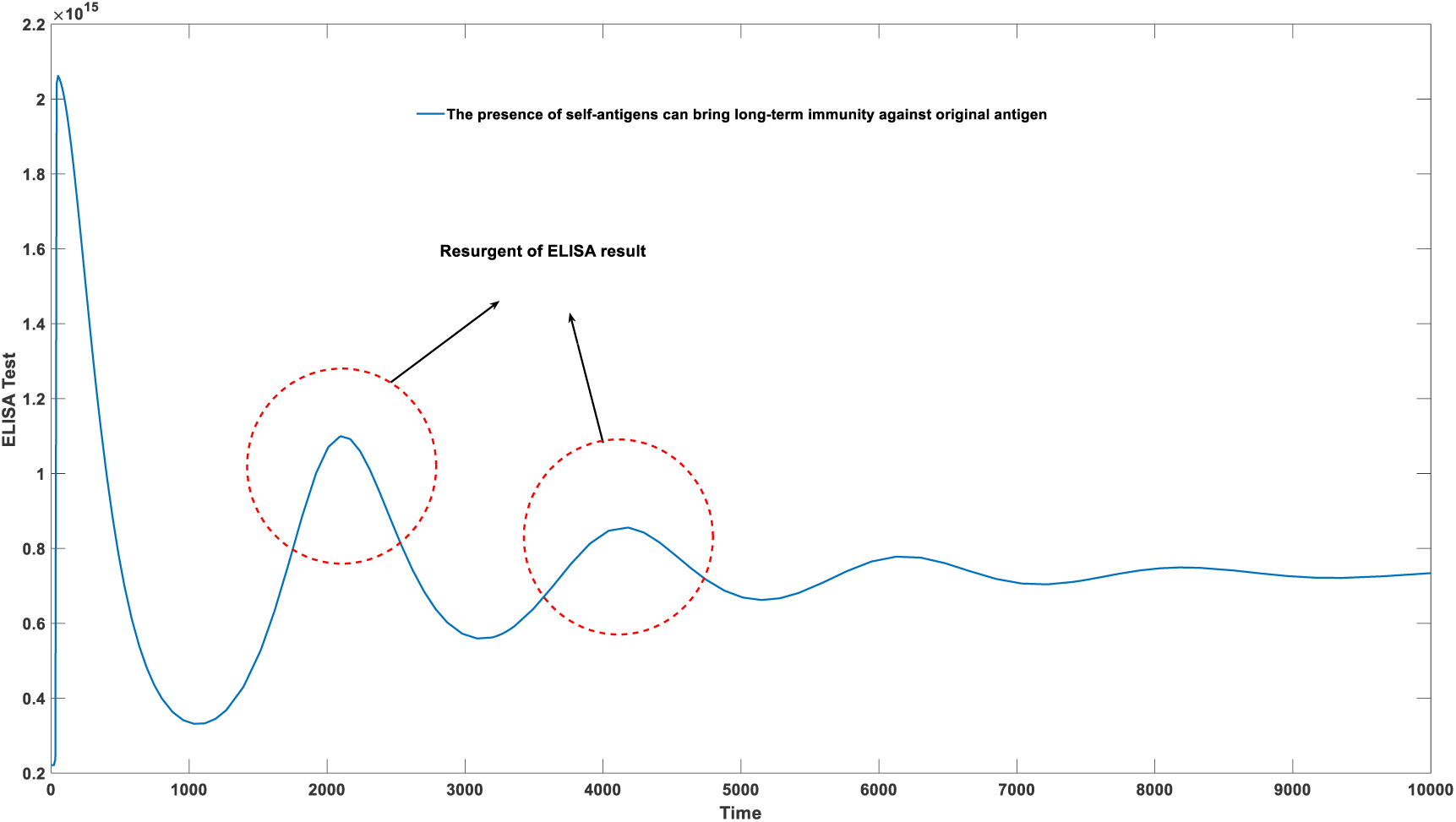
The presence of self-antigens can bring long-term immunity against virus with low mutation propensity. The red circles indicate a transient increase in the ELISA results.

Long COVID, or post-acute sequelae of SARS-CoV-2 infection (PASC), refers to the persistent symptoms experienced by some individuals following recovery from the acute phase of COVID-19. These symptoms can include fatigue, shortness of breath, cognitive impairment, and others, lasting weeks to months. The exact mechanisms underlying long COVID are not yet fully understood. Potential factors may include persistent viral presence, immune dysregulation, tissue damage, and neurological or psychological factors [44–45]. However, ongoing research aims to elucidate the complex interplay of these factors and identify effective treatments for those affected by this condition.

Our model suggests that the antigen-antibody complex serves as a direct indicator of symptom severity. Following acute infection, the virus is rapidly cleared with the generation of antibodies, leading to the swift disappearance of virus-antibody complexes, as depicted by the red solid line in Figure 4c. In contrast to the virus, self-antigen-like substances are continually replenished, thus when antibody concentrations are at higher levels, endogenous antigen-antibody complexes are not swiftly cleared, as illustrated by the yellow solid line in Figure 4c. Consequently, infected individuals may experience a prolonged immune response period, resulting in an extended duration of post-infection sequelae.

**Figure 4c:**
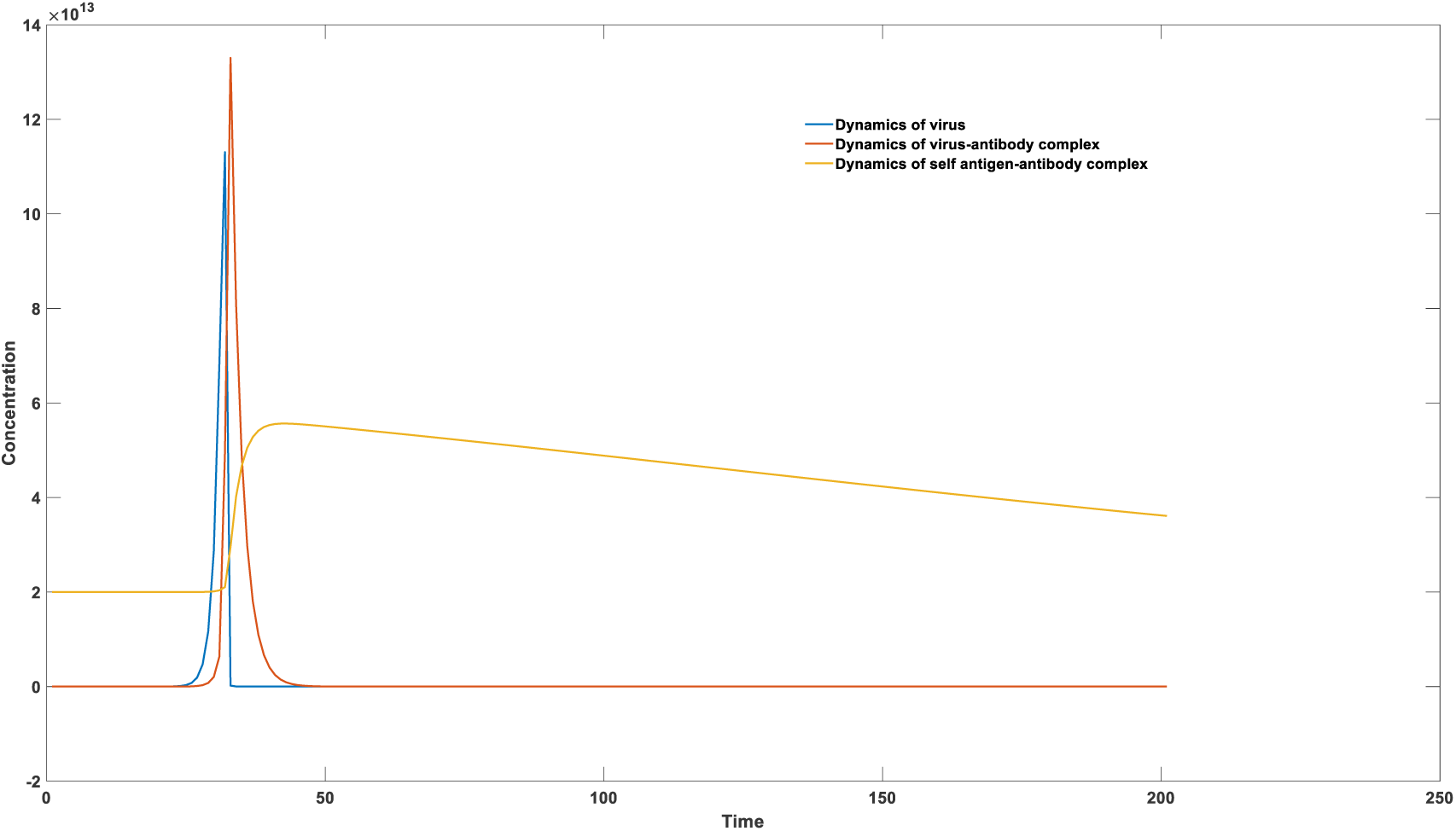
The presence of self-antigens may be a significant factor in long-term consequences of viral infections, such as long COVID. (comparison between virus-antibodies complex dynamics and self-antigen-antibodies complex dynamics). The solid blue line in the figure represents the dynamics of the virus, the solid red line represents the dynamics of the virus-antibody complex, and the solid yellow line represents the dynamics of the self-antigen-antibody complex. It can be observed from the figure that the virus-antibody complex rapidly diminishes after the infection ends, while the decline of the self-antigen-antibody complex is very slow. This may be one of the significant contributing factors to the development of post-infection sequelae.

### An exploration of the mechanism behind original antigenic sin

Original antigenic sin, also known as immunological imprinting or antigenic seniority, is a phenomenon observed in the immune response to pathogens, particularly viruses. It refers to the tendency of the immune system to preferentially recall and respond to previously encountered antigens, even when exposed to new, closely related antigens [46–48].

When an individual is exposed to a pathogen for the first time, such as during childhood, their immune system mounts a response and generates memory cells specific to the antigens of that pathogen. If the individual encounters a similar pathogen later in life with antigens resembling those of the original pathogen, the immune system may exhibit a biased response, primarily recalling and activating the memory cells specific to the original antigen. This phenomenon can potentially limit the effectiveness of the immune response against the new pathogen, as it may not produce antibodies or cells optimized for neutralizing or clearing the new threat.

One of the key unknowns in the study of original antigenic sin is the extent to which it impacts the effectiveness of vaccines and natural immunity against evolving pathogens. Researchers are still investigating how pre-existing immunity, shaped by previous exposures to related pathogens or vaccines, influences the immune response to novel strains or variants of viruses like influenza, dengue, and SARS-CoV-2 [48–50].

Additionally, the mechanisms underlying original antigenic sin are not fully elucidated. It is unclear why the immune system prioritizes memory cells specific to previous antigens over generating a response tailored to the current threat. Further research is needed to understand the molecular and cellular processes that govern this phenomenon, which could inform the development of more effective vaccines and therapeutic strategies.

Moreover, the implications of original antigenic sin for public health and disease control remain a topic of interest and debate. Understanding how prior immune experiences shape responses to new pathogens or vaccine formulations is crucial for designing vaccination campaigns, predicting disease outcomes, and managing infectious disease outbreaks effectively.

We have detailed the mechanism underlying the formation of original antigenic sin, as illustrated in Figure 5a. Prior to viral invasion, the binding coefficients of the corresponding antibody repertoire to the viral antigen exhibit a characteristic normal distribution. However, following viral infection, the infection reshapes the composition of the antibody repertoire, as depicted in the blue dashed box on the right. Whether it is the primary infection with the original strain or a variant strain, the antibody composition undergoes significant reshaping due to the insufficient concentration of initially high-binding affinity antibodies. If the host has not been previously infected with the original strain but is directly infected with the variant strain, there is a substantial amplification of excellent antibodies targeting the variant strain antigen, showing a high proportion of high-binding affinity antibodies. Conversely, when the host has been previously infected with the original strain, the distribution of antibodies targeting the variant strain antigen is altered, as depicted in the lower-left graph of Figure 5a. The levels of initially high-binding affinity antibodies increase, but their distribution levels are insufficient to counter the intracellular proliferation of the variant strain, resulting in a secondary infection phenomenon. At this point, the levels of high-binding affinity antibodies are elevated. The specific parameter processes are outlined in the supplementary materials. As illustrated in the lower-right graph of Figure 5a, however, due to their initially high levels, the elevation of these levels is not substantial enough, leading to effective suppression of the virus. However, the proportion of ultimately formed high-binding affinity antibodies is significantly smaller than in the absence of original antigenic sin.

**Figure 5a:**
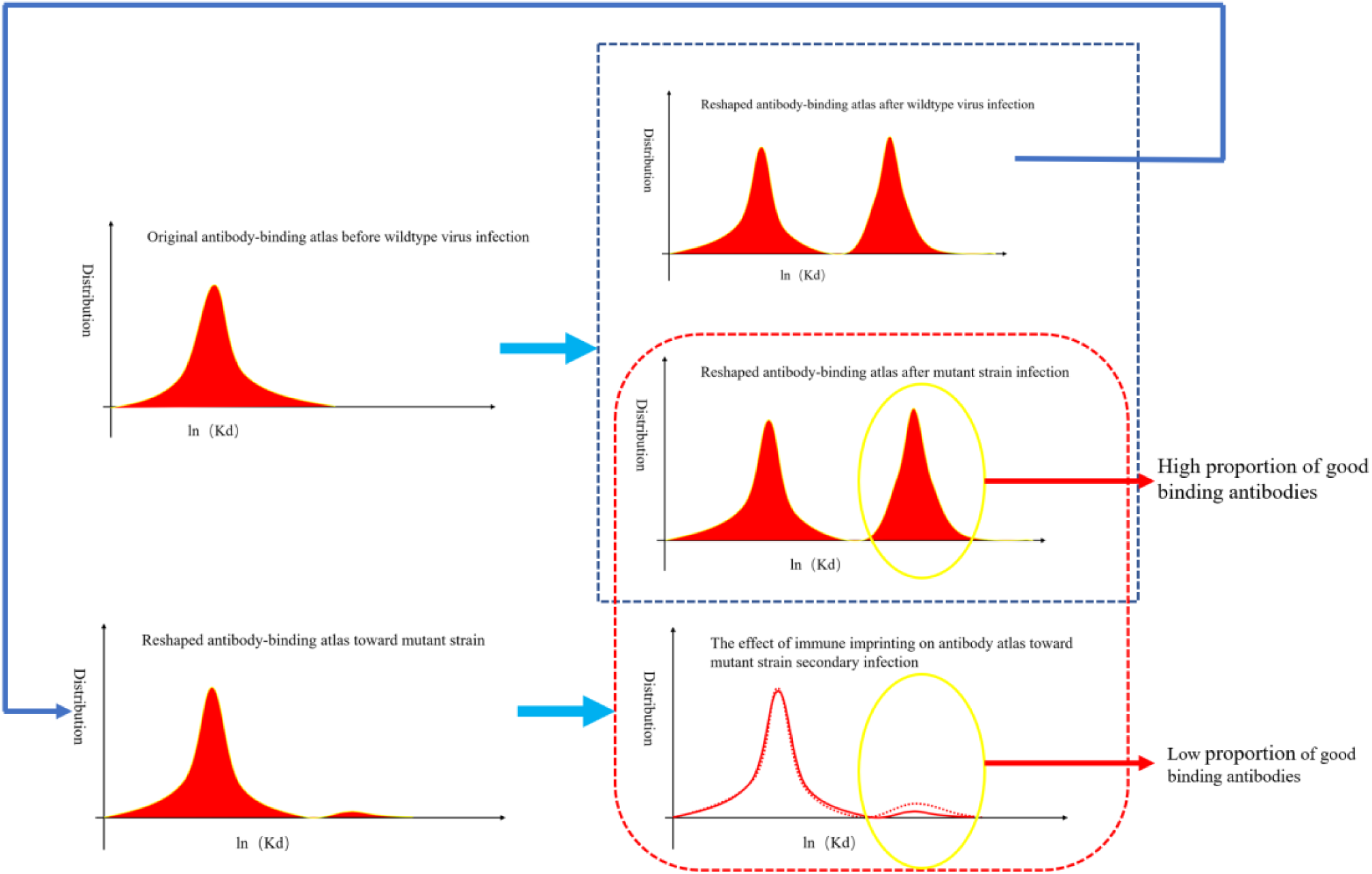
The drift of antibody binding profiles in response to mutant strains. The blue dashed box in the figure represents the changes in the antibody repertoire of the host during the initial infection with the original virus or mutant strain. If the host in the late stage, as shown in the upper figure, is reinfected with the mutant strain, these reshaped antibody repertoires will exhibit a profile similar to that in the lower left figure when facing the antigen of the mutant strain. Secondary infection will lead to the proliferation of good-binding antibodies, but the magnitude of this increase will be significantly reduced compared to the control group. Moreover, since most of these antibodies are derived from seeds provided by antibodies from the initial infection, the vast majority of antibodies generated against the mutant strain can cross-react with the wild type.

The phenomenon of secondary infection resulting from variant strains is further elucidated in Figure 5b, providing a more direct illustration. The figure illustrates that the mutation effect of the virus can lead to secondary infections, albeit typically less severe than the severity of primary infections, thanks to the effect of immune imprinting. However, it is important to note that due to the presence of immune imprinting, secondary infections with variant strains often do not result in the elicitation of high levels of specific antibodies targeting the variant strain.

**Figure 5b:**
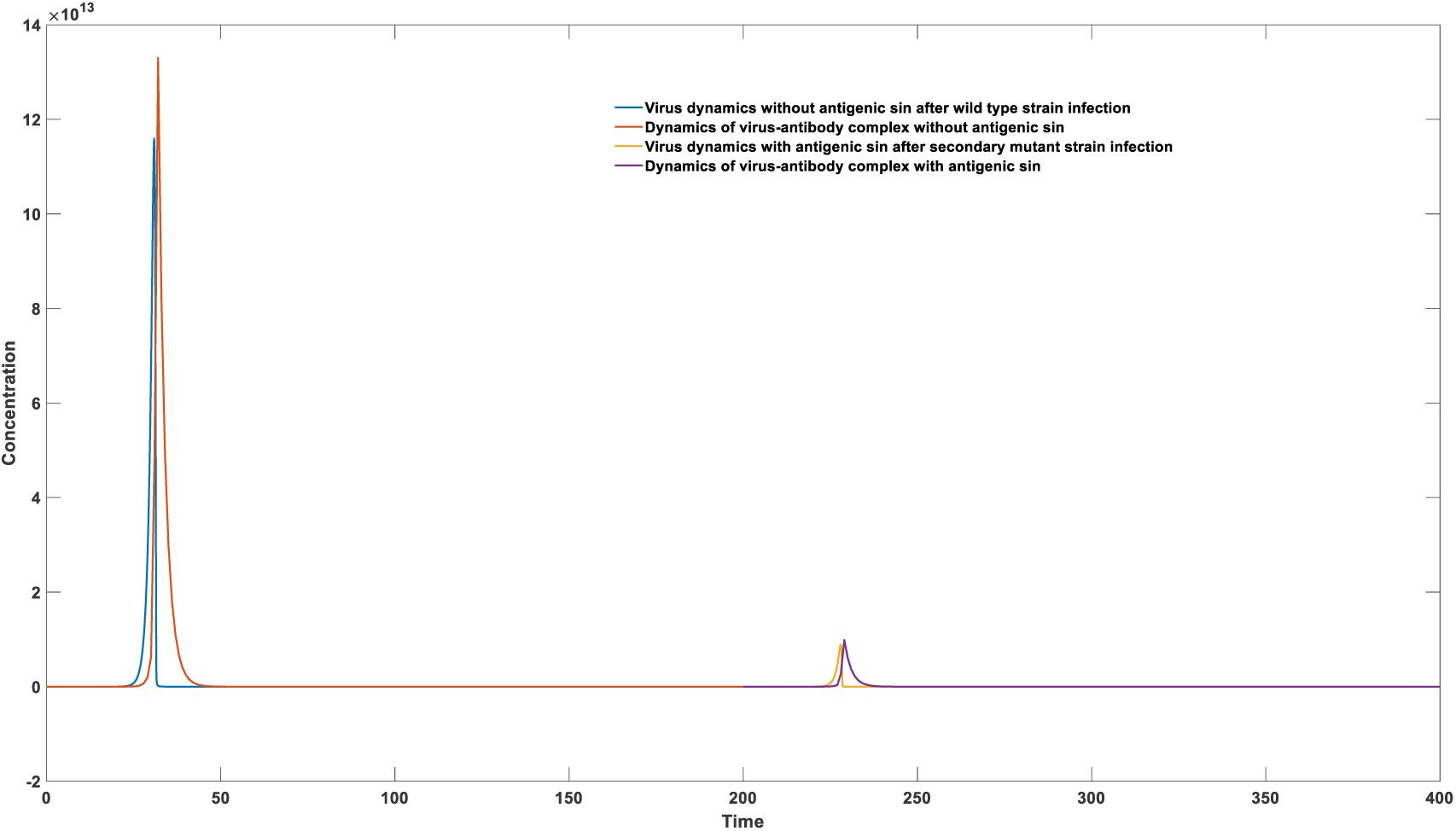
Mutation strain can lead to secondary infection. From the figure, it can be observed that the reshaping of the antibody repertoire during the initial infection does not completely prevent the proliferation of the mutant strain. However, the peak virus concentration and peak virus-antibody complex concentration resulting from secondary infection are greatly reduced compared to the initial infection.

The disparities in the reshaping of the organism’s antibody atlas induced by immune imprinting are depicted through a more comprehensive dynamic process in Figure 5c. As depicted in Figure 5c, in the absence of original antigenic sin, high-affinity antibodies targeting the variant strain are comprehensively elicited. However, in the presence of original antigenic sin, the elicitation of high-binding affinity antibodies is subject to a certain degree of inhibition. A more intuitive depiction of these changes can be referenced in Extended Video 5. The specific trends of each antibody are displayed in Extended Figure 3. Detailed information is provided as model 2.3 in supplementary materials.

**Figure 5c:**
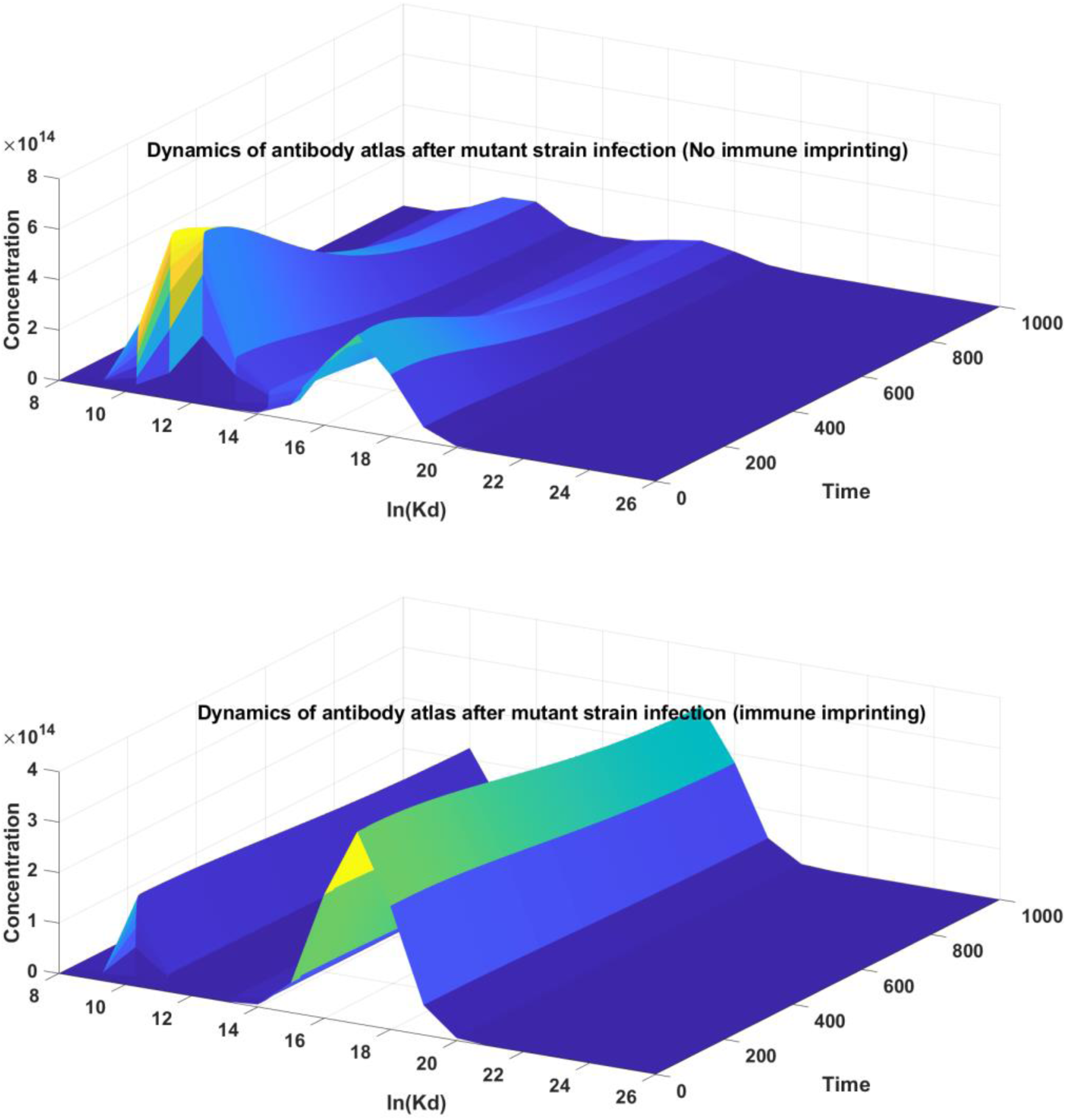
Comparison of antibody dynamics with and without original antigen sin. From the figure, it is evident that in the absence of immune imprinting, the proliferation amplitude of high-affinity antibodies is significantly greater than in the presence of immune imprinting.

When employing ELISA assays to quantify the levels of specific antibodies, the results, as depicted in Figure 5d, indicate a marked reduction in ELISA readings in the presence of original antigenic sin compared to its absence. Detailed information is provided as model 3 in supplementary materials.

**Figure 5d:**
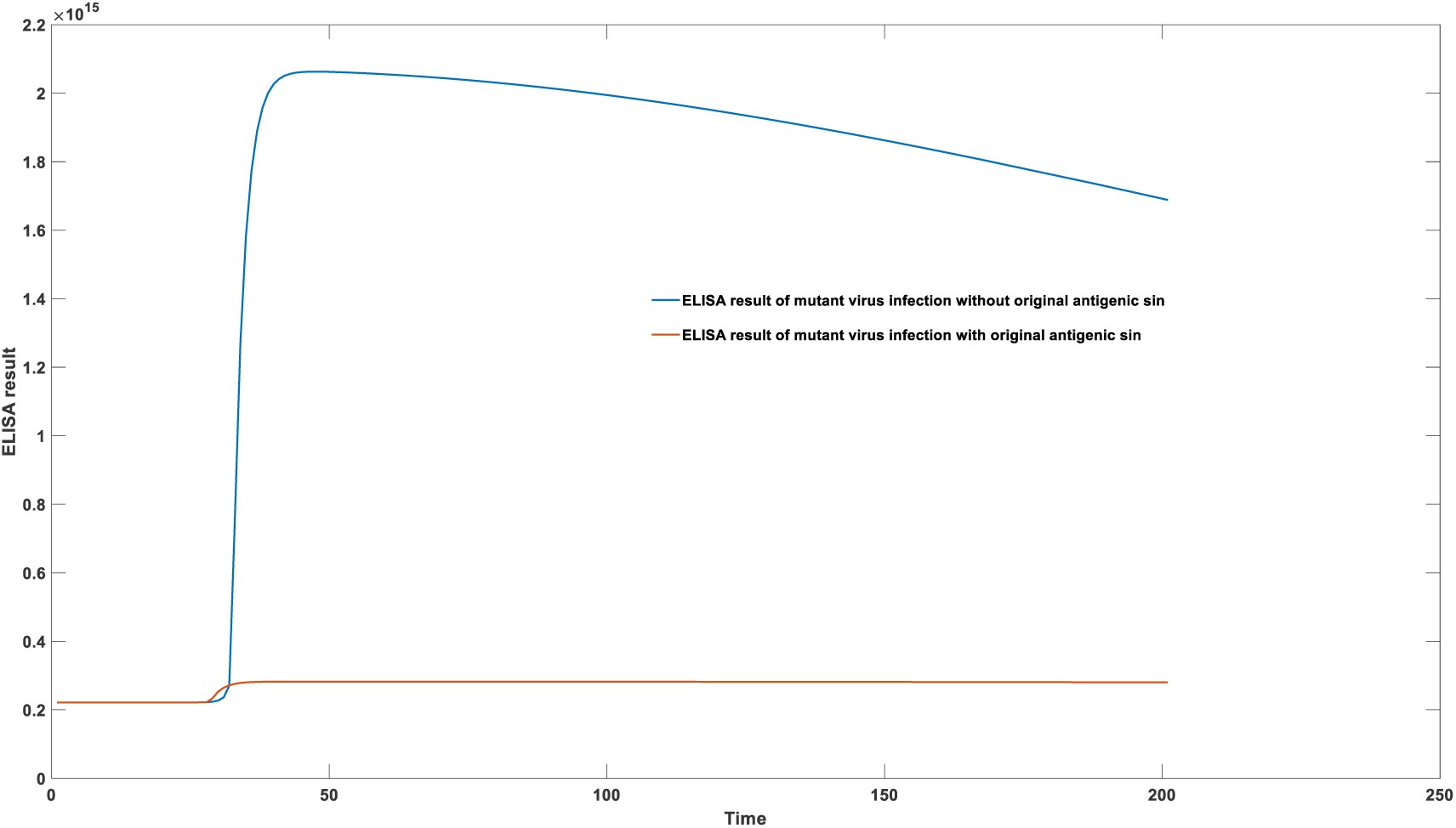
Comparison of ELISA results with and without original antigenic sin. From the figure, it can be observed that when original antigenic sin is present, the magnitude of the increase in ELISA following a secondary infection is significantly smaller than the initial infection in the absence of original antigenic sin.

While original antigenic sin may pose obstacles to antibody generation, it can remarkably mitigate the severity of secondary infections. As depicted in Figure 5e, in the presence of original antigenic sin, the initial high-affinity antibody levels result in significantly reduced peak viral loads and peak concentrations of antigen-antibody complexes compared to scenarios without immune imprinting. This phenomenon also elucidates many vaccine’s cross-protective effects [51–53].

**Figure 5e:**
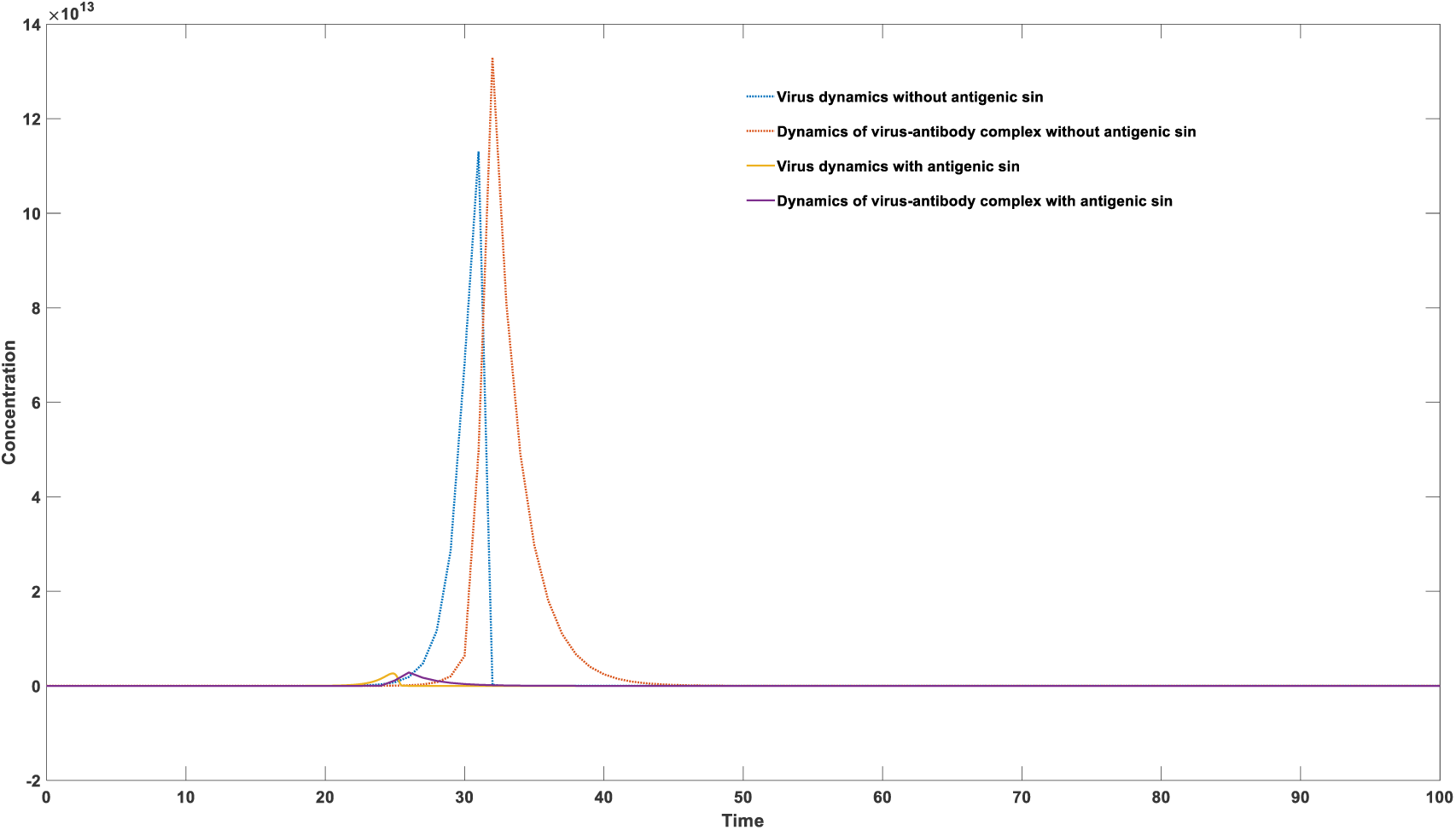
The original antigenic sin is beneficial in reducing infection symptom but would limit the proliferation of good binding antibodies. From the figure, it can be observed that both the peak virus concentration and the peak concentration of virus-antibody complexes are significantly lower during secondary infections in the presence of original antigenic sin compared to the primary infection. This suggests that original antigenic sin plays a beneficial role in reducing infection symptoms.

In many scenarios, there is a desire to elevate specific antibodies targeting mutant strains to sufficiently high levels to entirely prevent secondary infections caused by these strains. In such instances, the aim is to overcome the antibody generation impediments posed by immune imprinting, as illustrated in Figures 5c and 5d. Through natural infection, reversing immune imprinting is unattainable because the presence of a certain concentration of initial high-affinity antibodies prevents extensive viral replication, thus hindering the elevation of specific antibodies to a higher level. However, a single injection of a large quantity of antigen can reverse this trend, as exemplified by vaccination. As demonstrated in Figure 5f, when the injected dose is adequate, antigen administration for variant strains can eliminate the antibody proliferation barriers induced by original antigenic sin. This notion has been substantiated by numerous immunological experiments [54–55]. Detailed information is provided as model 2.4 in supplementary materials.

**Figure 5f:**
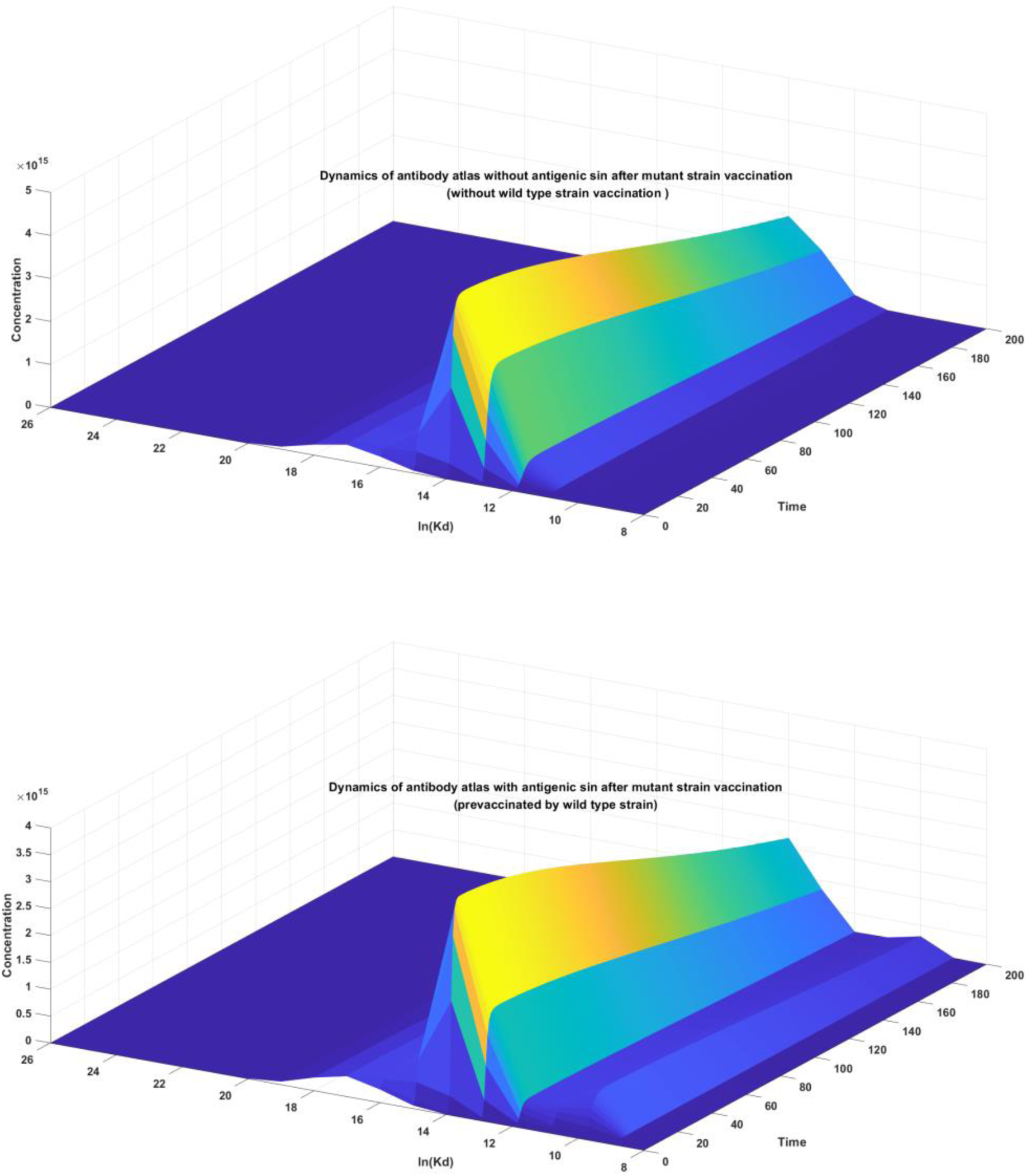
The original antigenic sin can be eliminated with booster vaccination. From the figure, it can be observed that following vaccination with the antigen of the mutant strain, the host can effectively achieve proliferation of high-affinity antibodies, thereby eliminating the antibody proliferation barrier caused by original antigenic sin.

### Parameter estimation and its application

One significant application of mathematical models lies in their predictive capabilities. This necessitates a comprehensive and reliable parameter set, inevitably involving parameter fitting. In our model, each parameter carries a corresponding physical significance, thus solving for parameters concerning populations or individuals can reflect differences in certain immunological traits. Additionally, although our model involves the dynamic changes of antibody atlas, the parameters requiring fitting are minimal, enhancing the reliability of parameter fitting. Specifically, our parameters primarily fall into two categories. One category involves parameters related to the initial distribution of antibodies. Since both (Kon) and (Koff) collectively determine (Kd), but (Kon) has a greater impact on antibody variation, we can assume (Koff)’s distribution to be a fixed value, fitting only parameters of the distribution of ln(Kon), such as mean and variance. The second category of parameters pertains to the interaction between the immune system and viruses, mainly comprising the virus proliferation coefficient, the antibody regeneration coefficient concerning specific virus-antibody complexes (primarily associated with helper T cell activity), and the clearance rate of virus-antibody complexes (mainly related to NK cells).

For fitting the first category of parameters, which does not involve the viral infection process, we primarily utilize ELISA dynamic data before host infection. Here, dynamic data refers not to ELISA data at different time points but rather the variation in ELISA data within the same sample testing process, encompassing more data information, including the situation of antibody (Kon) values, while the final ELISA result can only reflect the equilibrium constant (Kd). This parameter testing group includes only two parameters, the mean and variance of ln(Kon), and through the use of MCMC [56–58], we can accurately estimate their values. Once we obtain the numerical values of the first category of parameters, we can further estimate the second category of parameters based on ELISA results of patients during the virus infection process.

The purpose of parameter estimation is not to obtain the absolute numerical values of specific parameters but to reflect differences in parameters among different individuals or populations, thereby determining differences in their immunological characteristics. For changes in antibody atlas caused by infection with the same virus, we can consider the virus proliferation coefficient as a constant rather than a parameter. We need to fit two additional parameters, namely the antibody regeneration coefficient concerning virus-antibody complexes (primarily associated with helper T cell activity) and the clearance rate coefficient of virus-antibody complexes (mainly related to NK cells).

Using artificially set datasets, we demonstrate that the MCMC method can reproduce the original parameters, as shown in Figure 6. This indicates the effectiveness of our model in parameter fitting. The specific parameter fitting process is detailed in the methods section.

**Figure 6:**
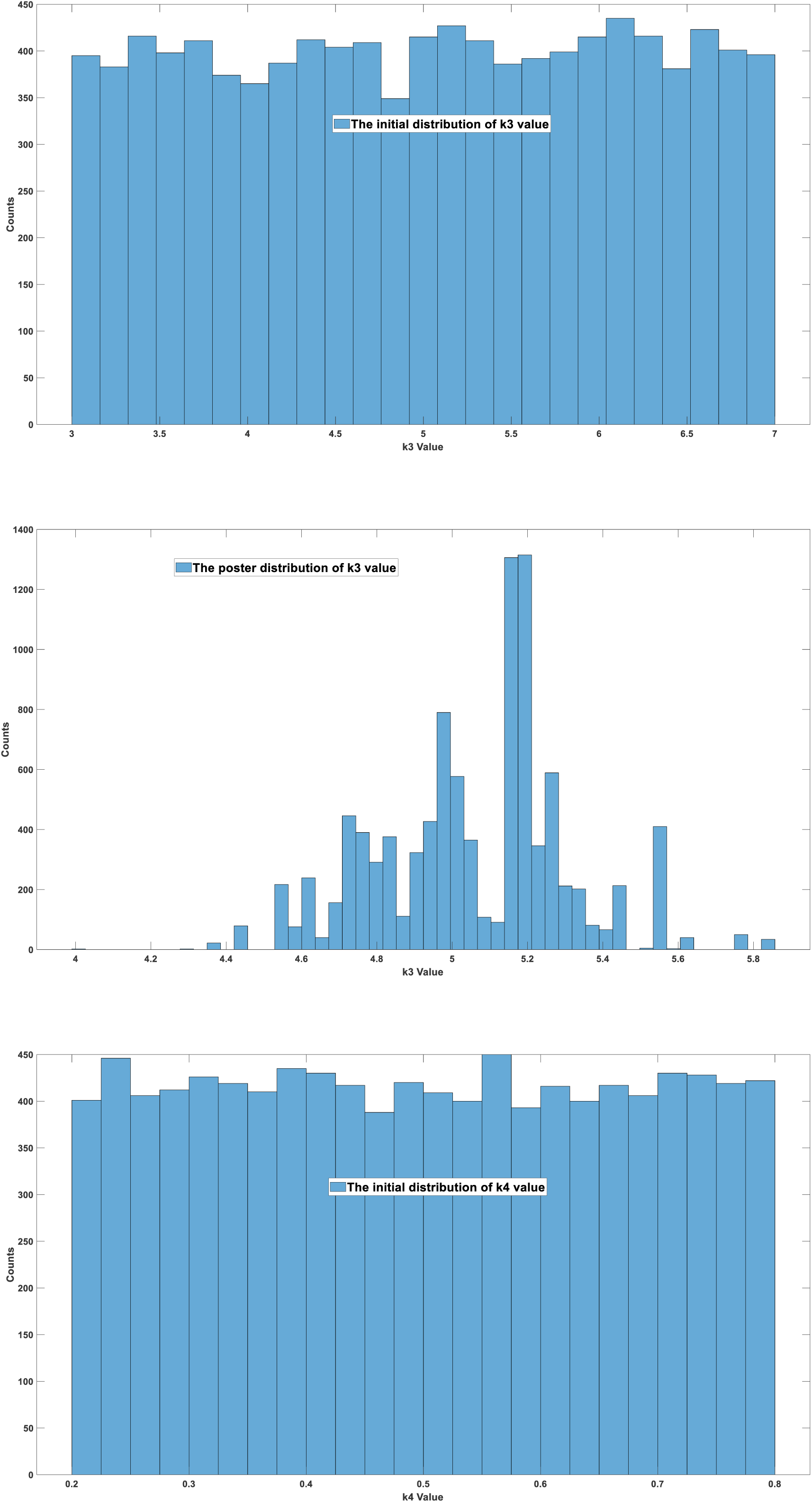
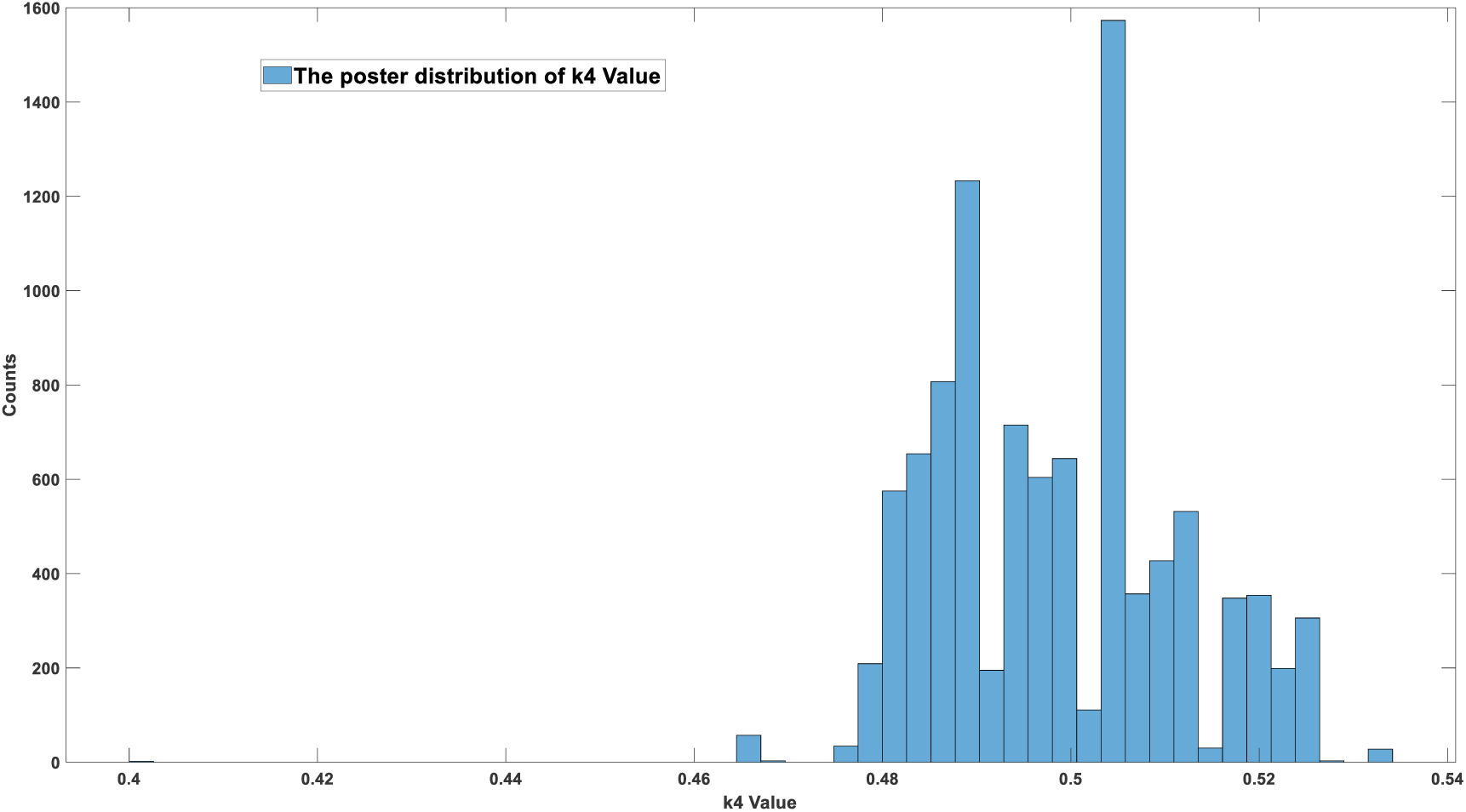
The prior and poster parameter distribution using MCMC method

We further conducted a sensitivity analysis on the parameters, and the detailed principles are described in the methods section. Specifically, we categorized these parameters into two types. The first type describes the static ability of the host to resist viral infection, mainly referring to the mean and standard deviation of ln(Kon) and ln(Koff), which represent the existing antibody binding parameters. It is evident that individuals with larger ln(Kon) and smaller ln(Koff) values possess stronger resistance to viral infection. Additionally, a larger standard deviation also confers increased resistance to viral infection. Here, we employed the same absolute fluctuation value to conduct sensitivity analysis on the parameters, including five characteristic representations: viral infectivity (total viral load over the infection period = integral of viral concentration at individual time points over the infection period), peak viral load, peak viral-antibody complex load (indicator of symptom severity), peak concentration in ELISA test (peak concentration of all binding antibodies), and long-term ELISA result (long-term sustained concentration of all binding antibodies).

The other type of parameters describes the dynamic ability of the host to resist viral infection, primarily encompassing the viral proliferation coefficient k1, the regeneration coefficient of antibody against the viral-antibody complex k3, and the clearance rate of the viral-antibody complex k4. Here, we employed a 10% relative fluctuation in parameter values to conduct sensitivity analysis. The results are presented in Table 1.

**Table 1:** Parameter sensitivity analysis results.

| Parameter Name | Virus transmission capacity | Peak viral loading | Peak virus-antibody concentration | Maximal ELISA result | Long-term ELISA result |
| --- | --- | --- | --- | --- | --- |
| Mean(lnKon) +1 | 216.54 | 252.05 | 225.04 | 5.80 | 0.43 |
| Std(lnKon) +0.1 | 1.58 | 1.58 | 2.39 | 0.88 | 0.39 |
| Mean(lnKoff) +1 | 0.06 | 0.07 | 0.26 | 0.05 | 0.01 |
| Std(lnKoff) +0.1 | 0.02 | 0.03 | 0.0056 | 0.022 | 0.0091 |
| $k_1$ +10% | 4.71 | 5.97 | 9.14 | 5.63 | 1.19 |
| $k_3$ +10% | -2.45 | -2.26 | -2.33 | -1.77 | -0.57 |
| $k_4$ +10% | 0.62 | 0.59 | 0.63 | -0.56 | -1.58 |

Table 1 indicates that the forward binding coefficient Kon significantly influences the severity of infection more than the reverse binding coefficient, which is consistent with our aforementioned conclusion. Moreover, it can be observed that the viral proliferation coefficient k1 is positively correlated with these five indicators, while k3 is negatively correlated with them. Interestingly, the clearance rate of the viral-antigen complex k4 is positively correlated with the first three indicators, but negatively correlated with the latter two. This suggests that an increase in the value of k4 will enhance the transmission ability of the virus, potentially exacerbating the symptoms of host infection. Simultaneously, an increased clearance capacity of the viral-antibody complex is disadvantageous for stimulating the production of more antibodies by the host, leading to lower ELISA levels. Furthermore, the value of k4 is closely related to the activity and quantity of NK cells, indicating that enhanced NK cell-mediated viral phagocytosis may lead to obstacles in antibody generation, a hypothesis supported by direct experimental evidence [75–76].

For the initial infection of the same unknown virus, different age groups often exhibit different parameter distributions. Elderly individuals typically have larger variances in static parameters due to the greater diversity of antibody types within their bodies. Increased variance enhances the ability to resist viral infection. Conversely, younger individuals often possess superior dynamic abilities to resist viral infection, such as larger k3 values, attributed to the stronger cytokine-secreting ability of their helper T cells in mediating B cell proliferation.

### Antibody atlas considering IgM-IgG isotype switching

Antibody isotype switching, particularly from IgM to IgG, is a process where B cells change the type of antibody they produce. Initially, B cells produce IgM antibodies, which are crucial for early immune responses. However, through isotype switching, B cells can mature into plasma cells that produce IgG antibodies, offering longer-lasting immunity and improved effectiveness against pathogens. This switch is vital for adaptive immunity, providing a diverse range of antibodies tailored to combat specific threats [59–60].

Considering antibody isotyping, the dynamics of the antibody atlas could be more intricate. Here, we delve into the kinetic changes of the antibody repertoire resulting from the isotype switching between IgM and IgG. We delineate the antibody profiles separately for IgG and IgM. As depicted in Figure 7a, for antibody repertoires not exposed to specific viral infections, IgM and IgG exhibit similar mean values in ln(Kon) and ln(Koff), yet with distinct variances. Given that IgG predominantly derives from the transformation of IgM post-infection, IgG manifests a smaller variance when encountering novel antigens. Conversely, owing to the broader diversity of IgM, its binding affinity to novel antigens displays a larger variance. Moreover, empirical data suggest that immunoglobulins within the human body are primarily IgG, with concentrations several to over ten times that of IgM [61]. Therefore, in our simulations, we set the overall concentration of IgG to be tenfold that of IgM.

**Figure 7a:**
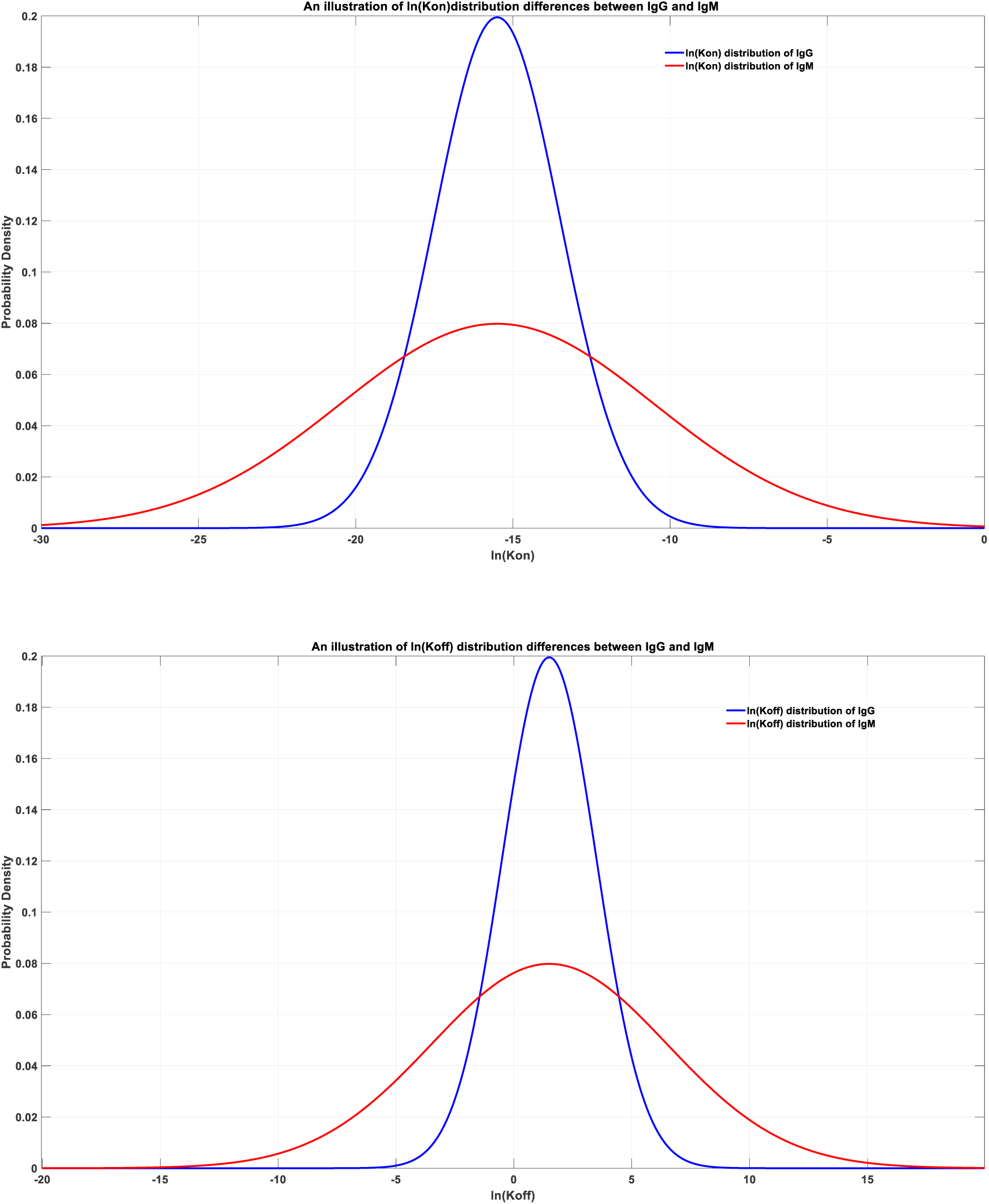
An illustration of differences in the distribution of IgG and IgM before virus infection. From the figure, it can be observed that both the Kon and Koff values for IgM exhibit a broader distribution and larger variance compared to IgG.

The variance in the distribution of binding coefficients of IgM to unknown antigens is greater, leading to the initial antibodies with high binding activity to unknown antigens primarily originating from IgM. Consequently, during the initial infection, IgM undergoes rapid proliferation. As illustrated in Figure 7b, the proliferation of high-binding activity IgM significantly precedes that of IgG. In the later stages, the proliferation of high-affinity IgG primarily stems from two sources. Firstly, it arises from the transformation of IgM, which also accelerates the decay of IgM, as evidenced by the rapid decline of IgM following its peak concentration. Secondly, it originates from the proliferation of IgG itself under antigenic stimulation. Furthermore, Figure 7b reveals that although IgM initiation precedes IgG, IgM experiences rapid decay, whereas IgG decay is more delayed, ultimately maintaining at a higher threshold level. Thus, IgG provides long-term protective mechanisms. For a more intuitive understanding of the dynamic changes, please refer to Extended Video 6. The specific trends of each antibody are displayed in Extended Figure 4. Detailed information is provided as model 4 in supplementary materials.

**Figure 7b:**
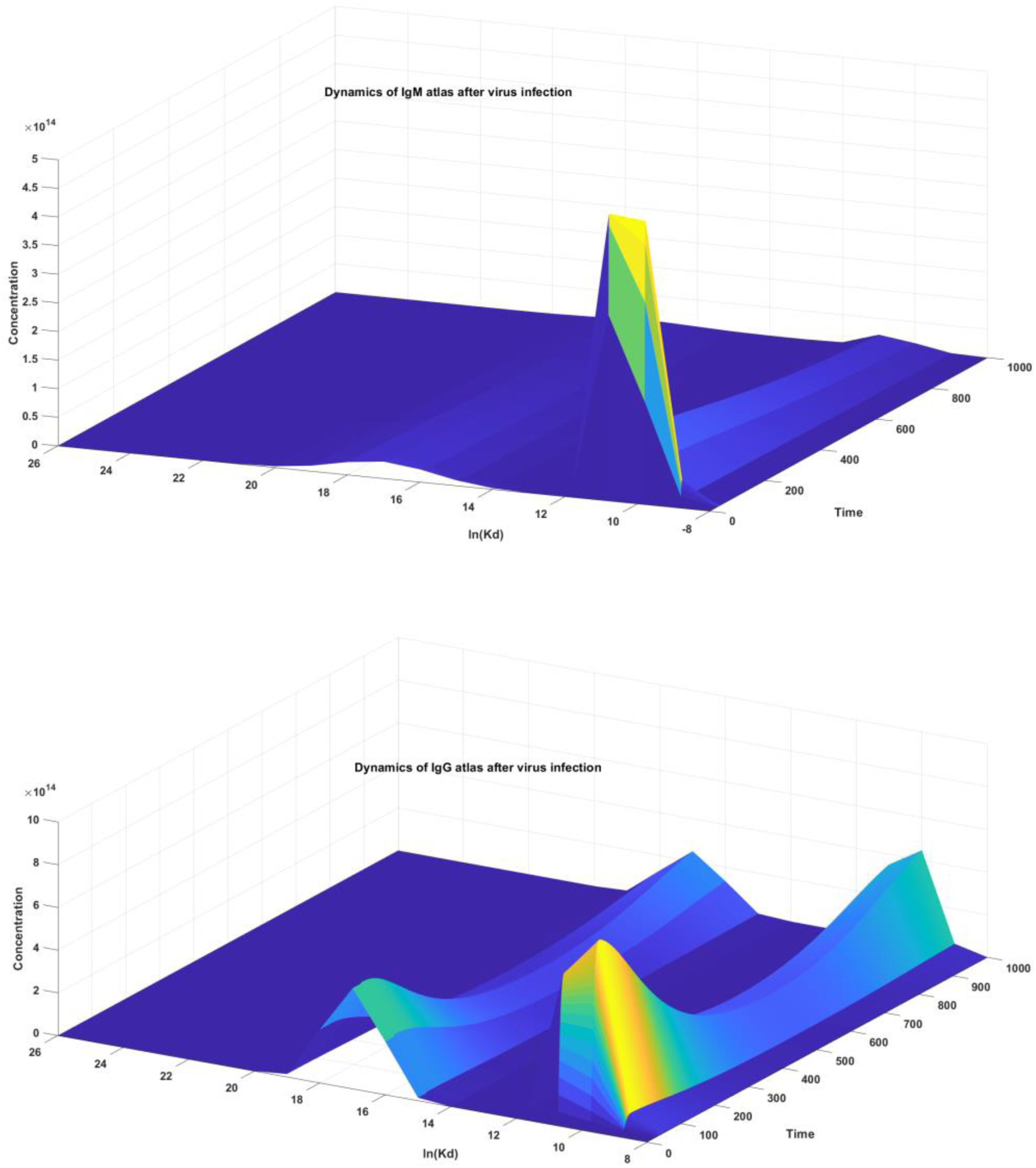
Antibody dynamics atlas of IgG and IgM after virus infection. The figure indicates that IgM undergoes proliferation earlier than IgG, but subsequently experiences rapid decay. Proliferation of IgG is significantly delayed, but its concentration declines more slowly after reaching peak levels and can be maintained at a higher concentration for an extended period.

If ELISA testing were employed to assess the levels of IgM and IgG, the kinetic curves during the initial infection would resemble those depicted in Figure 7c. IgG tends to stabilize at a relatively high level for an extended period, exhibiting certain fluctuations with the assistance of its own antigenic substances, whereas the concentration of IgM rapidly declines to a lower level. However, this scenario pertains to primary infections; for secondary infections caused by variant strains, the situation may differ. Due to the presence of immune imprinting effects, the initial high-binding IgG concentration in response to infection with variant strains does not plummet to an extremely low level. Consequently, during secondary infections with variant strains, IgG may undergo rapid elevation, with the magnitude and rate of increase potentially surpassing those of IgM. This phenomenon is particularly evident in the context of dengue virus infections, where clinical researchers often rely on the interplay between IgM and IgG to discern whether a patient is experiencing

**Figure 7c:**
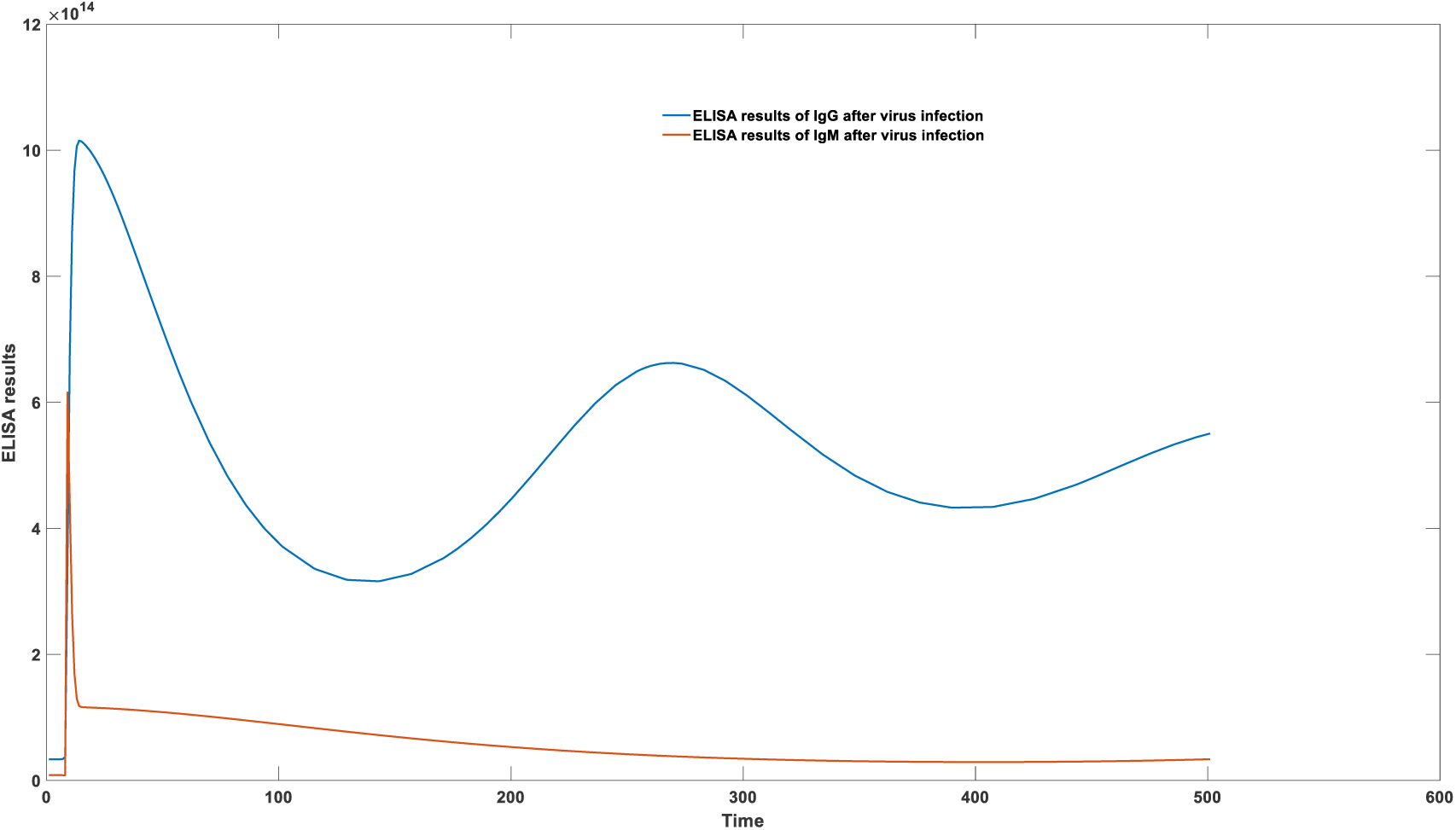
The ELISA test results of IgG and IgM after virus infection. From the figure, it is evident that IgG can maintain a relatively high concentration for an extended period, with occasional fluctuating increases. On the other hand, IgM exhibits a gradual declining trend. Therefore, the maintenance of persistent immunity is primarily achieved by IgG.

Two-dose vaccination involves administering two doses of a vaccine to achieve optimal immunity against a particular disease. This approach offers several advantages. Firstly, it ensures a robust immune response by priming the immune system with an initial dose and then reinforcing it with a booster dose, resulting in increased antibody levels and longer-lasting protection. Secondly, two doses allow for the development of memory B and T cells, which can provide rapid and effective immunity upon subsequent exposure to the pathogen. Additionally, two-dose vaccination strategies have been shown to enhance vaccine effectiveness, particularly against highly infectious or rapidly evolving pathogens, thus reducing the likelihood of breakthrough infections and contributing to herd immunity [64–65].

Here, we simulated the impact of a two-dose vaccination strategy on the elevation of IgM and IgG levels. As depicted in Figure 7d, upon initial vaccination, both IgM and IgG experience substantial increases, with IgM peaking earlier than IgG. Subsequently, IgM undergoes rapid decline, while IgG maintains a higher concentration level. Upon the second vaccination, IgG levels undergo further elevation, whereas IgM shows limited elevation and undergoes another round of decay, ultimately decreasing to a lower level. It is noteworthy that although the magnitude of IgM elevation is less pronounced compared to IgG, IgM still experiences a certain degree of elevation. This simulation underscores the significant role of sequential vaccination strategies in substantially boosting IgG levels, thereby enhancing vaccine efficacy.

**Figure 7d:**
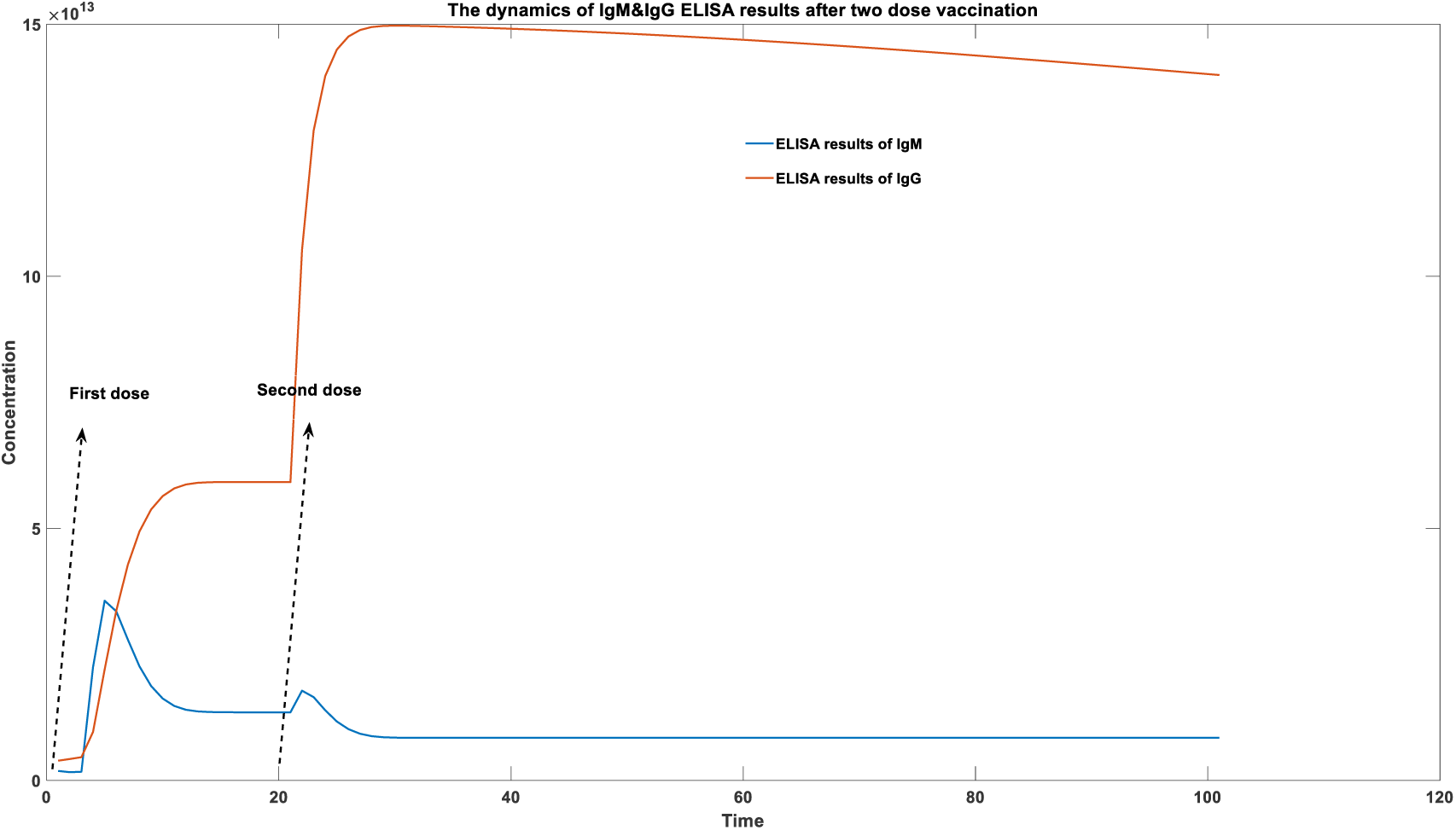
The dynamics of IgM and IgG after two dose vaccination. The figure demonstrates that a secondary vaccination can further elevate the concentration of IgG, while having a relatively minor impact on IgM.

## Discussion

The antibody response induced by viruses is a population-level change, not a proliferation response of specific antibodies. Viral infection reshapes the entire antibody atlas, leading not only to an increase in the proportion of corresponding high-affinity antibodies but also to a decrease in the proportion of antibodies with weak binding affinity to the viral antigen. Experimental data from antibody screening conducted based on binding activity using flow cytometry strongly support our simulation results [66–67]. Before viral infection, antibodies targeting viral antigens are densely distributed in regions with low binding activity, and this distribution is close to a normal distribution. However, following viral infection or vaccination, antibodies targeting viral antigens undergo differentiation and proliferation. A new peak appears in the high binding activity region, while the peak in the original low binding region tends to decrease.

The long-term maintenance of IgG in the immunological community is achieved through memory B cells. However, the maintenance of memory B cells has long been an unresolved mystery. Memory cells may not necessarily have a longer lifespan compared to ordinary B cells. Our model suggests that the maintenance of memory cells is a dynamic process, wherein, in the presence of self-antigens, the self-antibody repertoire is reshaped, allowing corresponding memory cells to be maintained at a stable level for an extended period. This maintenance effect of self-antigen-like substances on memory B cells manifests as fluctuations in IgG levels for a period following infection, a phenomenon corroborated by clinical data measuring long-term IgG antibody concentrations []. Additionally, our model indicates that the fundamental cause of secondary infection is viral mutation rather than a decrease in IgG levels following primary infection. For pathogens less prone to mutation, antibody reshaping resulting from natural infection or vaccination can sustain lifelong immunity.

Dynamic simulations indicate that while the immune system reshapes the antibody repertoire in two aspects—increasing binding coefficients and reducing dissociation coefficients—it notably favors augmenting the binding coefficient (Kon). Antibodies with higher (Kon) coefficients often demonstrate more efficient prevention of infection and viral clearance compared to those with better (Kd) coefficients. Selecting antibody sequences with high binding activity through experimental means can effectively realize molecular therapeutic interventions for infections. Many clinical antibody molecule selections are based on the equilibrium constant (Kd) of antibody-antigen binding [68–70]. However, our simulations reflect that during antibody selection, it is inadequate to simply prioritize antibodies with the best (Kd) coefficients; rather, antibodies with superior (Kon) coefficients should be selected through binding kinetics screening.

While flow cytometry can provide a more comprehensive reflection of changes in the antibody atlas, ELISA experiments remain the most convenient means for assessing specific antibody concentration levels, primarily due to cost and operational feasibility considerations. Our model computes ELISA results based on the mechanism of macromolecular binding. Our research indicates that ELISA results do not exhibit a linear relationship with the concentration of a specific antigen or with the overall antibody concentration; rather, they represent a comprehensive reflection of the interaction between the antibody atlas and the antigen. ELISA outcomes are related to changes in the overall composition of antibodies and cannot accurately reflect changes in the concentration of specific antibodies. Additionally, based on ELISA results, we can perform parameter fitting on the model to more intuitively demonstrate the dynamic changes in the antibody atlas.

From a dynamic perspective, our model effectively elucidates the origins of antigenic sin. In the absence of antibody-dependent enhancement (ADE), original antigenic sin can afford superior protection. However, due to antigenic sin, the proportion of highly active binding antibodies generated in response to variant strains is significantly smaller compared to the control group initially infected with variant strains. Our model reveals that while natural infection struggles to overcome the antibody generation impediment imposed by original antigenic sin, vaccination with variant strains can effectively mitigate immune imprinting.

Another crucial functionality of our model lies in its ability to systematically evaluate the virulence of viruses and the immune responses of hosts. The virulence of a virus post-infection primarily manifests in its replicative capacity. Host immunity is primarily reflected in two aspects: initial immunity, which pertains to the distribution of antibody binding strength against the viral antigen at the initial state, and dynamic immunity, which relates to the capacity for immune remodeling, mainly characterized by the ability of virus-antibody complexes to stimulate antibody production and the clearance rate of virus-antibody complexes, influenced by factors such as the activity of helper T cells and natural killer (NK) cells. Parameter fitting of different individuals or populations can reveal these differences, thus enabling better assessments of host and viral attributes. For instance, to investigate variations in the pathogenicity of different strains, it essentially involves identifying differences in the k1 value of the virus. By studying population infection data of different strains, including changes in viral levels within infected individuals and dynamic results from ELISA, we can fit this data to obtain differences in the proliferation coefficient (k1 value) of different strains, thereby determining variations in viral virulence. This method can also identify cases of defective virus infection within the same strain [71]. Moreover, if we aim to compare differences in individual immune response capabilities, we can fit the immunological coefficients of different individuals during infection with the same strain, primarily reflecting differences in k3 and k4 values. This enables the determination of individual immune characteristics against specific viruses, providing better insight into which demographics are more prone to developing severe illness. Future parameter fitting using real clinical data can better predict trends in antibody changes, while also elucidating differences in initial antibody distributions and antibody kinetic parameters among individuals. Such variations are closely associated with the severity of the disease.

The aftermath of viral infections can persist for a considerable duration, with the occurrence and duration of post-viral sequelae such as long COVID potentially correlated with interactions between antibody atlases and autoantigens. Parameter information obtained through fitting ELISA results enables further assessment of the probability of certain individuals experiencing such sequelae.

Our model further elucidates the differential distribution of IgG and IgM, wherein IgM, originating from initial B cell differentiation, exhibits significantly greater sequence diversity compared to IgG antibodies. Consequently, IgM presents lower overall concentrations and larger variance in antigen binding affinities, contrasting with IgG’s higher overall concentrations and smaller binding affinity variances. Thus, during initial infection, IgG often fails to rapidly proliferate high-affinity antibodies due to the initially low concentrations of antibodies in high-binding activity regions. Conversely, IgM can rapidly proliferate high-affinity antibodies. These antibodies undergo class switching to further transform into IgG, resulting in a rapid and significant decline in IgM levels post-infection. While IgG levels initially peak and gradually decline, this decline is not continuous, exhibiting periodic fluctuations thereafter. Experimental data strongly corroborate our mathematical simulations regarding the short-term and long-term dynamics of IgG and IgM. Future prospects may involve the artificial implantation of engineered monoclonal antibody B cells via targeted modifications [72–74], contingent upon favorable antibody kinetics and higher initial concentrations.

## Methods

### A simple mathematical model of one antibody-virus interaction

A simple mathematical representation of the immune response is described in the diagram below. In Fig. 8, x denotes the amount of antibody-antigen (virus) complex, y denotes the total number of antibodies, and z denotes the number of viruses. Process 1 represents the virus self-replication, characterized by a proliferation coefficient (k1). Process 2 signifies the virus-antibody binding process, with a forward reaction coefficient (k2) and a reverse dissociation coefficient (k-2). Process 3 denotes the positive feedback effect of virus-antibody complexes on antibody regeneration, governed by a feedback coefficient (k3). Process 4 illustrates the in vivo clearance of virus-antibody complexes, governed by a reaction coefficient (k4). Process 5 symbolizes the self-decay of antibodies, governed by a reaction coefficient (k5). Reaction 6 delineates the binding process between self-antigenic substances and antibodies, characterized by a forward reaction coefficient (k6) and a reverse dissociation coefficient (k-6). Reaction 7 delineates the positive feedback effect of self-antigen-antibody complexes on antibody regeneration, governed by a feedback coefficient (k7). Reaction 8 elucidates the clearance process of self-antigen-antibody complexes, governed by a coefficient (k8). Reaction 9 delineates the supplementary action of self-antigens, with a supplementation rate (C1).

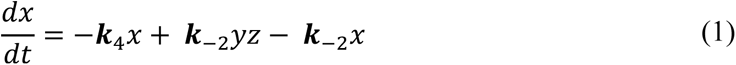

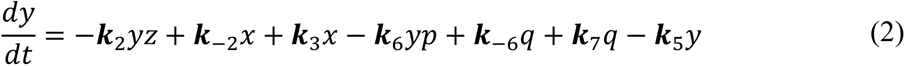

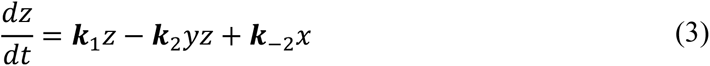

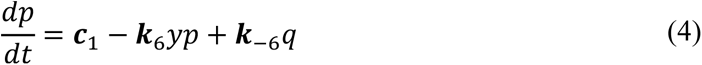

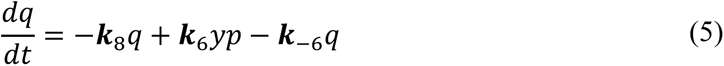

Based on the model described by Equations (1), (2), (3), it can be observed that the concentration of antibodies will eventually diminish to zero. This decay is attributed to the presence of a decay coefficient, denoted as ζ, which ensures that the antibodies fade away within a short time frame. However, in contrast to this observation, certain antibodies are capable of persisting in the human body for extended periods, offering lifelong protection. This phenomenon forms the fundamental basis for vaccine development. Immunologists commonly refer to these long-lasting B-cells or T-cells as “memory cells”. Although empirical studies have confirmed that these “memory cells” are distinct subclasses of immune cells (Inoue et al., 2018; Kurosaki et al., 2015), they exhibit similar half-lives to normal CD8+ cells (van den Berg et al., 2021). Consequently, the sustained antibody levels from “memory cells” can be attributed to a continuous stimulation triggering the proliferation of these memory cells. We propose that this stimulation arises from the presence of environmental antigen-like substances. These substances, which can be derived from various sources such as food, air, or even endogenous factors, exhibit weak binding affinity with neutralizing antibodies. Termed “environmental antigen-like substances”, they elicit weak signals upon presentation to T-cells, thereby promoting the proliferation of memory B-cells or T-cells. Due to their weak binding and the close similarity of protein sequences between these substances and our own body, their antigenicity is relatively low. Accordingly, the secreted stimulation signals by T-cells remain correspondingly weak. A dynamic equilibrium is established at a certain time point where the decay of memory cells is counterbalanced by the generation of new memory cells. A comprehensive elucidation of the virus-antibody interaction model can be found in our previous publication [13].

When vaccinated rather than naturally infected, as the virus lacks replicative activity, k1 becomes 0, accompanied by the natural decay coefficient Ω of the injected antigenic substance. Simultaneously, the initial viral antigen concentration increases to a relatively large value. The system of ordinary differential equations becomes:

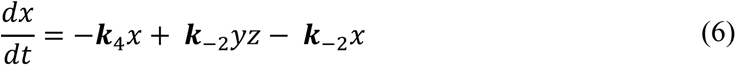

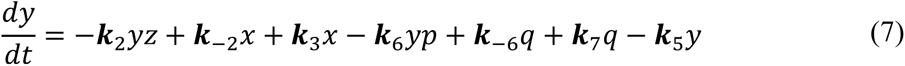

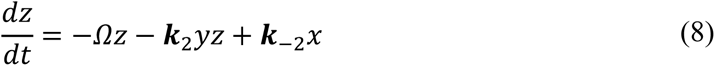

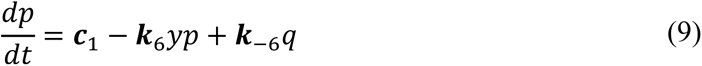

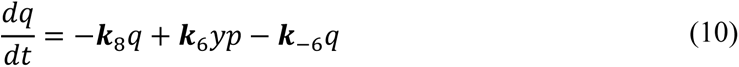

### A complex model involving the interplay between the antibody atlas and virus

For the simulation of an antibody atlas involving multiple types of antibodies, the interaction between each antibody and the virus follows the same dynamics as that of individual antibodies. The distinction lies in the unique association rate constants (Kon) and dissociation rate constants (K_off) for each antibody. Additionally, each antibody shares the same association and dissociation rate constants when interacting with self-antigens. The system of ordinary differential equations is represented as follows:

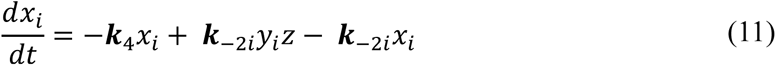

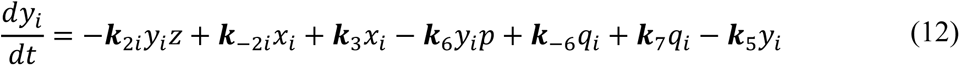

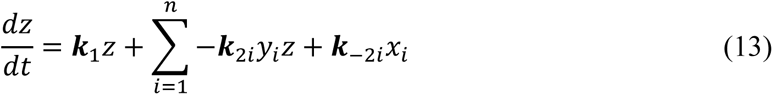

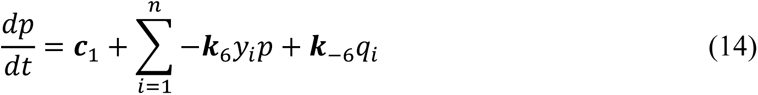

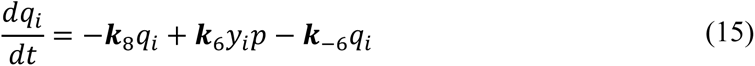

### Calculation of ELISA result

Since the ELISA test employs antigenic substances, denoted as (z), which neither degrade nor proliferate in the short term, and the ELISA test does not involve the clearance of antigen-antibody complexes within the body nor the regeneration of antibodies. Whereas (x) represents the overall ELISA results, i.e., the total concentration of complexes formed between various antibodies and antigens. The ordinary differential equations can be expressed as:

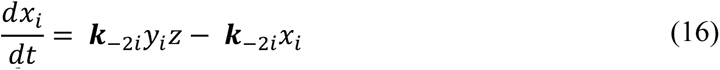

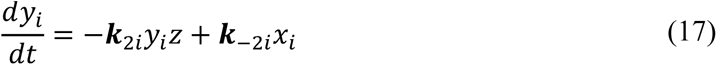

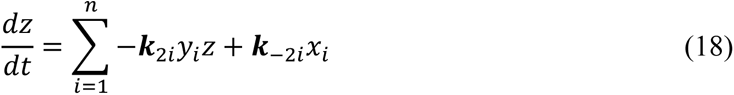

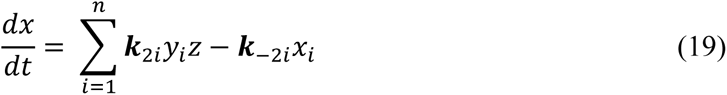

### The transformation of binding kinetics distribution when confronted with variant viral strains

Before introducing the transformation of the initial antibody distribution in response to different antigens, we need to introduce a concept known as the mutation coefficient (α). For two completely unrelated antigens, we consider the mutation coefficient to be positive infinity. For two identical antigens, we consider the mutation coefficient to be 1. We denote (f) as the reference normal distribution, (F) as the distribution of antibody binding parameters for a specific antigen, (N) as the total number of antibodies, and (F_New) as the new distribution of antibody binding parameters for the mutated antigen.

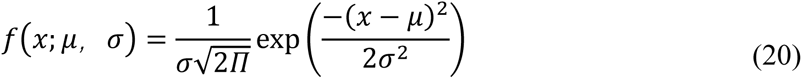

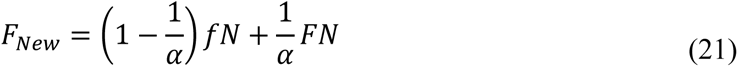

### Parameter estimation and sensitivity analysis

We did not perform parameter estimation based on existing clinical data. This is because our model requires the absolute concentration of antigenic substances and the total concentration of all antibodies in ELISA testing for accurate parameter fitting. Although these two pieces of data are relatively easy to measure, they are often not included in clinical databases. Traditional mathematical models do not necessitate the values of these two parameters. The primary objective of our research is to promote our model; therefore, we emphasize that using this model can effectively determine various coefficients. The fitting of the first type of coefficient, namely the parameters of initial antibody binding characteristics, can be achieved through fitting estimation using pre-infection ELISA data. The second type of coefficients encompasses two aspects: host-related coefficients, primarily including the proliferation coefficient of virus-antibody complexes and the in vivo clearance rate parameter of virus complexes, and virus-related coefficients, mainly the proliferation coefficient of the virus. Fitting a single parameter is relatively straightforward; we primarily focus on the dual-parameter fitting of host-related coefficients, which can be obtained through Markov Chain Monte Carlo (MCMC) methods. Our experimental data originate from simulated data based on assumed parameters, mainly including two aspects: the dynamic changes in virus titers and the dynamic changes in ELISA results. Here, there are only two parameters: (k3) and (k4). Since there are many compartments in the model, a single simulation requires considerable time. We used 10,000 simulations to calculate the posterior distribution of parameters. The likelihood for a single subject is:

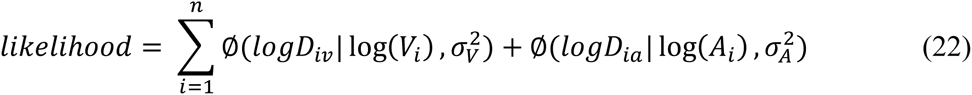

Here Φ is the probability density function of the normal distribution. The number of observations for a single individual is denoted by *n*, *D*iv is the *i*th viral titre measurement and *v*i is the model prediction of viral titre at the *i*th measurement. *σ*v is the error of viral titre measurements on a log10 scale, assumed to be 1. In addition, *D*ia is the *i*th antibody level measurement, *A*i is the model prediction of antibody level at the *i*th measurement, *σ*A is the error in antibody measurements assumed to be 1. Metropolis-Hastings Algorithm is applied to calculate the poster parameter distribution. The models were fitted using Markov Chain Monte Carlo (MCMC) methods using Matlab.

Extended Figure 1: Dynamic changes of all antibodies (100 types) in response to viral infection in the simulation. The binding parameters and dissociation constants for various antibodies can be referenced in Table S.11 in the supplementary materials.

Extended Figure 2: Dynamic changes of all antibodies (100 types) after viral infection in the absence of self-antigen.

Extended Figure 3: Differential kinetic changes of all antibodies (100 types) in primary infection (absence of immune imprinting) compared to secondary infection caused by variant strains (presence of immune imprinting).

Extended Figure 4: Dynamic changes of all IgM (100 types) and IgG (100 types) following viral infection in the simulation.

Extended Video 1: Dynamic changes of the forward binding coefficient ln(Kon) of the antibody repertoire following viral infection in the simulation.

Extended Video 2: Dynamic changes of the reverse dissociation coefficient ln(Koff) of the antibody repertoire following viral infection in the simulation.

Extended Video 3: Dynamic changes of the antigen equilibrium coefficient ln(Kd) of the antibody repertoire following viral infection in the simulation.

Extended Video 4: Dynamic changes of the antigen equilibrium coefficient ln(Kd) of the antibody repertoire following viral infection, comparing the presence and absence of self-antigen. The solid blue line represents the dynamic changes of ln(Kd) in the presence of self-antigen, while the solid red line represents the dynamic changes of ln(Kd) in the absence of self-antigen.

Extended Video 5: Dynamic changes of the antigen equilibrium coefficient ln(Kd) of the antibody repertoire comparing primary infection (absence of immune imprinting) with secondary infection caused by variant strains (presence of immune imprinting). The solid blue line represents the dynamic changes of ln(Kd) in the absence of immune imprinting, while the solid red line represents the dynamic changes of ln(Kd) in the presence of immune imprinting.

Extended Video 6: Dynamic changes of the antigen equilibrium coefficient ln(Kd) of IgM and IgG following viral infection in the simulation. The solid blue line represents the dynamic changes of ln(Kd) for IgG, while the solid red line represents the dynamic changes of ln(Kd) for IgM.

## Codes availability

Matlab source codes are available at https://github.com/zhaobinxu23/antibody_atlas_modeling

## Author Contributions

Conceptualization, Z.X.; methodology, Z.X. and J.X; writing—original draft preparation, Z.X.; writing—review and editing, Q.Z., D.W., J.S. and H.Z.; funding acquisition, Z.X. and Q.Z. All authors have read and agreed to the published version of the manuscript.

## Acknowledgments

We thank Dr Zuyi Huang from Villanova University. Yushan Zhu from Tsinghua University, Jian Song, Peiyan Guan, XiangYong Li, and Zhenlin Wei from Dezhou University for helpful conversations, comments, and clarifications.

---

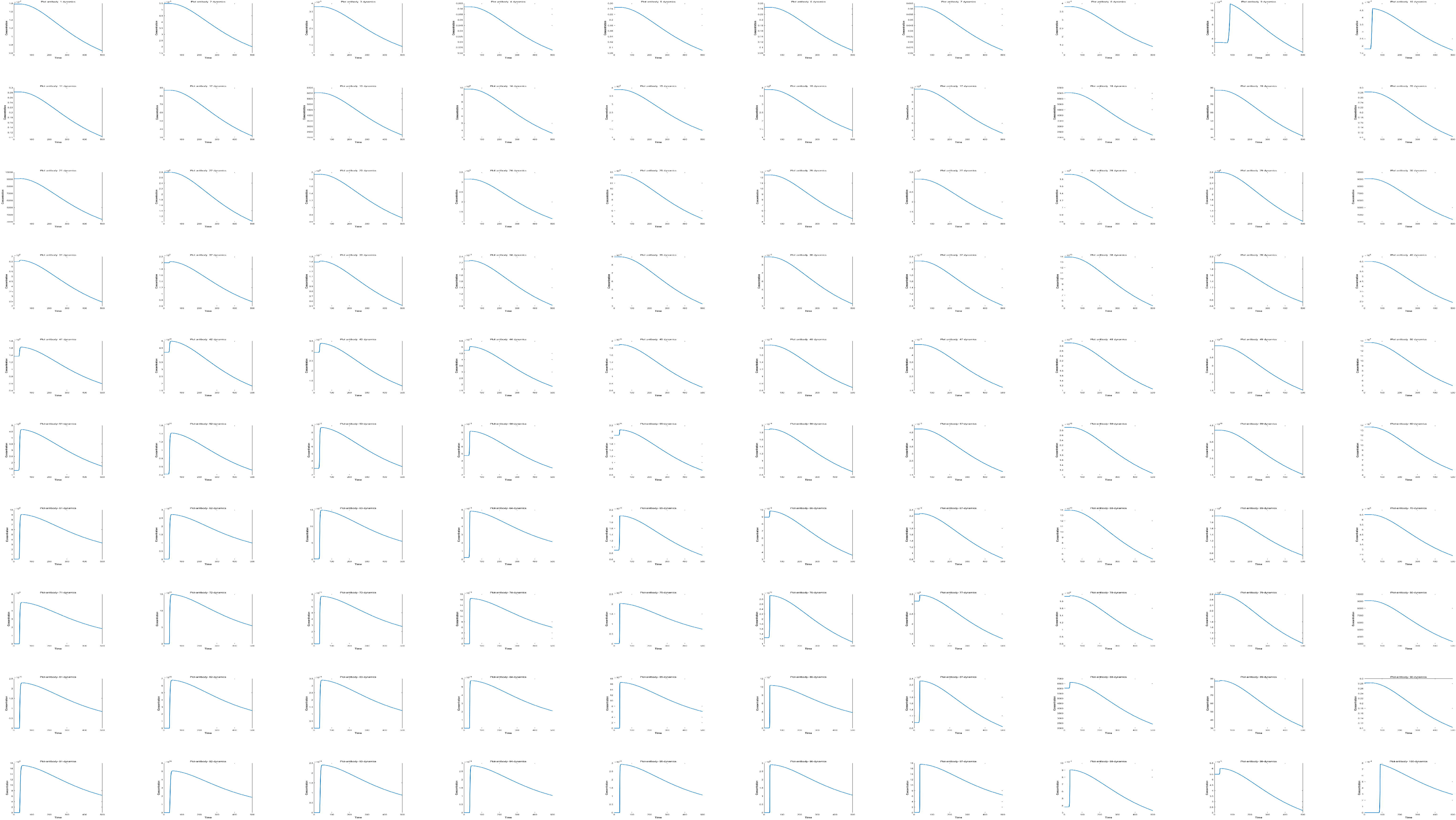
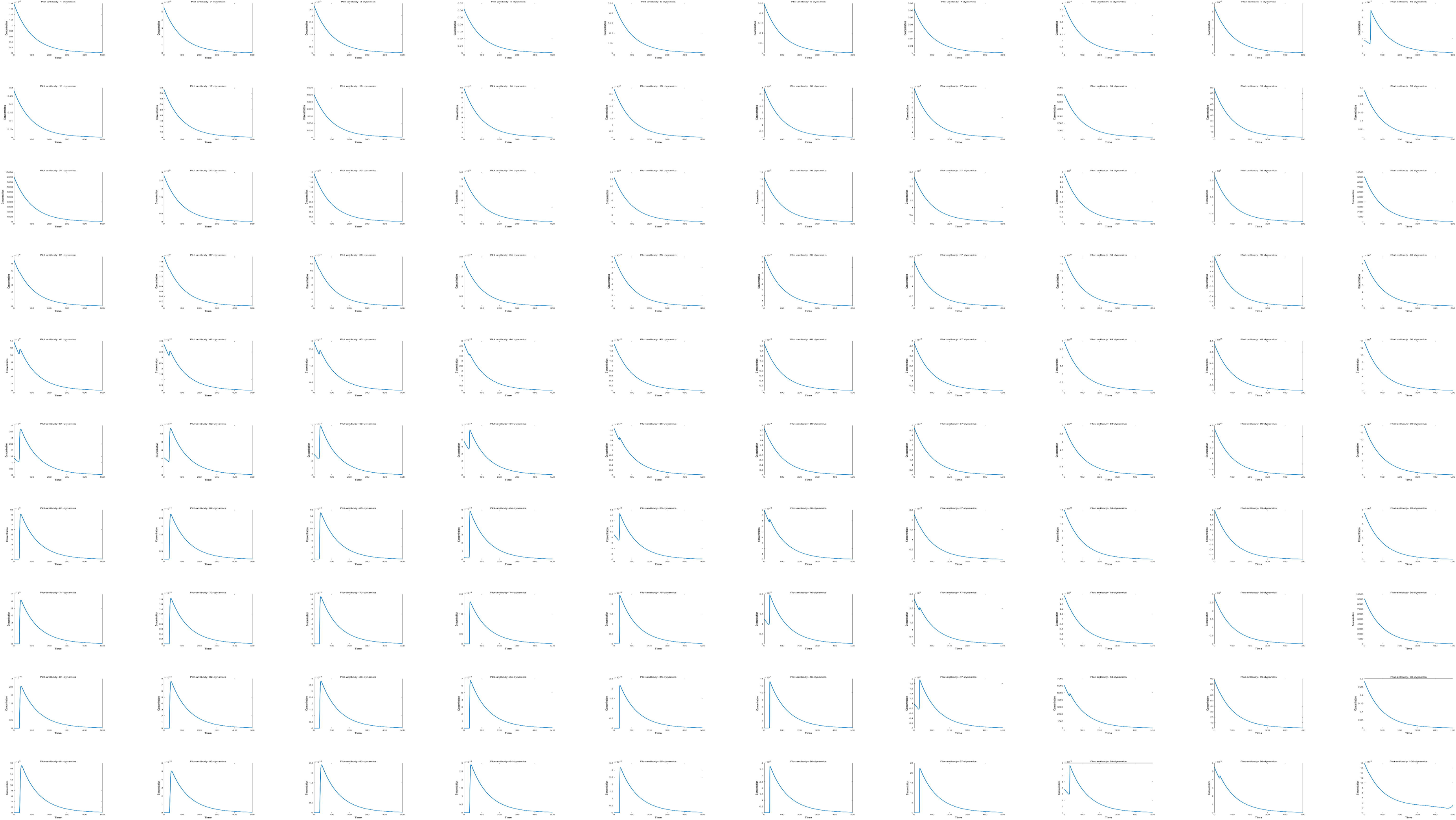
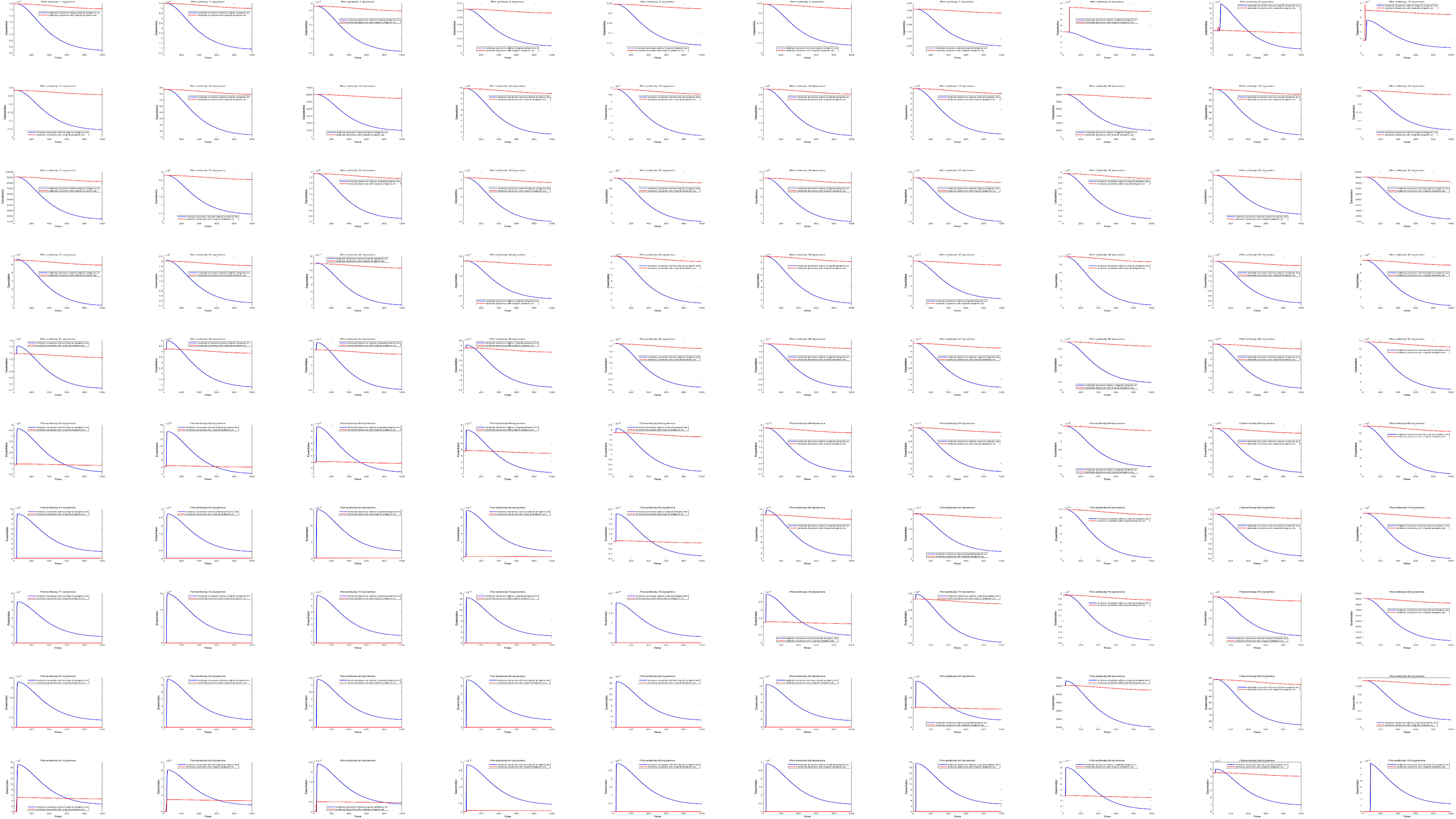
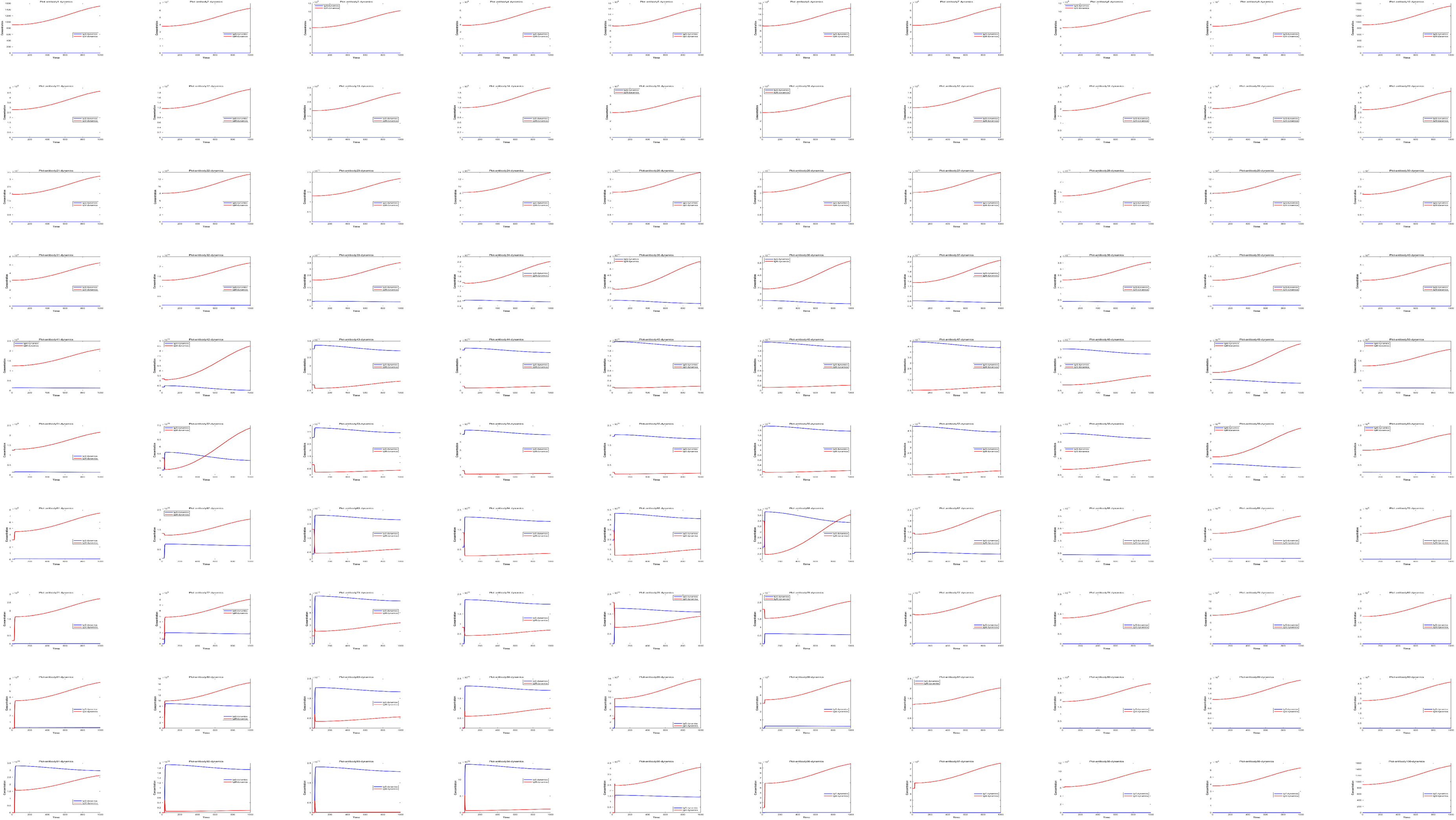

