## supplementary_materials for "Dynamic Modeling of Antibody Repertoire Reshaping in Response to Viral Infections"

### Unit selection and the distribution of Kon and Koff of experimental data

J. P. Landry et al. measured affinity constants of 1,450 monoclonal antibodies to peptide targets utilizing a microarray-based label-free assay platform [1]. Specifically, they measured the forward association rate constant (Kon) and the reverse dissociation rate constant (Koff) of these antibodies with peptides. The converted data in new units were plotted in Supplementary Figure 1 and Supplementary Figure 2.

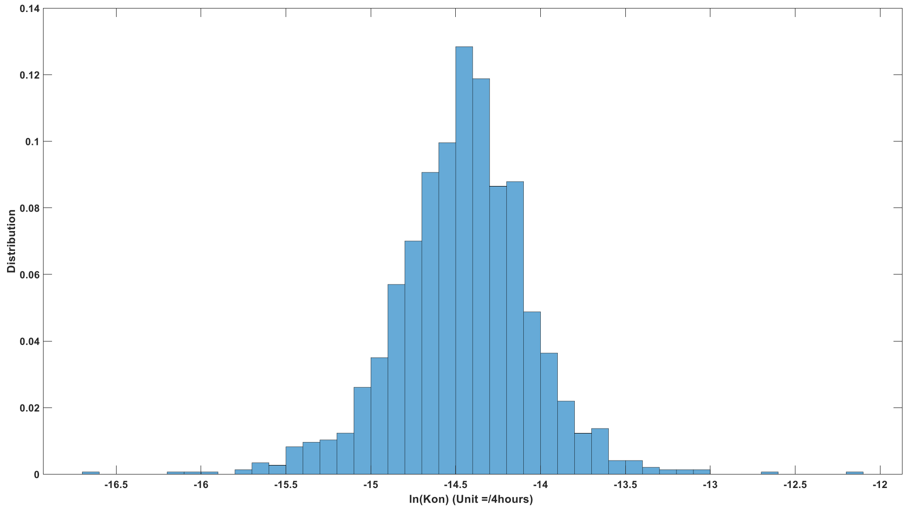


**Figure S1: ln（Kon）distribution of monoclonal antibodies to target peptide.**

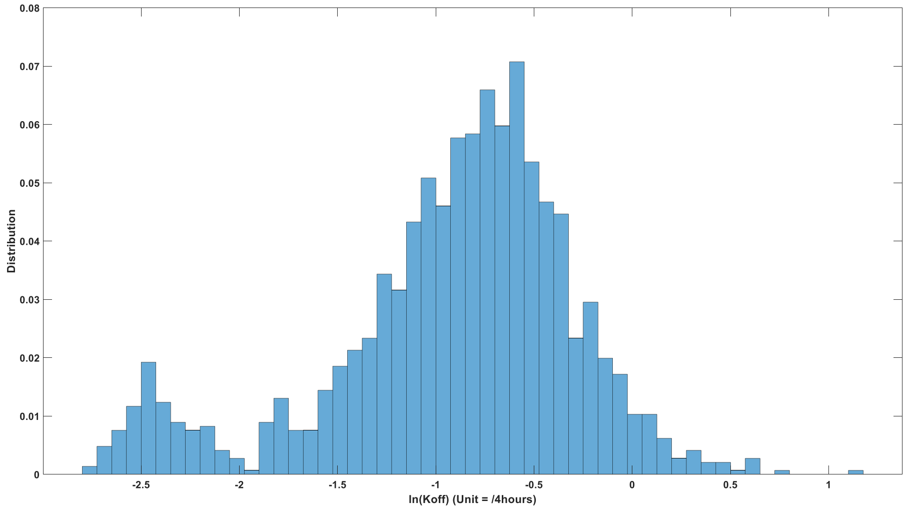


**Figure S2: ln（Koff）distribution of monoclonal antibodies to target peptide.**

The unit selected for ln(Kon) and ln(Koff) is 4h-1. While many parameters in our subsequent models are assumed, for the purpose of accurately reflecting the actual antibody generation process, we aim for the parameters to match the experimental data as closely as possible, particularly in terms of the antibody atlas distribution.

The average values of ln(Kon) and ln(Koff) are crucial. We use an average ln(Kon) of -15.5 and an average ln(Koff) of 1.5. Supplementary Figure 1 indicates an average ln(Kon) of -14.5, which is evidently larger than the result of a true random match, as it represents the binding affinity between monoclonal antibodies and their respective target peptides. Based on this value, we adjust the average ln(Kon) of the entire antibody library matched with random antigens to -15.5. Supplementary Figure 2 shows an average ln(Koff) around -1, which is smaller than the result of a true random match, as it represents the dissociation rate between monoclonal antibodies and their corresponding target peptides. Referencing this value, we increase the average ln(Koff) of the entire antibody library matched with random antigens to 1.5.

**The relationship between self-antigen concentration and its antibody binding affinity**

The set of ordinary differential equations for the simple model is as follows:

$$\frac{dx}{dt} = -\boldsymbol{k}_4 x + \boldsymbol{k}_{-2} yz - \boldsymbol{k}_{-2} x \tag{1}$$

$$\frac{dy}{dt} = -\boldsymbol{k}_2 yz + \boldsymbol{k}_{-2} x + \boldsymbol{k}_3 x - \boldsymbol{k}_6 yp + \boldsymbol{k}_{-6} q + \boldsymbol{k}_7 q - \boldsymbol{k}_5 y \tag{2}$$

$$\frac{dz}{dt} = \boldsymbol{k}_1 z - \boldsymbol{k}_2 yz + \boldsymbol{k}_{-2} x \tag{3}$$

$$\frac{dp}{dt} = \boldsymbol{c}_1 - \boldsymbol{k}_6 yp + \boldsymbol{k}_{-6} q \tag{4}$$

$$\frac{dq}{dt} = -\boldsymbol{k}_8 q + \boldsymbol{k}_6 yp - \boldsymbol{k}_{-6} q \tag{5}$$

Before viral invasion, antibodies exist in an equilibrium state, hence satisfying a quasi-steady state condition where the concentration of the virus-antibody complex (x) is 0, and the concentration of the virus (z) is 0. The system of equations is as follows:

$$\frac{dy}{dt} = -\boldsymbol{k}_6 yp + \boldsymbol{k}_{-6} q + \boldsymbol{k}_7 q - \boldsymbol{k}_5 y = 0 \tag{6}$$

$$\frac{dp}{dt} = \boldsymbol{c}_1 - \boldsymbol{k}_6 yp + \boldsymbol{k}_{-6} q = 0 \tag{7}$$

$$\frac{dq}{dt} = -\boldsymbol{k}_8 q + \boldsymbol{k}_6 yp - \boldsymbol{k}_{-6} q = 0 \tag{8}$$

The initial value of the antibody concentration, $y_0$, is known. The variable parameters include the initial concentration of self-antigenic substances, $p_0$, the initial concentration of the complex formed by antibodies and self-antigenic substances, $q_0$, the affinity coefficient, $k_6$, and dissociation coefficient, $k_{-6}$, of self-antigenic substances, as well as the complement factor, c1, of self-antigens. These five variable parameters satisfy equations 6-8. Therefore, by arbitrarily determining the initial values of two parameters, the numerical values of the other three parameters can be calculated to satisfy equations 6-8.

**Mathematical modeling of virus-antibody interaction (Model 1)**

Model 1.1 is utilized to simulate the self-antigen's role in maintaining antibody interactions, focusing solely on the kinetic characteristics of individual antibodies. Model 1.2 is employed to simulate the varying proliferative potentials of antibodies (four types) with different binding kinetic constants during the viral infection process. Model 1.3 is designed to simulate the impact of antibodies with two distinct forward binding coefficients on the dynamics of viruses and virus-antibody complex formation. Model 1.4 elucidates that antibodies with superior Kon coefficients are more effective in inhibiting virus proliferation and controlling the severity of infection compared to those with superior Kd coefficients.

| Reaction index | Reaction |
|---|---|
| 1 | virus → virus |
| 2 | Antibody + virus ⇄ Antibody-virus Complex |
| 3 | Antibody-virus Complex → Antibody |
| 4 | Antibody-virus Complex → Cleared by immune system |
| 5 | Antibody → Degradation |
| 6 | Antibody + self-antigen ⇄ Antibody-self-antigen Complex |
| 7 | Antibody-self-antigen Complex → Antibody |
| 8 | Antibody-self-antigen Complex → Cleared by immune system |
| 9 | Replenish from environment → self-antigen |

Table S1: Reaction index and the name of each reaction in model 1

| Parameter Name | Description | Value |
|---|---|---|
| $k_1$ | Virus proliferation rate | 1 |
| $k_2$ | Forward binding constant between antibody and virus | $10^{-12}$ |
| $k_{-2}$ | Dissociation constant of antibody-virus complex | 0.01 |
| $k_3$ | Feedback constant of antibody-virus complex on the regeneration of antibody | 5 |
| $k_4$ | Clearance rate of antibody-virus complex | 0.5 |
| $k_5$ | Antibody degradation rate | 0.01 |
| $k_6$ | Forward binding constant between antibody and self-antigen | k5*(k7+k4) /k4/$10^{12}$ |
| $k_7$ | Dissociation constant of antibody-self-antigen complex | 10 |
| $k_{-7}$ | Feedback constant of antibody-self-antigen complex on the regeneration of antibody | 1 |
| $k_8$ | Clearance rate of antibody-self-antigen complex | 0.5 |
| $C_1$ | Replenish rate of self-antigen | 2*$10^9$ |

Table S2：Parameter set in model 1.1.

| Variables | Meaning | Initial value |
|---|---|---|
| z | virus | 10 |
| y | Antibody | 2*$10^{11}$ |
| x | Antibody-virus complex | 0 |
| p | Self-antigen | $10^{12}$ |
| q | Antibody-self-antigen complex | y*k5/(k1-k4) |

Table S3：Time-dependent variables of model 1.1

The set of differential equations of model 1.1 is represented below:

$$\frac{dx}{dt} = -\boldsymbol{k}_4 x + \boldsymbol{k}_{-2} yz - \boldsymbol{k}_{-2} x \quad (1)$$

$$\frac{dy}{dt} = -\boldsymbol{k}_2 yz + \boldsymbol{k}_{-2} x + \boldsymbol{k}_3 x - \boldsymbol{k}_6 yp + \boldsymbol{k}_{-6} q + \boldsymbol{k}_7 q - \boldsymbol{k}_5 y \quad (2)$$

$$\frac{dz}{dt} = \boldsymbol{k}_1 z - \boldsymbol{k}_2 yz + \boldsymbol{k}_{-2} x \quad (3)$$

$$\frac{dp}{dt} = \boldsymbol{c}_1 - \boldsymbol{k}_6 yp + \boldsymbol{k}_{-6} q \quad (4)$$

$$\frac{dq}{dt} = -\boldsymbol{k}_8 q + \boldsymbol{k}_6 yp - \boldsymbol{k}_{-6} q \quad (5)$$

| Parameter Name | Description | Value |
|---|---|---|
| $k_1$ | Virus proliferation rate | 1 |
| $k_2$(antibody-1) | Forward binding constant between antibody-1 and virus | $10^{-12}$ |
| $k_{-2}$(antibody-1) | Dissociation constant of antibody-1-virus complex | 0.1 |
| $k_2$(antibody-2) | Forward binding constant between antibody-2 and virus | $10^{-12}$ |
| $k_{-2}$(antibody-2) | Dissociation constant of antibody-2-virus complex | 1 |
| $k_2$(antibody-3) | Forward binding constant between antibody-3 and virus | $9*10^{-11}$ |
| $k_{-2}$(antibody-3) | Dissociation constant of antibody-3-virus complex | 0.09 |
| $k_2$(antibody-4) | Forward binding constant between antibody-4 and virus | $8*10^{-11}$ |
| $k_{-2}$(antibody-4) | Dissociation constant of antibody-4-virus complex | 0.08 |
| $k_3$ | Feedback constant of antibody-virus complex on the regeneration of antibody | 5 |
| $k_4$ | Clearance rate of antibody-virus complex | 0.5 |
| $k_5$ | Antibody degradation rate | 0.01 |
| $k_6$ | Forward binding constant between antibody and self-antigen | $k_5*(k_7+k_4)/k_4/10^{12}$ |
| $k_7$ | Dissociation constant of antibody-self-antigen complex | 10 |
| $k_{-7}$ | Feedback constant of antibody-self-antigen complex on the | 1 |

| | | |
|---|---|---|
| | regeneration of antibody | |
| $k_8$ | Clearance rate of antibody-self-antigen complex | 0.5 |
| $C_1$ | Replenish rate of self-antigen | $4*10^3$ |

Table S4: Parameter set in model 1.2.

| Variables | Meaning | Initial value |
|---|---|---|
| z | virus | 100 |
| y1 | Antibody-1 | $10^5$ |
| y2 | Antibody-2 | $10^5$ |
| y3 | Antibody-3 | $10^5$ |
| y4 | Antibody-4 | $10^5$ |
| x1 | Antibody-1-virus complex | 0 |
| x2 | Antibody-2-virus complex | 0 |
| x3 | Antibody-3-virus complex | 0 |
| x4 | Antibody-4-virus complex | 0 |
| p | Self-antigen | $10^{12}$ |
| q1 | Antibody-1-self-antigen complex | y1*k5/(k1-k4) |
| q2 | Antibody-2-self-antigen complex | y2* k5/(k1-k4) |
| q3 | Antibody-3-self-antigen complex | y3* k5/(k1-k4) |
| q4 | Antibody-4-self-antigen complex | y4* k5/(k1-k4) |

Table S5: Time-dependent variables of model 1.2

The set of differential equations of model 1.2-1.4 is represented below:

$$\frac{dx_i}{dt} = -\boldsymbol{k}_4 x_i + \boldsymbol{k}_{-2i} y_i z - \boldsymbol{k}_{-2i} x_i \qquad (6)$$

$$\frac{dy_i}{dt} = -\boldsymbol{k}_{2i} y_i z + \boldsymbol{k}_{-2i} x_i + \boldsymbol{k}_3 x_i - \boldsymbol{k}_6 y_i p + \boldsymbol{k}_{-6} q_i + \boldsymbol{k}_7 q_i - \boldsymbol{k}_5 y_i \qquad (7)$$

$$\frac{dz}{dt} = \boldsymbol{k}_1 z + \sum_{i=1}^{n} -\boldsymbol{k}_{2i} y_i z + \boldsymbol{k}_{-2i} x_i \quad (8)$$

$$\frac{dp}{dt} = \boldsymbol{c}_1 + \sum_{i=1}^{n} -\boldsymbol{k}_6 y_i p + \boldsymbol{k}_{-6} q_i \quad (9)$$

$$\frac{dq_i}{dt} = -\boldsymbol{k}_8 q_i + \boldsymbol{k}_6 y_i p - \boldsymbol{k}_{-6} q_i \quad (10)$$

| Parameter Name | Description | Value |
|---|---|---|
| $k_1$ | Virus proliferation rate | 1 |
| $k_2$(antibody-1) | Forward binding constant between antibody-1 and virus | $2*10^{-15}$ |
| $k_{-2}$(antibody-1) | Dissociation constant of antibody-1-virus complex | 10 |
| $k_2$(antibody-2) | Forward binding constant between antibody-2 and virus | $10^{-15}$ |
| $k_{-2}$(antibody-2) | Dissociation constant of antibody-2-virus complex | 10 |
| $k_3$ | Feedback constant of antibody-virus complex on the regeneration of antibody | 5 |
| $k_4$ | Clearance rate of antibody-virus complex | 0.5 |
| $k_5$ | Antibody degradation rate | 0 |
| $k_6$ | Forward binding constant between antibody and self-antigen | 0 |
| $k_7$ | Dissociation constant of antibody-self-antigen complex | 0 |
| $k_{-7}$ | Feedback constant of antibody-self-antigen complex on the regeneration of antibody | 1 |
| $k_8$ | Clearance rate of antibody-self-antigen complex | 0.5 |
| $C_1$ | Replenish rate of self-antigen | 0 |

Table S6：Parameter set in model 1.3.

| Variables | Meaning | Initial value |
|---|---|---|
| z | virus | 100 |
| y1 | Antibody-1 | $10^{14}$ |

| | | |
|---|---|---|
| y2 | Antibody-2 | $10^{14}$ |
| x1 | Antibody-1-virus complex | 0 |
| x2 | Antibody-2-virus complex | 0 |
| p | Self-antigen | 0 |
| q1 | Antibody-1-self-antigen complex | 0 |
| q2 | Antibody-2-self-antigen complex | 0 |

Table S7：Time-dependent variables of model 1.3

| Parameter Name | Description | Value |
|---|---|---|
| $k_1$ | Virus proliferation rate | 1 |
| $k_2$(antibody-1) | Forward binding constant between antibody-1 and virus | $10^{-12}$ |
| $k_{-2}$(antibody-1) | Dissociation constant of antibody-1-virus complex | 0.1 |
| $k_2$(antibody-2) | Forward binding constant between antibody-2 and virus | $2*10^{-12}$ |
| $k_{-2}$(antibody-2) | Dissociation constant of antibody-2-virus complex | 1 |
| $k_3$ | Feedback constant of antibody-virus complex on the regeneration of antibody | 5 |
| $k_4$ | Clearance rate of antibody-virus complex | 0.5 |
| $k_5$ | Antibody degradation rate | 0.01 |
| $k_6$ | Forward binding constant between antibody and self-antigen | $k_5*(k_7+k_4)/k_4/10^{12}$ ($10^{12}$ = initial value of p) |
| $k_7$ | Dissociation constant of antibody-self-antigen complex | 10 |
| $k_{-7}$ | Feedback constant of antibody-self-antigen complex on the regeneration of antibody | 1 |
| $k_8$ | Clearance rate of antibody-self-antigen complex | 0.5 |
| $C_1$ | Replenish rate of self-antigen | $2*10^9$ |

Table S8：Parameter set in model 1.4.

| Variables | Meaning | Initial value |
|---|---|---|
| z | virus | 10 |
| y1 | Antibody-1 | $2*10^{11}$ |
| y2 | Antibody-2 | $2*10^{11}$ |
| x1 | Antibody-1-virus complex | 0 |
| x2 | Antibody-2-virus complex | 0 |
| p | Self-antigen | $10^{12}$ |
| q1 | Antibody-1-self-antigen complex | $y1*k_5/(k_1-k_4)$ |
| q2 | Antibody-2-self-antigen complex | $y2*k_5/(k_1-k_4)$ |

Table S9：Time-dependent variables of model 1.4

**Mathematical modeling of antibody atlas dynamics (Model 2)**

Model 2.1 represents the dynamic changes of antibody atlas in the absence of viral infection, emphasizing the maintenance role of self-antigens on the antibody repertoire, with an initial viral concentration of 0. Model 2.2 depicts the dynamic changes of antibody atlas in the presence of viral infection, while also simulating the dynamics of the antibody repertoire following viral infection in the absence of self-antigens, with an initial viral concentration of 100. Model 2.3 compares the characteristic changes in the antibody atlas between primary infection and the presence of immune imprinting. Model 2.4 illustrates that vaccination helps eliminate the antibody generation barriers caused by immune imprinting, with a replication coefficient k1 for the virus set to -0.02(decay constant of virus antigen in the vaccine), while the initial antigen concentration is set to a relatively high value.

| Parameter Name | Description | Value |
|---|---|---|
| $k_1$ | Virus proliferation rate | 1 |
| $k_2$ (antibody-M) (M = 10*(i-1) + j) | Forward binding constant between antibody-1 and virus | $Kon_i$ |
| $k_{-2}$(antibody-M) | Dissociation constant of antibody-1-virus complex | $Koff_i$ |
| $k_3$ | Feedback constant of antibody-virus complex on the | 5 |

| | | |
|---|---|---|
| | regeneration of antibody | |
| $k_4$ | Clearance rate of antibody-virus complex | 0.5 |
| $k_5$ | Antibody degradation rate | 0.01 |
| $k_6$ | Forward binding constant between antibody and self-antigen | $k_5*(k_7+k_4)/k_4/10^{16}$ ($10^{16}$ = initial value of p) |
| $k_7$ | Dissociation constant of antibody-self-antigen complex | 10 |
| $k_{-7}$ | Feedback constant of antibody-self-antigen complex on the regeneration of antibody | 1 |
| $k_8$ | Clearance rate of antibody-self-antigen complex | 0.5 |
| $C_1$ | Replenish rate of self-antigen | $10^{13}$ ($10^{13}$ = initial overall antibody concentration $*k_5$) This value equal to 0 in the absence of self-antigen |

Table S10：Parameter set in model 2.1 and model 2.2.

| Parameter Name | Value |
|---|---|
| $Kon_1$ | $10^{-20}$ |
| $Kon_2$ | $10^{-19}$ |
| $Kon_3$ | $10^{-18}$ |
| $Kon_4$ | $10^{-17}$ |
| $Kon_5$ | $10^{-16}$ |
| $Kon_6$ | $10^{-15}$ |
| $Kon_7$ | $10^{-14}$ |
| $Kon_8$ | $10^{-13}$ |

| | |
|---|---|
| $Kon_9$ | $10^{-12}$ |
| $Kon_{10}$ | $10^{-11}$ |
| $Koff_1$ | $10^{-3}$ |
| $Koff_2$ | $10^{-2}$ |
| $Koff_3$ | $10^{-1}$ |
| $Koff_4$ | 1 |
| $Koff_5$ | 10 |
| $Koff_6$ | $10^2$ |
| $Koff_7$ | $10^3$ |
| $Koff_8$ | $10^4$ |
| $Koff_9$ | $10^5$ |
| $Koff_{10}$ | $10^6$ |

Table S11：Parameter set of Kon and Koff.

| Variables | Meaning | Initial value |
|---|---|---|
| z | virus | 0 in model 2.1, 100 in model 2.2. |
| $y_M$ (M = 10*(i-1) + j) | Antibody M | Prob_A(i)*Prob_B(j)*overall_initial_antibody (overall_initial_antibody = $10^{15}$) |
| $x_M$ | Antibody-M-virus complex | 0 |
| p | Self-antigen | $10^{16}$ (this value equal to zero in the absence of self-antigen) |
| $q_M$ | Antibody-M-self-antigen complex | $y_M*k_5/(k_1-k_4)$ (this value equal to zero in the absence of self-antigen) |

Table S12：Time-dependent variables of model 2.1 and model 2.2.

| Parameter Name | Value |
|---|---|

| | |
|---|---|
| Prob_A(1) | P(X ≤ -19.5) |
| Prob_A(2) | P(-19.5< X ≤ -18.5) |
| Prob_A(3) | P(-18.5< X ≤ -17.5) |
| Prob_A(4) | P(-17.5< X ≤ -16.5) |
| Prob_A(5) | P(-16.5< X ≤ -15.5) |
| Prob_A(6) | P(-15.5< X ≤ -14.5) |
| Prob_A(7) | P(-14.5< X ≤ -13.5) |
| Prob_A(8) | P(-13.5< X ≤ -12.5) |
| Prob_A(9) | P(-12.5< X ≤ -11.5) |
| Prob_A(10) | P(-11.5< X) |
| Prob_B(1) | P(x ≤ -2.5) |
| Prob_B(2) | P(-2.5< x ≤ -1.5) |
| Prob_B(3) | P(-1.5< x ≤ -0.5) |
| Prob_B(4) | P(-0.5< x ≤ 0.5) |
| Prob_B(5) | P(0.5< x ≤ 1.5) |
| Prob_B(6) | P(1.5< x ≤ 2.5) |
| Prob_B(7) | P(2.5< x ≤ 3.5) |
| Prob_B(8) | P(3.5< x ≤ 4.5) |
| Prob_B(9) | P(4.5< x ≤ 5.5) |
| Prob_B(10) | P(5.5< x) |

Table S13： Caculation of Prob_A and Prob_B. x follows normal distribution N(1.5,0.8), X follows normal distribution N(-15.5,0.5).

| Parameter Name | Description | Value |
|---|---|---|

| | | |
|---|---|---|
| $k_1$ | Virus proliferation rate | 1 in model 2.3;<br>-0.02 in model 2.4 |
| $k_2$ (antibody-M)<br>(M = 10*(i-1) + j) | Forward binding constant between antibody-1 and virus | $Kon_i$ |
| $k_{-2}$(antibody-M) | Dissociation constant of antibody-1-virus complex | $Koff_i$ |
| $k_3$ | Feedback constant of antibody-virus complex on the regeneration of antibody | 5 |
| $k_4$ | Clearance rate of antibody-virus complex | 0.5 |
| $k_5$ | Antibody degradation rate | 0.01 |
| $k_6$ | Forward binding constant between antibody and self-antigen | $k_6*(k_7+k_4)/k_4/10^{16}$<br>($10^{16}$ = initial value of p) |
| $k_7$ | Dissociation constant of antibody-self-antigen complex | 10 |
| $k_{-7}$ | Feedback constant of antibody-self-antigen complex on the regeneration of antibody | 1 |
| $k_8$ | Clearance rate of antibody-self-antigen complex | 0.5 |
| $C_1$ | Replenish rate of self-antigen | $10^{13}$<br>($10^{13}$ = initial overall antibody concentration $*k_5$) |

Table S14：Parameter set in model 2.3 and model 2.4.

| Variables | Meaning | Initial value |
|---|---|---|
| z | virus | 100 in model 2.3; $10^{15}$ in model 2.4 |
| $Y_M$ (M = 10*(i-1) + j) | Antibody M | Prob_A(i)*Prob_B(j)*overall_initial_antibody*$\left(1-\frac{1}{\alpha}\right)+\frac{1}{\alpha}*$ $y_{M\_reshaped}$<br>(overall_initial_antibody = $10^{15}$; α represents mutation coefficient; $y_{M\_reshaped}$ represents reshaped level of antibody_M after wild strain infection. α equal to ∞ in primary infection case) |

| | | |
|---|---|---|
| $X_M$ | Antibody-M-virus complex | 0 |
| p | Self-antigen | $10^{16}$ |
| $q_M$ | Antibody-M-self-antigen complex | $Y_M*k_5/(k_1-k_4)$ |

Table S15：Time-dependent variables of model 2.3 and model 2.4.

**Calculation of ELISA results (Model 3)**

In the calculation of ELISA results, the tested viral antigen lacks self-replicating functionality, hence the value of k1 is 0. However, the initial concentration of the antigen is a large numerical value, which varies depending on the experimental conditions of the ELISA and the concentration level of the antigen used. Here, we set it as $10^{16}$. Moreover, this reaction only involves the binding and dissociation processes of antigen-antibody interactions. Ultimately, the desired outcome is the concentration of the total antigen-antibody complexes.

| Parameter Name | Description | Value |
|---|---|---|
| $k_1$ | Virus proliferation rate | 0 |
| $k_2$(antibody-M) (M = 10*(i-1) + j) | Forward binding constant between antibody-1 and virus | $Kon_i$ |
| $k_{-2}$(antibody-M) | Dissociation constant of antibody-1-virus complex | $Koff_i$ |

Table S16：Parameter set in model 3.

| Variables | Meaning | Initial value |
|---|---|---|
| z | Virus antigen | $10^{16}$ |
| $Y_M$ (M = 10*(i-1) + j) | Antibody M | $Y_{M\_reshaped}$ ($Y_{M\_reshaped}$ represents reshaped level of antibody_M at certain time point.) |
| $X_M$ | Antibody-M-virus complex | 0 |

Table S17：Time-dependent variables of model 3.

**Mathematical modeling of antibody atlas considering antibody isotype switching**

**(Model 4)**

In this model, we consider two different types of antibodies, namely IgM and IgG. They exhibit distinct distribution characteristics in terms of their binding affinity to antigens. Specifically, the ln(Kon) for IgG follows a normal distribution with parameters (-15.5, 0.4), while the corresponding ln(Kon) for IgM follows a normal distribution with parameters (-15.5, 0.8). Both IgG and IgM follow a normal distribution with parameters (1.5, 0.8) for ln(Koff). The model also incorporates the conversion coefficient, $k_9$, from IgM to IgG.

| Parameter Name | Description | Value |
|---|---|---|
| $k_1$ | Virus proliferation rate | 1 |
| $k_2$(IgG-M)(M = 10*(i-1) + j) | Forward binding constant between M$^{th}$ IgG and virus | IgG_Kon$_i$ |
| $k_{-2}$(IgG-M) | Dissociation constant of M$^{th}$ IgG -virus complex | IgG_Koff$_i$ |
| $k_2$(IgM-M)(M = 10*(i-1) + j) | Forward binding constant between M$^{th}$ IgM and virus | IgM_Kon$_i$ |
| $k_{-2}$(IgM-M) | Dissociation constant of M$^{th}$ IgM -virus complex | IgM_Koff$_i$ |
| $k_3$ | Feedback constant of antibody-virus complex on the regeneration of antibody | 5 |
| $k_4$ | Clearance rate of antibody-virus complex | 0.5 |
| $k_5$ | IgG degradation rate | 0.01 |
| $k_{5'}$ | IgM degradation rate | 0.02 |
| $k_6$ | Forward binding constant between antibody and self-antigen | $k_6*(k_7+k_4)/k_4/10^{16}$ ($10^{16}$ = initial value of self-antigen) |
| $k_7$ | Dissociation constant of antibody-self-antigen complex | 10 |
| $k_{-7}$ | Feedback constant of antibody-self-antigen complex on the regeneration of antibody | 1 |
| $k_8$ | Clearance rate of antibody-self-antigen complex | 0.5 |
| $k_9$ | The transformation coefficient from IgM to IgG | 0. |
| $C_1$ | Replenish rate of self-antigen | $10^{13}$ ($10^{13}$ = initial overall antibody concentration |

| | | |
|---|---|---|
| | | $*k_5)$ |

Table S18：Parameter set in model 4.

| Variables | Meaning | Initial value |
|---|---|---|
| z | virus | 100 |
| $y_M$ (M = 10*(i-1) + j) | $M^{th}$ IgG | Prob_A_IgG(i)*Prob_B_IgG (j)*overall_initial_IgG (overall_initial_IgG = $10^{15}$) |
| $R_M$ (M = 10*(i-1) + j) | $M^{th}$ IgM | Prob_A_IgM(i)*Prob_B_IgM(j)*overall_initial_IgM (overall_initial_IgM = $10^{14}$) |
| $x_M$ | $M^{th}$ IgG-virus complex | 0 |
| $S_M$ | $M^{th}$ IgM-virus complex | 0 |
| p | Self-antigen | $10^{16}$ |
| $q_M$ | $M^{th}$ IgG - self-antigen complex | $y_M*k_5/(k_1-k_4)$ |
| $V_M$ | $M^{th}$ IgM - self-antigen complex | $R_M*k_5/(k_1-k_4)$ |

Table S19：Time-dependent variables of model 4.

| Parameter Name | Value |
|---|---|
| Prob_A_IgG (1) | $P(X \leq -19.5)$ |
| Prob_A_IgG (2) | $P(-19.5< X \leq -18.5)$ |
| Prob_A_IgG (3) | $P(-18.5< X \leq -17.5)$ |
| Prob_A_IgG (4) | $P(-17.5< X \leq -16.5)$ |
| Prob_A_IgG (5) | $P(-16.5< X \leq -15.5)$ |

| | |
|---|---|
| Prob_A_IgG (6) | P(-15.5< X ≤ -14.5) |
| Prob_A_IgG (7) | P(-14.5< X ≤ -13.5) |
| Prob_A_IgG (8) | P(-13.5< X ≤ -12.5) |
| Prob_A_IgG (9) | P(-12.5< X ≤ -11.5) |
| Prob_A_IgG (10) | P(-11.5< X) |
| Prob_B_IgG (1) | P(x ≤ -2.5) |
| Prob_B_IgG (2) | P(-2.5< x ≤ -1.5) |
| Prob_B_IgG (3) | P(-1.5< x ≤ -0.5) |
| Prob_B_IgG (4) | P(-0.5< x ≤ 0.5) |
| Prob_B_IgG (5) | P(0.5< x ≤ 1.5) |
| Prob_B_IgG (6) | P(1.5< x ≤ 2.5) |
| Prob_B_IgG (7) | P(2.5< x ≤ 3.5) |
| Prob_B_IgG (8) | P(3.5< x ≤ 4.5) |
| Prob_B_IgG (9) | P(4.5< x ≤ 5.5) |
| Prob_B_IgG (10) | P(5.5< x) |
| Prob_A_IgM (1) | P(X' ≤ -19.5) |
| Prob_A_IgM (2) | P(-19.5< X' ≤ -18.5) |
| Prob_A_IgM (3) | P(-18.5< X' ≤ -17.5) |
| Prob_A_IgM (4) | P(-17.5< X' ≤ -16.5) |
| Prob_A_IgM (5) | P(-16.5< X' ≤ -15.5) |
| Prob_A_IgM (6) | P(-15.5< X' ≤ -14.5) |
| Prob_A_IgM (7) | P(-14.5< X' ≤ -13.5) |
| Prob_A_IgM (8) | P(-13.5< X' ≤ -12.5) |

| | |
|---|---|
| Prob_A_IgM (9) | P(-12.5< X' ≤ -11.5) |
| Prob_A_IgM (10) | P(-11.5< X') |
| Prob_B_IgM (1) | P(x' ≤ -2.5) |
| Prob_B_IgM (2) | P(-2.5< x' ≤ -1.5) |
| Prob_B_IgM (3) | P(-1.5< x' ≤ -0.5) |
| Prob_B_IgM (4) | P(-0.5< x' ≤ 0.5) |
| Prob_B_IgM (5) | P(0.5< x' ≤ 1.5) |
| Prob_B_IgM (6) | P(1.5< x' ≤ 2.5) |
| Prob_B_IgM (7) | P(2.5< x' ≤ 3.5) |
| Prob_B_IgM (8) | P(3.5< x' ≤ 4.5) |
| Prob_B_IgM (9) | P(4.5< x' ≤ 5.5) |
| Prob_B_IgM (10) | P(5.5< x') |

Table S20： Caculation of Prob_A and Prob_B. x follows normal distribution N(1.5,0.8), X follows normal distribution N(-15.5,0.4). x' follows normal distribution N(1.5,0.8), X' follows normal distribution N(-15.5,0.8).

**Modeling the impact of serum pretreatment on ELISA result**

To mitigate the occurrence of non-specific binding in the samples, various preprocessing methods are commonly employed, with one prevalent approach being the pretreatment of samples using bovine serum [2]. Our model simulates how sample pretreatment influences ELISA results.

We employed four concentration gradients of bovine serum, namely 0, $10^{15}$, $2 \times 10^{15}$, and $5 \times 10^{15}$. Antibodies present in bovine serum can interact with antibodies in the sample, effectively removing a portion of the antibodies before binding with the viral antigen. This significantly impacts the relative values of ELISA results for the samples. The outcomes are depicted in Figure S3.

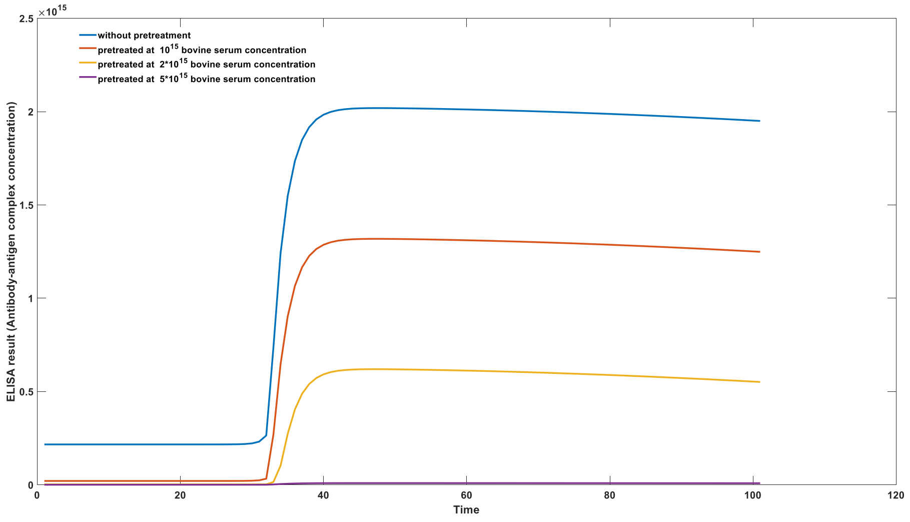


**Figure S3: Modeling the impact of serum pretreatment on ELISA result**

From Figure S3, it is evident that untreated samples exhibit higher initial ELISA values. While the absolute ELISA values post-viral infection are also elevated, the relative increment is limited, leading to decreased detection sensitivity. Treatment of samples with an appropriate concentration of bovine serum reduces both the absolute ELISA values post-viral infection and the initial ELISA values significantly, thereby enhancing detection sensitivity. Excessive concentrations of bovine serum treatment result in all ELISA values decreasing to very low levels. However, it should be noted that sample pretreatment does not effectively eliminate non-specific binding. Additionally, different experimenters often employ varied pretreatment methods and concentrations, resulting in ELISA measurements being relative to the initial state. The same sample may yield vastly different ELISA relative values after different pretreatments. Thus, solely relying on ELISA results cannot determine changes in specific antibodies accurately, as this change represents a shift in the overall antibody atlas of the host, rather than the proliferation of individual or a few specific antibodies. For instance, while the blue solid line may suggest a several-fold increase in neutralizing antibodies, the red solid line may indicate a hundred-fold increase in specific antibodies.
